# CellWalker2: multi-omic discovery of hierarchical cell type relationships and their associations with genomic annotations

**DOI:** 10.1101/2024.05.17.594770

**Authors:** Zhirui Hu, Pawel F. Przytycki, Katherine S. Pollard

## Abstract

CellWalker2 is a graph diffusion-based method for single-cell genomics data integration. It extends the CellWalker model by incorporating hierarchical relationships between cell types, providing estimates of statistical significance, and adding data structures for analyzing multi-omics data so that gene expression and open chromatin can be jointly modeled. Our open-source software enables users to annotate cells using existing ontologies and to probabilistically match cell types between two or more contexts, including across species. CellWalker2 can also map genomic regions to cell ontologies, enabling precise annotation of elements derived from bulk data, such as enhancers, genetic variants, and sequence motifs. Through simulation studies, we show that CellWalker2 performs better than existing methods in cell type annotation and mapping. We then use data from the brain and immune system to demonstrate CellWalker2’s ability to discover cell type-specific regulatory programs and both conserved and divergent cell type relationships in complex tissues.

## Introduction

Single-cell technologies are improving our understanding of the cellular composition of complex tissues. They increase the resolution at which it is possible to study cell types, defined here as distinct cell lineages as well as related cells with varying functions or differing states. This knowledge is propelling new discoveries about cellular diversity across evolution, development, and disease. In order to achieve this, one of the fundamental computational tasks is to label cells from either single-cell RNA sequencing (scRNA-Seq) or single-cell ATAC sequencing (scATAC-Seq). Many downstream analyses, such as identifying differentially expressed genes or differentially accessible regions, rely on accurately annotating cells to different cell types. Many methods have been developed for cell labeling. Some of them are based on classical machine learning methods (e.g. Seurat[1], Signac[2], ArchR[3], CellTypist[4], cisTopic[5], snapATAC[6], and LIGER[7]), while others leverage deep learning (e.g. MARS[8], scArches[9], and scANVI[10]). Beyond discrete cell types, methods such as velocyto[11] and CellRank[12] infer cell trajectories or cell fates using RNA velocity, while MIRA[13] does so using expression and accessibility.

As more and more single-cell data in similar tissues or conditions is generated from different research groups, it has become important to be able to compare cell type annotations across studies. Single-cell datasets from the same tissue often have distinct cell type labels due to biological variation across samples, different modalities of the data (e.g. scRNA-Seq, scATAC-Seq or multiome data), variable sequencing depths, different computational methods or tuning parameters (e.g., clustering resolution), and divergent choices when naming cell clusters. Mapping and comparison between the cell type labels are imperative for integrating single-cell data from different groups and comparing the cell type-specific biological findings from each study. Although cell type labels can be manually compared after annotating cells using both sets of labels, few computational methods can automatically match the cell type labels and provide a probabilistic measure of the mapping. Moreover, cell types usually have hierarchical structures. Cell type labels from different datasets might have different resolutions and they may agree only upon certain levels of the hierarchy. Correctly placing a cell type on a cell-type hierarchy can facilitate meaningful comparisons between similar cell types in different samples or conditions. However, most existing methods do not take into account the hierarchical structure inherent in cell-type labels, potentially leading to failures in capturing correspondences across different levels. MARS, treeArches[14] and CellHint[15] can directly map cell type labels. MARS trains a neural network model for cell type classification, treeArches uses kNN classifiers after embedding cells via a deep learning model, and CellHint develops a predictive clustering tree algorithm to compare cell types. The latter two build cell-type hierarchies upon integrating multiple datasets but cannot compare existing cell-type hierarchies directly.

Integrating data from different omics modalities can also facilitate interpretation of various annotations from bulk data at the single-cell level. For example, scATAC-Seq data can be used to assess activities of regulatory regions predicted from transcription factor (TF) motifs or identified from bulk ChIP-Seq or ATAC-Seq data[16, 17, 18, 19]. Similarly, single nucleotide polymorphisms (SNPs) from expression quantitative trait locus (eQTL) experiments or genome-wide association studies (GWAS) can be attributed to particular cell types using scATAC-seq, which aids in elucidating the molecular mechanism of these genetic variants [20]. Most methods (e.g. Signac, snapATAC, ArchR) identify cell type-specific annotations as a downstream task: after clustering, annotating cells and identifying differentially accessible regions (DARs), they usually test for the enrichment of bulk-derived annotations in cell type-specific peaks. However, every step loses some information from the original sequencing data of each cell, and cell-type labeling and DAR identification can have large uncertainties especially in complex tissues (e.g. brain), thereby complicating the calculation of statistical significance for associations between annotations and cell types. cisTopic uses a topic model to simultaneously cluster cells and regions, which enables users to identify TF motifs for each topic, but these are not directly linked to cell types. In contrast, CellWalker[19] provides a framework that directly assigns bulk-derived labels to cell types by constructing a graph of cells and cell types using scATAC-Seq data. While this enables regions to be mapped to cell types, the statistical significance of these mappings is not established, which makes it difficult to compare cell type-specific annotations across different conditions or species. In addition, although using a cell type hierarchy can increase the power to detect cell type-specific regions (e.g. eQTLs [21]), most existing methods can only map bulk-derived annotations to a single level of a cell type hierarchy.

Motivated by these gaps in the single-cell toolkit, we sought to combine the individual strengths of existing methods into a single integrative modeling framework while ensuring that the resulting method provides robust performance and estimates of statistical significance. Our approach was to significantly extend the CellWalker graph-diffusion model to add 1) hierarchical relationships between cell types, 2) a permutation null distribution for estimating statistical significance, 3) flexibility to use scRNA-seq, scATAC-seq, or multiome data, and 4) functionality for comparing cell types across contexts. Our open-source software, CellWalker2, can be used to assign cell type labels to either genome coordinates (binding sites, SNPs, or other elements) or cells from single-cell experiments using any cell type ontology (hierarchical or not). The model also enables statistical comparisons between cell types from two or more ontologies, allowing users to assess the similarity of cell types across species, disease states, and research groups.

We first describe the design of CellWalker2 and compare its performance to existing methods for specific analysis tasks using simulations and existing biological knowledge. We then apply CellWalker2 to single-cell data from human peripheral blood mononuclear cells (PBMCs) and brain cortex samples from multiple developmental stages and species. These investigations demonstrate the benefits of using hierarchical cell type ontologies and assessing statistical significance to overcome biases of current methods, such as attraction to abundant cell types, low sensitivity for intermediate cell types or broad cell lineages, and inability to infer when no confident cell type mapping exists. We conclude that CellWalker2’s unified graph-based framework is a robust model for single-cell data integration.

## Design

CellWalker2 serves as a modeling and statistical inference tool used after processing raw sequencing reads. It takes count matrices as inputs (Figure 1A), specifically gene-by-cell and/or peak-by-cell matrices from scRNA-Seq and scATAC-Seq respectively, allowing it to naturally plug in downstream of existing single cell processing software, e.g. Seurat and Signac. CellWalker2 builds a graph (Figure 1B) that integrates these inputs, plus a cell type ontology (e.g., tree of cell type relationships with marker genes for each leaf node) and optionally genome coordinates for regions of interest (e.g., genetic variants, regulatory elements). The resulting graph is heterogeneous, with nodes representing cells, cell types (“labels”), and (if provided) genome coordinates (“annotations”). CellWalker2 can be applied to any number of label sets with little modification. The algorithm then conducts a random walk with restarts on this graph and computes an influence matrix. From sub-blocks of the influence matrix, CellWalker2 learns relationships between different nodes. For instance, label-to-label similarities enable users to compare different cell type ontologies by learning how cell types in one context (e.g., lab, disease state, species) map to cell types in another. With hierarchical ontologies, CellWalker2 provides relationships not only for cell types with marker genes, but also for internal nodes that represent broader cell types. As additional examples, cells can be mapped to cell types using cell-to-label similarities, and bulk-derived genomic elements and genetic variants can be mapped to cell types using annotation-to-label similarities. Finally, CellWalker2 performs permutations to estimate the statistical significance (Z-scores) of these learned associations (Figure 1C).

**Figure 1:**
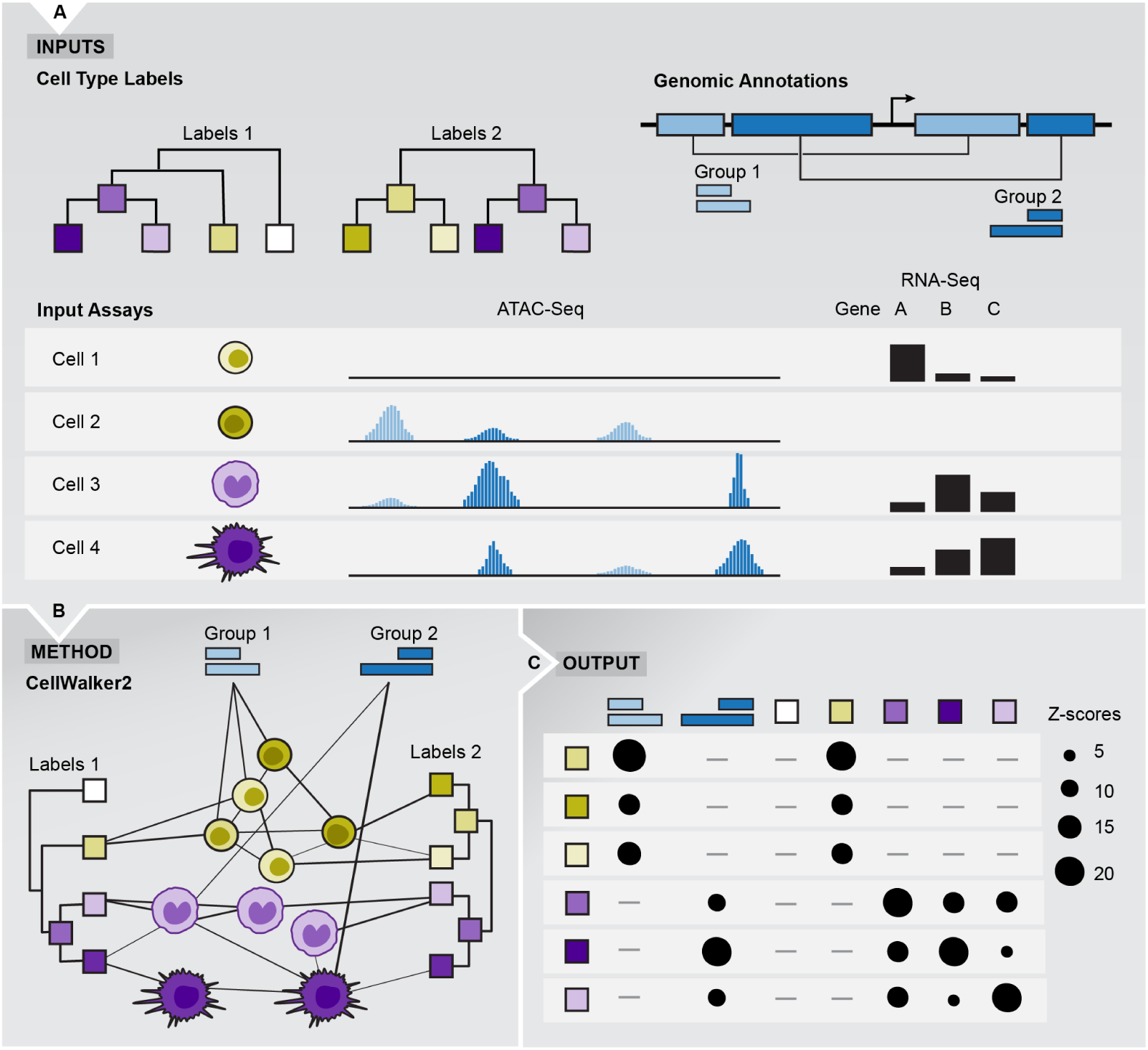
Overview of CellWalker2. (A) The inputs to CellWalker2 are 1) one or more sets of cell type labels with marker genes and an optional hierarchical structure; 2) cells with RNA-Seq and/or ATAC-Seq data; and 3) optionally, gene sets or annotations with genome coordinates that may be derived from bulk assays (e.g., candidate regulatory elements and genetic variants). (B) CellWalker2 constructs a graph with labels, cells, and annotations as nodes. Cells are connected to each other, to labels, and to annotations with edge weights that are computed based on the available assays for the cell. Cell type labels connect to cells with RNA-seq based on expression of their marker genes, gene sets connect to cells also through RNA-Seq, and annotations connect to cells with ATAC-seq based on chromatin accessibility of their genome coordinates. A random walk on the graph is performed to calculate the influence scores between all pairs of nodes. The normalized influence score from each cell to cell type labels is used to annotate cells. (C) CellWalker2 outputs Z-scores that measure the statistical associations between each cell type label and 1) every annotation and 2) all other cell types. This general framework is flexible and can be modified for different applications by generating graphs with different combinations of assays, labels, and annotations (Figure S2).

These new features distinguish CellWalker2 from the CellWalker method, which did not model hierarchical relationships or assess statistical significance, used only open chromatin data and not gene expression to quantify similarity, and employed an *ad hoc* method to map genome coordinates to labels rather than including coordinates as nodes in the graph. In the following, we highlight these new functionalities by first describing the CellWalker2 model and then demonstrating: 1) cell annotation using scRNA-seq data, 2) comparing cell-type hierarchies using scRNA-Seq data, and 3) mapping bulk-derived regulatory regions to cell types using multiomic data. For each application, we describe the required inputs to CellWalker2, the graph model, and the inference that is performed using the resulting influence matrix.

### Constructing cell graphs

CellWalker2 constructs a graph with three types of nodes: cells, labels, and annotations (Figure 1B; STAR Methods). Building a heterogeneous graph allows CellWalker2 to compute the statistical significance of mappings between different node types in a coherent manner. Cell nodes may possess scATAC-Seq, scRNA-seq, or multiomic data. This data is used to compute cell-to-cell edges using commonly employed similarity metrics and a K Nearest Neighbors algorithm. It is also used to compute cell-to-label edges based on the expression (or accessibility) of each label’s marker genes in each cell and annotation-to-cell edges based on accessibility of the genome regions in each cell. Hierarchical label-to-label edges are an input to CellWalker2 that is either part of the cell type ontology or pre-computed using marker gene or global expression similarity.

### Computing influence scores and Z-scores

CellWalker2 does a random walk with restarts on the graph and computes the influence matrix, which is the transition probability at the steady state of the random walk (STAR Methods). This matrix summarizes how strongly each node in the graph is associated with all other nodes given the graph topology and edge weights, and it can be used to derive relationships amongst labels from two or more ontologies in the context of the cells, from labels to cells or annotations, and amongst cells. To quantify the statistical significance of relationships in the influence matrix, CellWalker2 computes a Z-score for each entry by comparing the observed value with its expectation and variance under a permutation null distribution. The way this distribution is estimated depends on the type of query (STAR Methods) but generally involves permuting graph edges while maintaining node degree. Controlling for node degree is critical, because labels that are prevalent and connected to many cells tend to have larger influence scores just by chance. In contrast, our Z-scores are robust to variability in node degrees and quantify the statistical significance of node associations (Figure S1).

### Cell annotation

To annotate cells, the cell graph contains cell type labels and cells with gene expression data from the query dataset (Figure S2A) or both query and reference dataset (Figure S2B). Cell type labels contain marker genes obtained from the reference dataset. Then, CellWalker2 computes the cell-to-label influence scores and normalizes these by the total influences of each label. It assigns each cell to the cell type label with the largest normalized influence score, and the vector of normalized scores across all labels can be used as a probabilistic assignment.

To benchmark CellWalker2, we designed a series of simulation scenarios where cell labels are known (STAR Methods, Figure S3a), ran CellWalker2 and the commonly employed tool Seurat ignoring the known labels, and compared their performance. In the easy scenario, the two datasets (reference and query) are simulated in the same way without batch effects or dropouts. In the medium scenario, we added batch effects between the two dataset as well as dropouts, with more dropouts in the query dataset (Figure S3b). We further probed the role of cell label prevalence by altering the number of cells in the reference dataset. In the hard scenario, we reduced the distance between all cell types in both reference and query datasets further (Figure S3c). The last two scenarios are more challenging but more realistic for scRNA-Seq studies.

Both CellWalker2 and Seurat perform equally well in the easy scenario (Figure 2A). With batch effects and dropouts, CellWalker2 using only cells from the query dataset to construct the cell graph performs better than Seurat (Figure 2A). Only including cells from query dataset has the computational advantage when the reference dataset has many cells (e.g. annotating cells to a cell atlas), while Seurat needs to integrate cells from both datasets. After adding cells from the reference dataset to the graph, CellWalker2 performs even better (Figure 2A). We hypothesize that this is due to the reference dataset having less noise, because adding an equal number of cells from the query dataset does not change CellWalker2’s performance. When we perturbed cell type composition by decreasing the size of cell type 4, we observed that Seurat annotates more cells from this label to the related and more abundant cell type, while CellWalker2 is robust to the change (Figure S4). Finally, if the cell types in the reference dataset are also not well separated, using a hierarchical cell type ontology further improves the performance of CellWalker2 to some degree. We conclude that hierarchical ontologies and leveraging information from cells in the reference dataset that complements the marker gene data at label nodes both help CellWalker2 to outperform Seurat.

**Figure 2:**
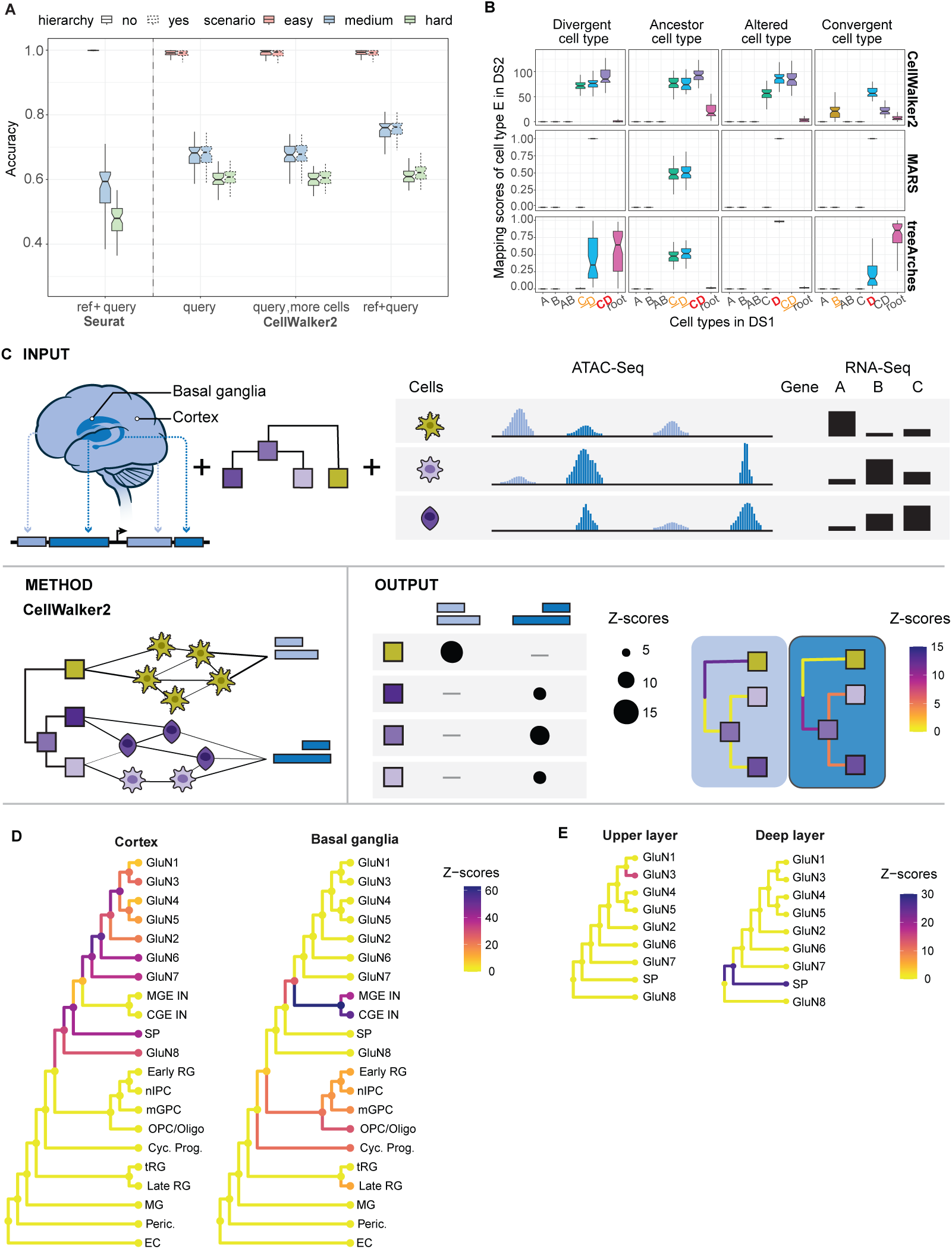
Examples show CellWalker2’s functionalities: cell annotation, cell type mapping, and labeling bulk-derived annotations with cell types. (A) For cell annotation, CellWalker2 outperforms Seurat in simulations. Colors represent different simulation scenarios from easy (no batch effect or dropout) to hard (batch effect and more dropout). “Hierarchy” indicates whether edges between cell types are included. For CellWalker2, different alternatives are implemented: “query” only includes cells from the query dataset to construct the graph, “query, more cells” only includes cells from the query dataset but increases the number of cells to be equal to the total number of cells in the query and reference datasets, and “ref+query” includes cells from both the reference and query datasets in the cell graph. (B) For cell type mapping between simulated dataset 1 and 2 (DS1 and DS2), CellWalker2 outperforms treeArches and MARS. Boxplots show mapping cell type E of DS2 to different cell types in DS1 in four simulation scenarios (columns). The scores on the vertical axes are Z-scores for CellWalker2 and probabilities for MARS and treeArches. In each panel, the horizontal axis represents different cell types in DS1. AB: parent node of A and B; CD: parent of C and D. CellWalker2 can map to all nodes in the cell type hierarchy, treeArches to tip and root nodes but not internal nodes, and MARS only to tips. The closest cell type of E (i.e., the expected output given the cell type relationships) is in red and bold, while the second closest cell types are in orange and underlined (See Figure S5 for details on cell type relationships). (C) Illustration of using CellWalker2 to map regulatory elements (pREs) from different brain regions to cell type labels. Colors of rectangles represent two groups of regulatory elements. Each track is the ATAC-Seq profile of a single cell with RNA-Seq for three genes shown as barplots on the right. The first cell is from a cell type in which light blue elements have higher chromatin accessibility, while the last two are from two related cell types in which dark blue elements have higher accessibility. (D) and (E) pREs in different brain regions of human developing telencephalon [23] show differential chromatin accessibility. (D) Z-scores of mapping basal ganglia versus cortex specific pREs to the cell type hierarchy in Trevino et al.. (E) Z-scores of mapping upper versus deep layer specific pREs from prefrontal cortex onto the subtree of excitatory neurons. Nodes and branches of the cell type tree are colored by Z-scores.

### Mapping cell type hierarchies

CellWalker2 maps cell types from two or more datasets using a graph that includes nodes for all cells from the combined datasets (each with scRNA-seq data) and for the labels (each with marker genes) from each (potentially hierarchical) cell type ontology. Edges are computed as for a single dataset, including edges between cells from different datasets (Figure S2C). The influence matrix includes scores for all pairs of labels from different ontologies. These are converted to Z-scores using a null distribution obtained by permuting edges between cells and labels, as described above, providing the statistical significance of the mapping of cell types between datasets.

We performed simulations to compare CellWalker2 with treeArches, which can add cell types in a new scRNA-Seq dataset to an existing cell type ontology, and MARS, which can provide probabilistic naming of a cell cluster in the new scRNA-Seq dataset. We simulated four scenarios using Splatter (Figure S5) for mapping four cell types in dataset 2 (DS2) to those in dataset 1 (DS1). Cell types A,B,C are shared between datasets but cell type E in DS2 is added to the cell type hierarchy of DS1 in different ways (STAR Methods). In all cases, CellWalker2 assigns the largest Z-score to the cell type closest to cell type E, and also discovers cell types similar to cell type E, with Z-score decreasing as the similarity diminishes (Figure 2B; Figure S6). On the other hand, MARS and treeArches failed to recognize the similarity to B in the Convergent cell type scenario. MARS maps E to D with probability 1, whereas treeArches assigns large probabilities to root, which means that it either treats E as an entirely new cell type or is unable to locate E, as it partially shares features with multiple cell types. In this case, we also showed that the Z-score mapping cell type E to B gradually increases as we increase the similarity between cell type E to cell type B (Figure S7; Note S1). Moreover, Z-scores from CellWalker2 effectively differentiate between various cases. In comparison, MARS only managed to distinguish the Ancestor cell type case from others, while treeArches struggled to differentiate between the Divergent and Convergent cell type cases.

To see how the number of cells affects the mapping results, we varied the number the cells of cell type D in the ‘Divergent cell type’ case, where cell type E in DS2 has equal distance to cell type C and D in DS1. treeArches’s results are dominated by the major cell types, as it maps cell type E to cell type C with probability approaching 1 when the number of D cells decreases and to C and D with roughly equal probability when cell types C and D have equal number of cells. However, CellWalker2’s results remain similar despite varying in the number of cells in which it maps cell type E to the parent of C and D with the largest Z-scores then to C and D with similar Z-scores (Figure S8). Thus, CellWalker2’s cell type mapping results are less influenced by prevalent cell types.

### Cell type labeling for bulk-derived annotations

For assigning cell type labels to annotations with genome coordinates, the CellWalker2 graph includes nodes for labels, for annotations, and for cells with scATAC-seq and/or scRNA-seq. Some cells must have multiome data in order to connect nodes for annotations to the graph via their chromatin accessibility and to connect nodes for labels via expression of their marker genes (Figure S2D). But the graph may also include cells with only scRNA-seq or only scATAC-seq, which provide additional information about cell types (Figure S2E). Cells with multiomic data serve as bridges connecting cells with only one modality and creating edge paths between regions and cell type labels; a similar idea is exploited in [22]. Genome coordinates can represent any elements of interest, typically bulk-derived annotations, such as gene sets, enhancers, genetic variants, or motifs within open chromatin regions.

We validated this functionality using 19,151 predicted regulatory elements (pREs) identified from bulk ATAC-seq of different anatomical regions of the human developing cortex [23]. As the cell type composition of different regions of the cortex is known, we could use this information to benchmark CellWalker2’s label-to-annotation mappings versus commonly employed statistical tests as well as the original CellWalker method. To construct the cell graph, we utilized a human developing cortex dataset which contains scRNA-seq and scATAC-seq at different developmental stages as well as 10X multiomic data at post-conception week (pcw) 21 [24] (Figure 2C). We first constructed a cell graph with scRNA-seq and multiomic data. First, we used CellWalker2 to label the cell types of 6,941 pREs from cortical plate samples and 3,463 from basal ganglia. Cortical plate pREs received larger Z-scores for excitatory neurons, while basal ganglia pREs had higher Z-scores for inhibitory neurons as well as progenitors and radial glia (Figure 2D), consistent with expectations. Wilcoxon tests comparing the distribution of edge weights to basal ganglia versus cortical plate pREs for each cell type defined in [24] shows a similar cell type specificity pattern as CellWalker2 (Figure S9b), but this method requires a predefined set of cell types and cannot integrate cells from one study with an ontology from another source. As a third method, we first identified differentially accessible regions (DARs) using psuedobulked scATAC-seq for each cell type and then tested for enrichment of cortical plate and basal ganglia pREs overlapping these DARs with Fisher’s exact test. We detected enrichment of DARs for cycling progenitors and radial glia in cortex-specific pREs whereas these cell types do not enter the cortical plate in early development (Figure S9c). In contrast to the other methods, the original CellWalker does not assign distinct cell types to pREs from cortex versus basal ganglia (Figure S9a), suggesting that using a cell type hierarchy and modeling genome regions as graph nodes notably boost the performance of CellWalker2.

To evaluate CellWalker2 on smaller sets of pREs in a context where cell type differences are more subtle than cortex versus basal ganglia, we zoomed in on excitatory neurons of the dorsolateral prefrontal cortex (PFC) and repeated the above analysis using 2,333 pREs from the upper cortical layers versus 445 from deep layers. Consistent with expectations, CellWalker2 Z-scores map upper layer pREs to glutameteric neurons (GluN3) and deep layer pREs to subplate (SP) (Figure 2E). Adding scATAC-seq cells to the graph revealed significant mappings between upper layer pREs and additional subtypes of glutameteric neurons (e.g. GluN5) (Figure S10a), and these mappings varied as expected when using scATAC-seq from an earlier developmental stage (pcw 16) or a more fine-grained cell type ontology (Figure S10b and S10c). In summary, given a set of genome coordinates, CellWalker2 can identify the particular cell types in which the chromatin in these regions is most accessible using single cell multiomic data and/or scATAC-Seq data.

### Software

We implemented CellWalker2 in R by extending the CellWalkR package[25]. The open-source software, available at Github https://github.com/PFPrzytycki/CellWalkR/tree/dev, includes documentation and vignettes. CellWalker2’s functions and pipelines are shown in Figure S11.

## Results

To demonstrate how the robustness and flexible functionality of CellWalker2 enables biological discovery, we analyzed single-cell data from three contexts encompassing different complex tissues, developmental stages, and species. We also compared our findings to results from other methods.

### Human peripheral blood mononuclear cells

PBMCs are heterogeneous and contain many closely related cell types, exemplified by various kinds of T cells and immune cells that transition into alternative states upon stimulation. In this dynamic landscape, TFs emerge as central regulators, governing gene expression and cellular functions within PBMCs. Consensus on cell type definitions across studies is lacking, as is a comprehensive list of activating TFs in cell types or lineages. Here, we demonstrate that CellWalker2 can map cell types to multiple levels of a cell type hierarchy, discern distinct cell states and identify transcription factors specific to particular cell types or lineages.

#### Mapping cell type hierarchies by scRNA-Seq data

We analyzed two human PBMC datasets [26, 27] in which different strategies were used to define cell types at a high resolution: marker gene expression [26] versus cellular functions [27], complicating direct comparisons between the two cell type ontologies.

Before deeply exploring results from CellWalker2, we compared its cell type mappings to those of MARS and treeArches in the context of existing knowledge about PBMC biology. MARS fails to map 38% of cell types, including known correspondences between platelets and megakaryocytes and between plasmacytoid dendritic cells (pDC and DC c4-LILRA4), which are correctly mapped by CellWalker2 and treeArches. Furthermore, only 55% of MARS’s top mappings overlap with the top three CellWalker2 mappings, mostly due to MARS making errors (Figure S12), whereas 82% of treeArches’s top-mapped cell types overlap with the top three CellWalker2 mappings, and these top Z-scores usually have similar magnitude. Some differences between CellWalker2 and treeArches appear to stem from treeArches being biased towards more prevalent cell types, as we saw in simulations. For example, treeArches maps CD4 cytotoxic T lymphocytes to more prevalent CD8 cytotoxic T lymphocytes (Figure S12). TreeArches also misses the correspondence between plasma cells and B c05-MZB1-XBP1, where *XBP1* is a marker of plasma cells [28]. CellWalker2 identifies both of these biologically supported mappings, and it also performs well on cell types with multiple markers, such as associating Mono c4-CD14-CD16 with both CD14 and CD16 monocytes (Figure S12). Supporting these findings, CellWalker2’s Z-scores are correlated with the proportion of overlapping markers (Figure S13), with higher marker concordance than treeArches (Figure 3A).

**Figure 3:**
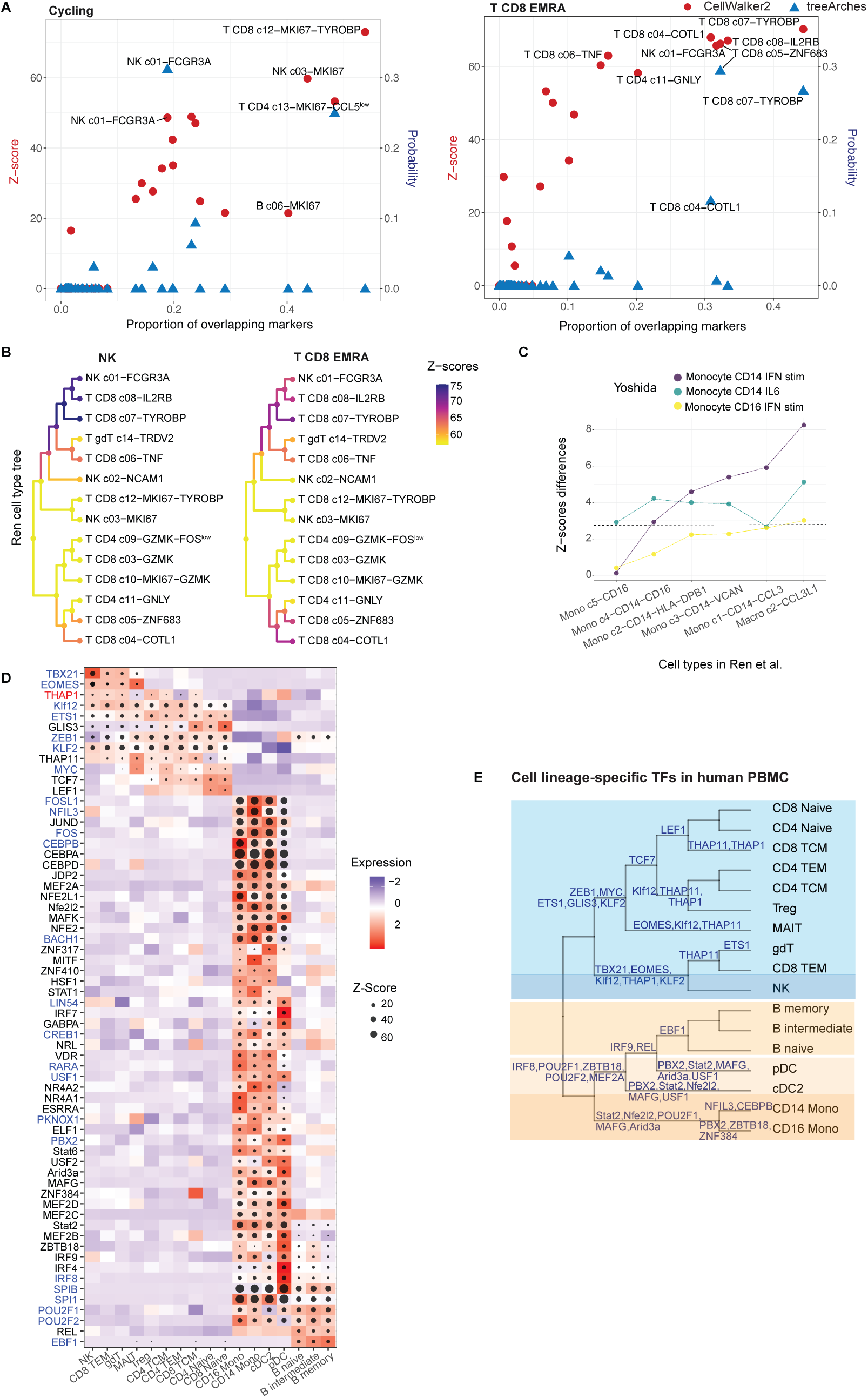
Mapping cell types and identifying cell type-specific TFs in human PBMC using CellWalker2. (A) Z-scores are correlated with the proportion of overlapping positive markers between cell types. Left and right: mapping scores of cycling and T CD8 EMRA cell type to other cell types in Ren et al.. For each plot, the left Y-axis (red) is Z-score by CellWalker2 and the right (blue) is probability by treeArches; X-axis, the proportion of overlapping positive markers. (B) CellWalker2 maps NK to a single clade and T CD8 EMRA to multiple clades on the cell type tree in Ren et al.. The branches and nodes are colored by Z-scores. (C) Z-scores can reflect the difference between a stimulated cell type and its unstimulated counterpart. Y-axis shows differences in Z-scores between mapping stimulated and unstimulated monocytes from Yoshida et al. to monocyte related cell types in Ren et al. (X-axis), see STAR Methods. Dashed line: Bonferroni-adjusted P-value <0.05. (D) and (E) CellWalker2 identifies cell type-specific TFs. (D) Each row is a TF and each column is a cell type. The size of the dot represents Z-score. The color of each square is the standardized gene expression of a TF in a particular cell type. TF names colored red are universal stripe factors and blue are other stripe factors [34]. (E) Tips and internal nodes of the cell type tree for selected TFs. The top 5 TFs (ranked by the correlation between Z-score and expression level) are shown for each node. Shaded regions on the cell type tree reflect different classes of cell types (see STAR Methods).

One feature of CellWalker2 is that it usually maps labels in one dataset to multiple hierarchically related labels in the other dataset. Z-scores generally cluster cell types into 4 groups corresponding to different subtrees of the PBMC ontology: plasma and B cells, monocytes, NK and cytotoxic T cells, and other T cells (Figure S12). Instead of trying to pinpoint a single tip within each clade, which could be unrealistic due to low signal-to-noise ratio or real biological differences, CellWalker2 is able to map to multiple related labels and their ancestral nodes (e.g., NK in Figure 3B). Z-scores also enable CellWalker2 to quantify similarity to multiple labels from different parts of the cell type tree (e.g., T CD8 EMRA in Figure 3B and cycling cells in Figure S12). The magnitude of Z-scores reflect the relative significance of hits for these multi-mapping labels. When there is a strong 1:1 mapping, CellWalker2’s Z-score will be highest for that leaf node (e.g., platelets and pDCs), whereas an ancestral node representing a broad cell type will score higher when there is more ambiguity. Furthermore, Z-scores capture differences between stimulated cell types and their unstimulated counterparts (e.g., baseline versus IFN-stimulated monocytes and NK cells in Figure 3C), whereas treeArches assigns these cell types with similar probabilities. We see this flexibility as an advantage, but users should be aware that it decreases the chance of seeing one highly specific cell type mapping for each label in the query dataset.

#### Cell type-specific transcription factors by multiomic data

We next applied CellWalker2 to human PBMC 10X multiomic data with the goal of identifying cell type-specific transcription factors (TFs). We built a cell graph in which TF motif instances in the genome are annotation nodes and then computed Z-scores for their annotation-to-label mappings. We analyzed these results alongside TFs discovered using ArchR, Signac, and SCENIC+. ArchR and Signac are examples of traditional TF motif enrichment analyses that call DARs and then treat them equally. In contrast, CellWalker2 assigns higher Z-scores to TFs whose motifs are in regions connected to many cells in a given cell type. This may enhance CellWalker2’s ability to identify cell type-specific TFs, given that DARs exhibit varying degrees of cell type specificity, particularly when dealing with cell types at a finer resolution and calling DARs from sparse scATAC-Seq data. SCENIC+ identifies cell-type specific regulatory modules by combining information about TF expression, motif accessibility, and target gene expression. This motivated us to filter TFs identified by CellWalker2 by removing those with poor correlation between expression levels and Z-scores or low expression in the cell types with high Z-scores, though we did not use target genes.

CellWalker2 identifies many cell type-specific TFs including well-known regulators, such as *TBX21* and *EOMES* in CD8 TEM and NK cells, *LEF1* for T cells, *TCF7* in T cells and *EBF1* in B cells (Figure 3D). In contrast, some of these regulators of specific lineages show broad cell type enrichment patterns when analyzed by Signac or ArchR (e.g., *TBX21*, *EOMES* and *EBF1*) (Figure S14). Additionally, Signac p-values are less correlated with TF expression than are CellWalker2 Z-scores and ArchR p-values (Figure S14a). Many well-known cell type-specific TFs are identified by both CellWalker2 and SCENIC+ (Figure S15). However, the scores from SCENIC+ better distinguish TFs with similar motifs, such as *SPI1* and *SPIB* having higher scores in monocytes and B cells, respectively (Figure S14). On the other hand, some TFs have very low expression in at least one of the cell types identified by SCENIC+ (e.g., *ETS1*, *KLF2* and *LEF1* in B cells) (Figure S14c), suggesting these could be false positives or TFs that function at very low expression levels.

A distinctive feature of CellWalker2 is its ability to place TFs on the cell type hierarchy, going beyond mappings to individual cell types (Figure 3E). Ancestral nodes receive high Z-scores when TFs play regulatory roles in multiple related subtypes or the resolution from scATAC-Seq data is insufficient to distinguish between related cell types despite their showing distinct marker gene expression in the data used to build the cell type ontology. For example, CellWalker2 identifies *EBF1* as an active regulator for the ancestral node of all B cells, *POU2F1* in B cells, DCs and monocytes, *LEF1* and *TCF7* in different clades of T cells, and *TBX21* and *EOMES* in the ancestor of NK and effector memory T cell types. In other cases, the activity of TFs does not fully correspond to the hierarchical structure of the cell type ontology. For instance, *THAP11* and *THAP1* are significant in multiple types of T cells and NK cells. In summary, for each TF, CellWalker2 can discover cell types or subgroups of cell types on the cell type tree in which it may have putative regulatory functions. On the other hand, CellWalker2 can identify groups of TFs that might collectively have regulatory functions in groups of similar cell types.

### Human developing cortex

The developing cortex is complex organ with numerous interrelated cell types. Cell type labels often vary across different studies, impeding comparisons and integrative analyses. Here, we use two independently collected human developing cortex scRNA-Seq datasets [29, 30] having different cell type classification criteria (STAR Methods) to illustrate this challenge. We hypothesized that CellWalker2 could overcome this challenge using the ability of its cell graph to capture cross-study relationships and generate probabilistic mappings between nodes.

#### Annotating cells by scRNA-Seq data

We annotated cells in the Polioudakis et al. dataset (query) using the cell types defined in Nowakowski et al. (reference) using CellWalker2 and Seurat, and compared the cell annotations with the original cell type labels in the Polioudakis et al. dataset. As Seurat cannot use a hierarchical cell type ontology, we first ran CellWalker2 without considering cell type relationships. Although the top-mapped cell types are mostly consistent between methods (Figure 4A), CellWalker2 additionally provides probabilistic mappings to related cell types (Figure S16a).

**Figure 4:**
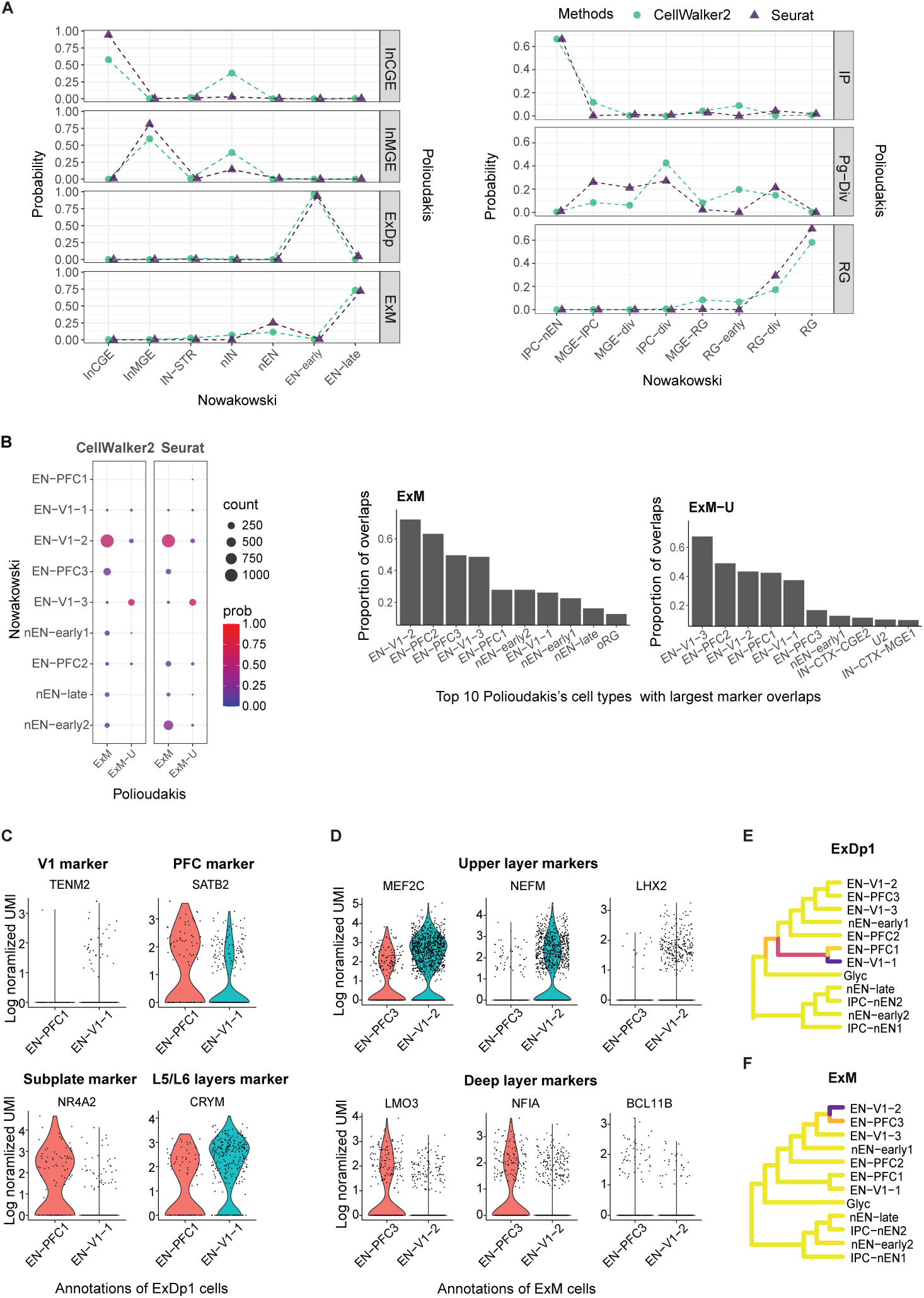
Annotating cells with a reference cell type hierarchy using human developing cortex scRNA-Seq data. (A) Probabilities (Y-axis) of annotating cells (rows) in Polioudakis et al.’s dataset by the cell types from Nowakowski et al. (X-axis) using CellWalker2 (emerald circles) versus Seurat (violet triangles). Similar cell types are grouped together (Table S1 and S2). Left: cell types are arranged from inhibitory to excitatory neurons; Right: cell types are arranged from intermediate progenitors (IP) to radial glias (RG). (B) CellWalker2’s cell annotations show expected differences across subtypes of maturing excitatory neurons and high proportions of marker gene overlap between query cells and the annotated cell types. Left: Heatmap showing annotations for ExM and ExM-U cells. The color and size of each dot presents the probability and the number of cells mapped (at least 5 cells are shown). Right: Barplots showing the proportion of positive markers of ExM ane ExM-U that overlap with positive markers of cell types in Nowakowski et al.. Top 10 cell types with largest marker overlaps are shown. (C) ExDp1 cells annotated to either EN-PFC1 or EN-V1-1 by CellWalker2 show differential expression of layer or region specific marker genes. (D) ExM cells annotated to either EN-PFC3 or EN-V1-2 by CellWalker2 show differential expression of upper or deep layer specific genes. (E) and (F) CellWalker2 annotates ExDp1 and ExM cells in the Polioudakis et al.’s dataset by the cell type hierarchy from Nowakowski et al.. Colors of the branches represent the percentage of cells annotated to each cell type on the tree. Explanation of abbreviations in Table S1 and S2.

We used excitatory neurons to further explore the challenges of cross-study cell annotation because both datasets have multiple subtypes of excitatory neurons from different layers, developmental stages or areas. The Polioudakis et al. dataset includes five types of excitatory neurons: two deep layer subtypes (ExDp1 and ExDp2), maturing (ExM), upper layer enriched maturing (ExM-U), and migrating (ExN). The Nowakowski et al. dataset not only separates excitatory neurons from different developmental stages and layers but also brain areas, e.g. early-born deep layer/subplate excitatory neuron in visual and prefrontal cortex area (EN-V1-1 and EN-PFC1, respectively). CellWalker2 successfully mapped ExM to early and late-born excitatory neurons (EN-PFC3 and EN-V1-2), while mapping most ExM-U cells to late-born excitatory neurons (EN-V1-3) (Figure 4B), consistent with upper layer neurons developing later[29] and high marker overlaps between these mapped pairs of cell types (Figure 4B). In contrast to CellWalker2, Seurat maps a large portion of ExM cells to newborn excitatory neurons (nEN-early2) (Figure 4B), which is dubious given that these cells are in different maturation stages and the fact that ExM shares more marker genes with EN-V1 and EN-PFC cell types compared to nEN-early2 (Figure 4B). It is also consistent with our simulation results showing that Seurat is biased towards prevalent cell types, because nEN-early2 has the highest number of cells among all the excitatory neurons (∼27%).

When the reference labels contain information absent from the query cell ontology, CellWalker2’s cell-to-label mappings can refine our understanding of cell types in the query dataset. For example, ExDp1 cells are divided into deep layer excitatory neurons from different brain areas (EN-V1-1 versus EN-PFC1) based on their mappings to the Nowakowski et al. dataset (Figure S16a). Although all of these labels indicate early-born deep-layer excitatory neurons, the >1.5-fold expression differences of area markers, including *KCNJ6* and *SATB2* for PFC and *TENM2* for V1, validate the annotation of these cells as originating from distinct areas (Figure 4C). Moreover, these area-divided subgroups also express different laminar layer markers, including a 4-fold change for the subplate marker *NR4A2* and a 2-fold change for the layer 5/6 marker *CRYM* (Figure 4C). They also show different timings of neuronal cell birth (Figure 4D in [29]). Another example is ExM, for which subgroups of cells are annotated as two different cell types that show differential expression of markers for upper versus deep layer clusters identified in [30] (Figure 4D). By taking account of the cell type hierarchy, CellWalker2 can identify cells at an intermediate state between cell types. For instance, some ExDp1 and ExM cells were mapped to the ancestor cell types (Figure 4E and F), and the UMAP and expression level of markers support that this group of cells represents an intermediate state between two tip cell types (Figure S16b and c). Together, these results highlight the power of our cross-study cell annotation for refining cell classifications and underscore the relationship between neuronal cell types and their migration patterns during development.

#### Mapping cell type hierarchies by scRNA-Seq data

We then applied CellWalker2 to map cell type hierarchies of these two developing human cortex datasets and compared the results with treeArches and MARS. We mapped the cell types in the Polioudakis et al. dataset (query) onto those from Nowakowski et al. (reference), and also flipped them to evaluate if our findings were consistent in both directions. CellWalker2 and treeArches showed similar results overall, but MARS was quite different (Figure S17) and made errors for several rare, non-neuronal cell types.

Places where results from CellWalker2 and treeArches differed revealed the effects of several of our modeling choices. First, these tools handle cell types in the query ontology that do not strongly match a single cell type in the reference ontology differently; treeArches treats this as a new cell type and assigns it to the root node, whereas CellWalker2 generates a Z-score for all ancestral and leaf nodes. For example, CellWalker2 maps (highest Z-score) ExN to an ancestor node (Figure 5A), suggesting that excitatory neurons are more broadly defined in Polioudakis et al. than in Nowakowski et al.. This is supported by cell-to-cell distances from ExN to other cell types (Figure 5B and Figure S18). We observed that such mappings to internal nodes depend on CellWalker2’s use of a null distribution, because influence scores do not show the same adaptability and are maximal at the leaf nodes (Figure S19). treeArches, on the other hand, maps ExN to the root node (highest probability), and terminal cell types that have higher prevalence. Similar behaviours of both methods have been observed for mapping two types of inhibitory neurons (InCGE and InMGE) (Figure 5A, details in Note S2). If we manipulate the cell type resolution of the Nowakowski et al. ontology, for instance by amalgamating excitatory neuron cell types, treeArches assigns larger scores to the combined cell type and reduces its score at the root node (Figure S20b). Alternatively, if we remove cell types with the highest scores for ExN, treeArches’ scores increase for other cell types, including a prevalent cell type from a different developmental stage (Figures S20c and S20e). On the other hand, CellWalker2’s top cell types stay the same with only minor decreases in Z-scores. This suggests a second difference between CellWalker2 and treeArches that inclusion of closely related cell types in a dataset may lower treeArches’ mapping scores to each of them, particularly rare cell types due to treeArches’ sensitivity to compositional bias, but encourages CellWalker2’s mapping to all these cell types and their ancestor. These findings indicate that treeArches may be better able to map cell types when the two ontologies have comparable resolution, while CellWalker2’s use of internal nodes enables mappings between fine-resolution and broad cell types.

**Figure 5:**
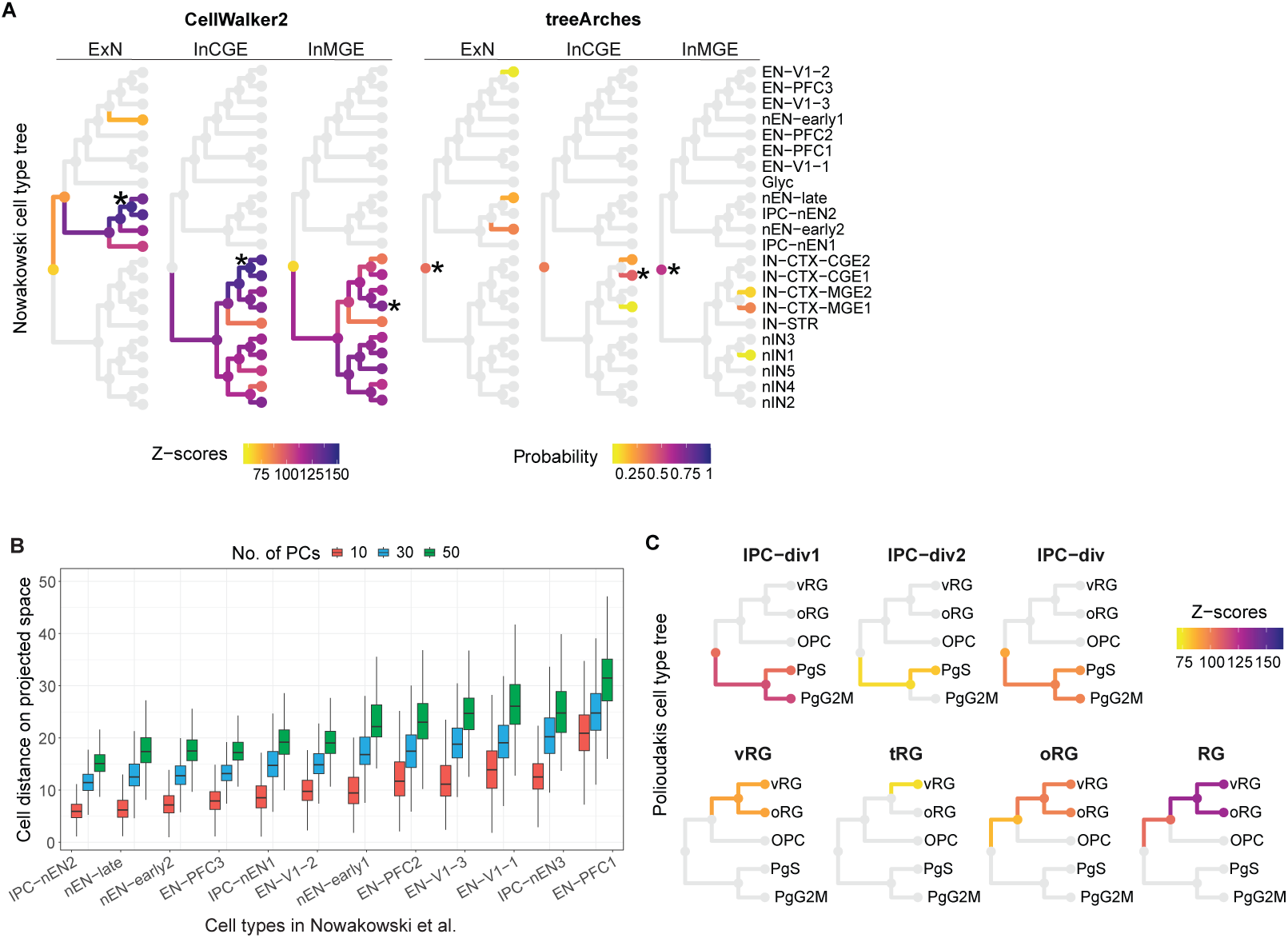
Mapping cell type hierarchies using scRNA-Seq data in human developing cortex. (A) Mapping ExN, InCGE, InMGE in Polioudakis et al.’s dataset onto the cell type hierarchy from Nowakowski et al. by CellWalker2 or treeArches. The nodes with the largest scores are indicated by ‘*’. The rejection probability that treeArches could not identify mapped cell types are 0.1, 0.03 and 0.02 for ExN, InCGE and InMGE respectively, which are not shown on the figure. (B) ExN is similar to the cell types mapped by CellWalker2, i.e. IPC-nEN2, nEN-late and nEN-early2. Euclidean distance between ExN cells from Polioudakis et al. and excitatory neurons of various types from Nowakowski et al.. Boxplot shows the distribution of cell-to-cell distances computed on the top principal components (PCs) using a subsample of 2000 ExN cells. The cell types on the X-axis are ordered by median cell-to-cell distance using 30 PCs. (C) Mapping radial glia (RG) and intermediate progenitor (IPC-div) cell types in Nowakowski et al. onto the cell type hierarchy of Polioudakis et al. using CellWalker2. IPC-div1 and IPC-div2 are two types of RG-like intermediate progenitor cells, with IPC-div as their parent; vRG, tRG and oRG are three types of radial glia, with RG as their ancestor. For (A) and (C), the branches and nodes are colored by Z-scores for CellWalker2 and by mapping probabilities for treeArches (Z-score > 75 and probability > 0.005 are shown). Explanation of abbreviations in Table S1 and S2.

As another example, CellWalker2 maps two types of radial glia in Nowakowski et al. (oRG, vRG) to all nodes in the radial glia subtree with roughly equal Z-scores, but maps another type of radial glia (tRG) weakly in the Polioudakis et al. ontology, which does not have a tRG cell type (Figure 5C). On the other hand, the ancestor nodes of all radial glia in these two datasets show strong correspondence with each other (Figure 5C and S21), indicating agreement at this broader level of classification. These findings indicate that CellWalker2’s ability to map to internal nodes of cell type hierarchies resolves problems that arise when cell type ontologies contain different cell types and non-unique cell type relationships.

Moreover, CellWalker2 facilitates the interpretation of cell types by mapping labels in another ontology that carry additional information about cell state. For the two groups of RG-like Intermediate progenitor cells in Nowakowski et al. (IPC-div1 and IPC-div2), although the Nowakowski et al. labels do not reference the cell cycle, the mapping result from CellWalker2 suggests that G2M versus S phase is one differentiating factor between them (Figure 5C). The reverse mapping of PgG2M and PgS to the cell types in Nowakowski et al. supports this conclusion, while also identifying a few other types of dividing and progenitor cell types with potential enrichment for cells in G2M or S phase (Figure S21).

Finally, since the threshold for calling marker genes is usually a subjective user choice, we showed that CellWalker2 is not very sensitive to the number of marker genes for each cell type, and that including more markers does not substantially alter the results (Figure S22a). As single cell data is often noisy, we also showed the robustness of CellWalker2 upon varying cell type composition and adding different intensities of noise to the cell-cell similarity matrix (Note S3; Figure S22b). Altogether, these analyses of scRNA-seq from the developing human brain demonstrate the robustness of CellWalker2, the importance of using a null distribution to compute Z-scores, and the flexibility and potential for new understanding of cell states that arises from using all nodes of cell type hierarchies.

### Cross-species comparison of neurons in mammalian motor cortex

We applied Cellwalker2 to BICCN scRNA-Seq and SNARE-Seq data from adult human, marmoset, and mouse motor cortex samples [31]. Although Bakken et al. mapped the cell types from each species to a cross-species consensus taxonomy, they did not assess statistical significance, nor did they investigate relationships across levels of the cell type hierarchy. To explore these gaps, we applied CellWalker2 to both cell types and subclasses (STAR Methods), investigating species individually and jointly, using graphs with around 10,000 cells per species. These analyses showed that CellWalker2 can compare GABAergic inhibitory neuron cell type hierarchies across species and leverage cell type-specific gene expression or chromatin accessibility to identify pathways and transcription factors that with conserved or divergent evolutionary patterns of cell type specificity.

#### Mapping subtypes of inhibitory neurons across species

First, we ran CellWalker2 to map cell types between the human and marmoset inhibitory neurons at the subclass level (Figure 6A), observing that cells from one species map to the corresponding subclass in the other species whenever one exists, and subclasses cluster into CGE-derived versus MGE-derived subclasses based on their Z-scores (Figure S23). Then, we applied CellWalker2 to map cell types across species within each subclass. Some cell types in marmoset can be mapped to a single cell type in human with a high Z-score. For example, the chandelier cells (Inh PVALB FAM194A) in marmoset, mapped to Inh PVALB COL15A1 in human which are also chandelier cells (Figure 6B). Although these two cell types have different cell type markers in human versus marmoset species, they share similar expression profiles that enable CellWalker2 to connect them and distinguish them from other Pvalb cells. On the other hand, some cell types receive similar Z-scores for multiple cell types within a group (Figure 6B). For instance, Inh SST ABI3BP is mapped to a subtree with three tips, all of which belong to a Sst cell cluster (Sst_3) in the consensus taxonomy of Bakken et al.. These high-scoring cell types are all present in upper layers and share many markers with Inh SST ABI3BP, while the other cell types in Sst_3 are associated with deep layers and share fewer markers (Figure 6C), suggesting that CellWalker2 refined the consensus taxonomy. Overall, cell type mapping results within subclasses show nested block-wise structure, indicating that consensus subgroups within subclasses of inhibitory neurons exist between human and marmoset (Figure S24). These results demonstrate that CellWalker2 can provide a more nuanced and hierarchical mapping between cell types than is possible with a consensus taxonomy.

**Figure 6:**
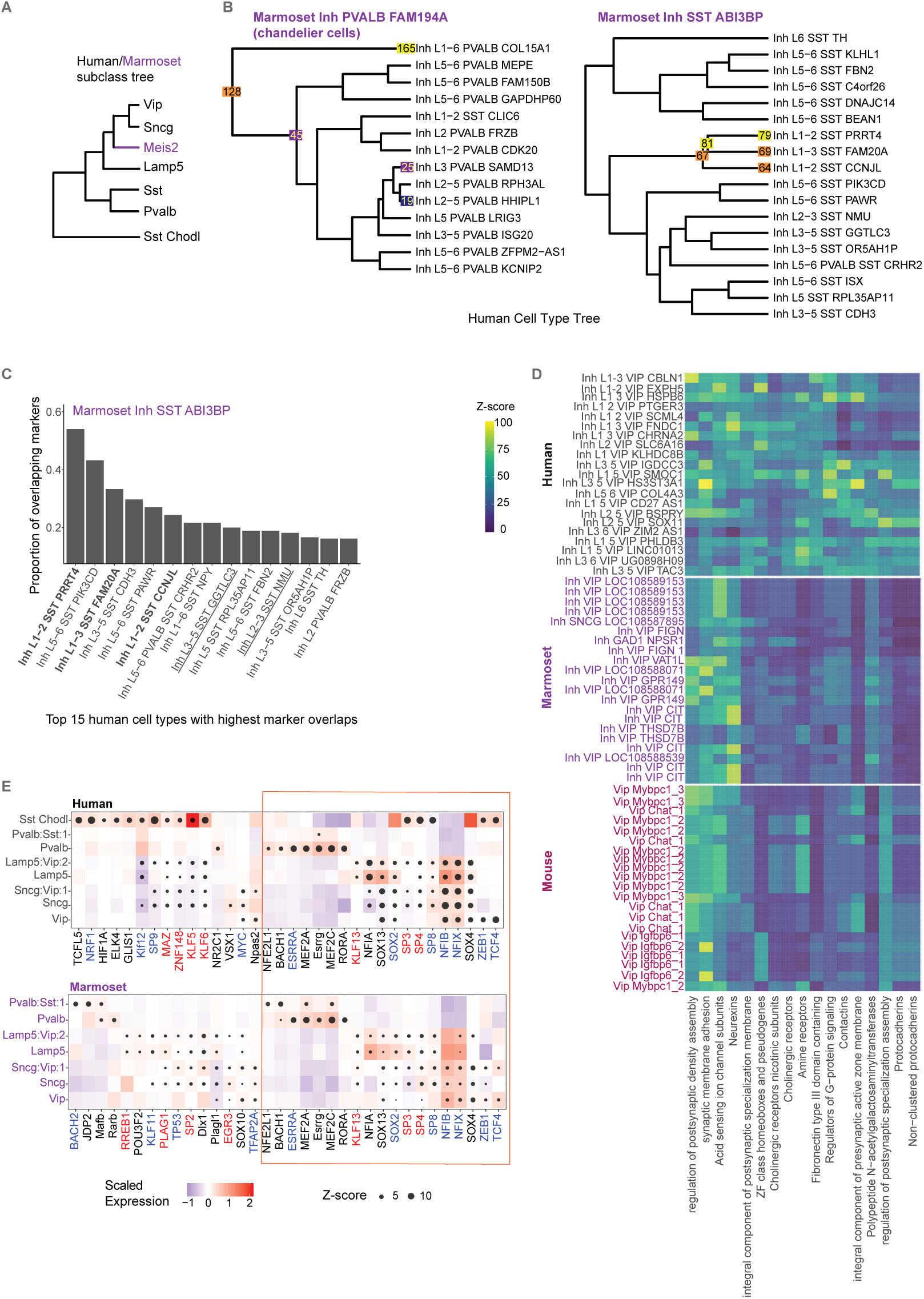
Cell type mapping, consensus and divergent transcription factors, and gene sets across species. (A) The marmoset and human cell consensus taxonomy of cell subclasses. Marmoset has Meis2 cells (purple) which are not present in the human ontology. Other subclasses are shared. (B) Marmoset cell types map onto the human cell type tree with either clear one-to-one relationships or with similar Z-scores for all nodes in a subtree of related cell types (e.g., Inh PVALB FAM194A and Inh SST ABI3BP). Human Inh PVALB (left) and SST (right) subtrees are shown. The top 5 cell types (largest Z-scores) are shown. The number and color of each node both reflect the magnitude of the node’s Z-score. Cell type names are based on two marker genes [31]. Human cell type names also contain laminae layer information. (C) High proportions of cell type markers overlap between marmoset Inh PVALB FAM194A cells and mapped human cell types. The top 15 human cell types with the largest overlap of positive marker genes are shown. The three cell types with the largest Z-scores are shown as bold and the other two cell types in the human and marmoset consensus cluster (Sst_3) definded in Bakken et al. are underlined. (D) Gene sets that are activated in human VIP cells show different cell type specificity across species. Three heatmap panels show the Z-scores (color scale) for mappings between gene sets and cell types in human (left), marmoset (middle), and mouse (right). Some of the pathways are active in all three species (e.g., neurexins, acidsensing ion channel subunits, synaptic and postsynaptic related pathways), while others are only active in human (e.g., protocadherins, cholinergic receptors, amine receptors, Contactins, regulation of G-protein signaling and presynaptic active zone membrane). Rows: gene sets, Columns: cell types. For each human cell type, the top matched cell type by CellWalker2 in marmoset or mouse is shown. (E) CellWalker2 identifies both consensus and unique cell subclass specific TFs in human and marmoset inhibitory neurons. See STAR methods for selection criteria. Columns: TFs, Orange box: consensus TFs between human and marmoset, Rows: node in cell subclass tree with internal nodes named by their two descendent nodes and depth in the tree. Rows are ordered identically in both species, except the additional Sst chodl cell type in human. Sst is not shown as no significantly associated TFs were found. The size of the dot represents Z-score. The color of each square is the standardized gene expression of a TF in a particular cell subclass. Red font: universal stripe factors, Blue font: other stripe factors.

Next, we mapped the entire cell type hierarchies in human and marmoset (Figure S25). Cell types within the same subclass have higher mapping scores in general, but some cell types from different subclasses have detectable similarity (Figure S26). For example, human Sst chodl cells, grouped together with other Sst cell types in the human ontology, not only map to Sst chodl in marmoset, but also to another subtree of Sst cell types in which Inh SST MPP5 is also labeled as Sst chodl in the consensus taxonomy [31]. This indicates that human Sst chodl cells have features of both chodl cells and some other Sst cells in marmoset, which are separate lineages of the marmoset cell type tree. Another example is a marmoset Sst cell type (Inh PVALB SST LRRC6) that expresses both *PVALB* and *SST* and shows similarity not only to the human Sst subtree but even higher similarity to the Pvalb subtree, consistent with this cell type having features of both Pval and Sst cells. The relationship between cell types across species could be more complicated than matching cell type trees using a consensus taxonomy, as the relationship between cell types can change as species evolve. CellWalker2 successfully identifies cell type similarities across different lineages of a cell type tree, including cases where a cell type in one species has features of multiple separate lineages in the other species.

Finally, we used CellWalker2 to integrate scRNA-Seq data from all three species. By including cells from all three species in the same graph, we can compare the cell type similarity between pairs of species. We observed that the mapping (Z-score) between human and marmoset cell types is in general stronger than that of human and mouse cell types (Figure S27a and S27b), as expected given the species’ evolutionary relationships. However, there are cases where a human cell type is more similar to some cell types in mouse than to any in marmoset (Figure S27c). By identifying the most similar cell type across non-human species, CellWalker2 can nominate the best animal model to study a particular disease-related cell type.

#### Comparison of cell type-specific gene sets across species

We next used CellWalker2 to investigate activated gene pathways from HGNC and SynGO [31] in each cell type (i.e., gene sets with higher average expression in a particular cell type) and compare these between species (STAR Methods). The Z-score matrix from CellWalker2 shows a similar pattern across species (Figure S28), meaning that the majority of gene sets have conserved cell type specificity. However, we also observed differences in gene set activity across species. Some of the differences come from cell types that do not exist in all species (e.g. the Meis2 subclass in Figure S28c). Other gene set activity differences come from cell types that are shared across species, but in which a gene set is more highly expressed for one species compared to the other. For example, in the Lamp5/Sncg subclass, Glutamate receptors and Catenins, which are important for normal cognitive development, show higher activity in several cell types for human compared to mouse, and to a lesser degree compared to marmoset (Figure S28d). For the Sst subclass, Glutamate ionotropic receptor kainate type subunits have a very high Z-score in the human SST GGTLC3 cell type, but lower Z-scores in marmoset and mouse (Figure S28e). *GGTLC3* is involved in gamma-glutamyltransferase activity and is associated with autism spectrum disorder [32]. Several genes in glutamate ionotropic receptor kainate type subunits (*GRIK1* and *GRIK3*) are marker genes of the human SST GGTLC3 cell type. Moreover, some pathways related to postsynaptic membrane have higher activity in human Sst cells than marmoset and mouse. For the Pvalb subclass, the active pathways in human are fairly conserved in mouse and marmoset, except that the small integrin binding ligand N-linked glycoprotein (SIBLING) family has higher Z-score in human (Figure S28f). The SIBLING family has been shown to affect cellular proliferation, differentiation and apoptosis including survival of dopaminergic neurons [33]. Gene sets that are activated in human VIP cells also show different cell type specificity across species (Figure 6D). Compared to gene set enrichment analysis, our approach does not need to label the cells first, which can be difficult for closely related cell types. In addition, CellWalker2 highlights the gene sets with higher average expression in a particular cell type rather than higher proportion of differentially expressed genes, without needing to select a differential expression threshold.

#### Cell subclass-specific regulatory regions and transcription regulators in human and marmoset

To elucidate the similarities and differences of the regulatory machinery across species, we applied CellWalker2 to compare DARs in different cell types and potential TFs that bound to these regions in motor cortex between human and marmoset using BICCN SNARE-Seq data. First, applying CellWalker2 on human DARs reveals that human cell type-specific DARs have higher chromatin accessibility in multiple cell types of the corresponding marmoset cell subclass, indicating that human and marmoset have similar open chromatin signatures on the subclass level (Figure S29). The limited number of marmoset cells makes it hard to map DARs on the cell type level.

Next, to investigate whether different species share cell type-specific transcription factors, we used CellWalker2 to identify TF motifs that are associated with open chromatin regions in different cell types. TFs with similar or co-occuring motifs could have similar Z-score profiles, so we further filtered out TFs not expressed in the corresponding cell subclass and compared the results between human and marmoset (Figure 6E). Since the marmoset dataset has a much smaller sample size, Z-scores are smaller in marmoset than that in human. Still, a lot of top TFs are shared between human and marmoset within similar cell subclasses (Note S4). Many of these TFs are stripe factors, which occupy regulatory regions broadly and have key roles in tissue-specific transcription [34]. CellWalker2 also discovered human unique cell type-specific TFs, such as *NPAS2*, *MYC* and *VSX1* in Vip and Sncg cells, and marmoset unique TFs (Note S4). *NPAS2* regulates GABAergic neurotransmission and associated with psychiatric disorders [35]. *MYC* promotes neuronal differentiation in developing neural tube [36], and *VSX1* contributes to interneuron development [37, 38]. Moreover, we also observed that CellWalker2 identifies more cell subclass-specific TFs compared to ArchR (Note S4).

In summary, CellWalker2 identifies TFs that potentially determine cell type identity and enables comparisons of these TFs across different species with measures of statistical significance.

## Discussion

The simulations and data explorations presented here have shown that CellWalker2 provides accurate and biologically meaningful statistical measures of association between cells and cell types, amongst cell types from different ontologies, and between cell types and annotations such as transcription factor binding motifs, gene sets, and genetic variants. Cell types can be represented as hierarchical ontologies that enable flexible labeling of cell states, including integration of datasets with different resolutions. Through carefully designed permutations, CellWalker2’s Z-scores quantify these associations in an unbiased manner. This enables users to detect associations across heterogeneous contexts, including different studies, disease states, or species. Cross-species comparisons with CellWalker2 enable researchers to trace the evolution of gene regulation at the cell type level, with potential to reveal the molecular mechanisms through which species diverge and specialize. Comparing cell types between human and other species can also help us identify suitable model organisms for studying human diseases.

Our analyses highlighted several features of CellWalker2 that contribute to its performance. One key example is CellWalker2’s ability to estimate a global measure of statistical significance for its mappings. The permutation null distribution preserves the marginal distribution of edges weights from cell type labels or from annotations. Thus, the algorithm will not bias towards prevalent cell types or annotations that occur many times in the genome. The resulting Z-scores are robust to variability in cell composition (e.g., dropping or merging cell types), sequencing depth, the number of cell type markers, noise in the cell-to-cell graph, or how cell types are defined. Another important distinction of CellWalker2 is its use of hierarchical cell type ontologies, including computation of Z-scores for not just terminal nodes representing specific cell types, but also all ancestral nodes. This enables CellWalker2 to identify a broader cell type mapping when a fine-resolution one does not exist. Furthermore, CellWalker2 incorporates a strategy to automatically tune model parameters to optimize performance, i.e. tune the weight parameters between cell-to-label to cell-to-cell edges, which are expected to depend on the relative confidence of cell-to-label edges but are usually unknown. Altogether, these design choices contribute to making CellWalker2 a graph-based model that can mitigate various sources of noise in single cell experiments.

CellWalker2 was designed to be highly flexible. For example, it can utilize different modalities of single cell data to map cells or annotations to cell type labels. This includes being able to model a mixture of cells from different experiments, some of which have ATAC-seq only, some with RNA-seq only, and others with multi-modal ATAC-seq and RNA-seq. Cells with all three combinations of data are not required, and indeed some applications only require RNA-seq (e.g., mapping labels between two ontologies). CellWalker2’s graph model can handle all of these combinations, enabling integration of datasets acquired using different single-cell platforms. Furthermore, CellWalker2 does not assign cells to specific cell types; instead, it treats cells as nodes in an interconnected graph. This approach proves advantageous for modeling overlapping cell states within the developing brain or delineating fine-grained cell subtypes in blood. In summary, CellWalker2 is an adaptable and robust tool that sheds new light into the cellular diversity of complex tissues and the genome sequences contributing to this diversity.

## Limitations

Though CellWalker2 provides statistical significance of the association between bulk-derived labels and cell types, it does not output the components of a regulatory module, such as regulatory regions and downstream target genes of a transcription factor. Other methods, like SCENIC+, FigR[39], and Pando[40] can identify regulatory modules. However, CellWalker2 can be utilized to filter for key cell types where a regulatory module is potentially active.

Another caveat is that the permutation schema in CellWalker2 does not directly utilize relationships between cell types, which would potentially underestimate the standard deviation of the influence score (Note S5). We also found that CellWalker2 could potentially assign higher Z-scores to rare cell types. When too few edges connect to a cell type label, the influence score under permutation would be close to zero and its standard deviation could be underestimated. We suggest that users increase the rounds of permutation in this case.

Lastly, for identifying cell type-specific TFs, CellWalker2 only considers motifs appearing in genomic regions that are more accessible in certain cell types, which could lead to false positives as many TFs have similar motifs or frequently co-occur in the same regions. To reduce false positives, we further filtered TFs, retaining those that are expressed in given cell type and have positive correlation between expression and Z-score. A more systematic approach is a focus for future work.

## Acknowledgements

This research was supported by National Institute of Mental Health (NIMH) grant numbers U01MH116438 (KSP), R01MH109907 (KSP), R01MH123179 (KSP), and Additional Ventures. We also thank Alex Pollen, Tomasz Nowakowski and Nadia Roan for discussions on the results, Ryan Corces for help on using ArchR, Sean Whalen for providing information on pREs and all Pollard lab members for suggestions on this project.

## Author contributions

KSP conceptualized the study, contributed to data analysis and interpretation, and revised the manuscript. ZH developed the method, conducted the experiments, analyzed the data, and contributed to the manuscript writing. PFP initialized the study, contributed to the study design, provided critical feedback on data analysis, and revised the manuscript. All authors read and approved the final version of the manuscript.

## STAR Methods

### Specifying edge weights for the graph

The graph in CellWalker2 includes four types of edges: cell-to-cell, cell-to-label, cell-to-annotation and label-to-label. The label-to-label edges are optional (i.e., users can work with discrete cell types without defining their relationships to each other), and in this study we focused on the commonly used label-to-label graph structure of a binary, hierarchical tree, although CellWalker2 can utilize ontologies with other topologies.

For cell-to-cell edges, CellWalker2 first computes the cell-to-cell similarity based on gene expression and/or chromatin accessibility profile of the cells and constructs a K nearest neighbor (KNN) graph (K = 30 by default). The cell-to-cell edge weight is based on shared neighbors on the KNN graph. For cells with RNA-Seq data, the cell-to-cell similarity is based on the correlation of gene expression profiles. CellWalker2 projects the gene expression data onto a low dimensional space (dim = 30 by default) using PCA and the cell-to-cell distance is the Euclidean distance in the latent space. The low dimensional space is defined using all the cells with RNA-Seq data. We normalize the distances by their largest value to make them between 0 and 1 (denoted as d), and the cell-to-cell similarity is defined as −logd. We also standardize the similarities to make them comparable with other data modalities. For cells with ATAC-Seq data, the cell-to-cell similarity is computed as the Jaccard or Cosine similarity of the vectors of peak presence/absence in each cell. We include peaks that appear in 0.2%-20% cells. Then we take the logarithm and standardize the similarities to make it comparable with other modalities. Although CellWalker2 provides several distance metrics for the similarity of scATAC-Seq profiles, including Jaccard, Cosine and latent semantic indexing (lsi), in our experiments, we observe that these metrics perform similarly. For cells with multiomic data (i.e., both RNA-Seq and ATAC-Seq), the cell-to-cell similarity is a weighted average of RNA-Seq and ATAC-Seq similarity. If we have both unpaired scATAC-Seq and/or scRNA-Seq data and multiomic data, we use multiomic data as a bridge to connect cells with unpaired scATAC-Seq and/or scRNA-Seq data. We compute cell-to-cell similarity between cells from multiomic and unpaired scRNA-Seq data using gene expression profiles and from multiomic and unpaired scATAC-Seq data using chromatin accessibility profiles. Then we combine the cell-to-cell similarity matrices from various modalities of the data, construct a KNN graph, and obtain a joint cell-to-cell graph (Figure S30).

The cell-to-cell type label edge weights are based on the gene expression level of the marker genes of each cell type. The edge weight between a cell and a cell type label is defined as: 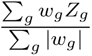. The summation is over all marker genes for a cell type, and w_g_ is the weight of each marker gene which users can specify (by default, log fold change between mean expression level in one cell type versus the rest), and Z_g_ is the standardized gene expression in the cell. If cell type labels have graph structure, such as a hierarchical tree, we include internal nodes of the cell type ontology into the graph and connect internal nodes with tips. The edges reflect the (hierarchical) relationships between cell types. For a binary tree, the edge weight going up or down the tree could be unequal to limit the random walks from going through the root node and passing information from one side of the tree to the other.

Annotations can be gene sets or any genome coordinates of interest, such as a group of regulatory elements, genetic variants, or TFs. The edge weight between cells and gene set is the average standardized expression level of all genes in the gene set. On the other hand, the edge weight between cells and genome coordinates is the overall accessibility of the genomic regions (i.e., we sum up all the reads in the ATAC-Seq peaks within those regions). For TFs, we identify motifs of interest, such as those within bulk or cell type specific (DARs) open chromatin regions. We identified TF binding motifs from JASPAR2020 [41] in each region, and connected a TF to a cell by summing up that cell’s ATAC-seq reads within genomic regions that contain the TF motif.

Lastly, we tune the weights between cell-to-cell and other types of edges. If the task is to map cell type labels, we optimize the cell homogeneity score [30] such that labels can classify all the cells well. If the task is to map annotations to cell types, we minimize the entropy of influence scores between labels to cell types (‘label entropy’) such that the scores are more specific to certain cell types. The weight between label-to-label and cell-to-label edges is also tunable, depending on how deeply users wish to map cells on a hierarchical cell type ontology. If the weight between labels is large, the random walk will be more likely to reach the internal nodes of the tree or even the root node.

### Computing Z-score from the influence score matrix

To estimate the Z-score between any pair of cell type labels, we compare the observed value for the corresponding entry of the influence matrix with its null distribution assuming independence between cells and labels. To generate this null distribution, we permute the edges between cells and cell type labels. In detail, we re-sample the edge weights between cells and labels while keeping the marginal distributions of edge weights of each cell and each label stay close to its empirical distribution. First, to estimate the marginal distribution, we discretize the edge weights into 0 plus 10 equally-spaced intervals between 0 to 1 for each cell type label and 5 intervals for each cell as the number of labels connecting to each cell is smaller than the number of cells connecting to each label. We compute the proportion of edge weights in each interval for each cell or label. We tried discretizing into quantiles of edge weights, but the results were worse because the edge weight distributions usually have long tails. Second, for each cell type label, we resample the cells connecting to that label while preserving the number of cells in each edge weight interval. The probability to sample a cell is proportional to the probability that the edge weights from that cell lie in such interval. Finally, we uniformly sample values within each interval as the new edge weights. We also implemented different permutation strategies, either permuting the edges from cells to one of the cell type hierarchies or both. A cell is more likely to be linked with nearby cell types in the hierarchy, which will be lost after permutation. Preserving the edges between cells to cell type labels in generating the null distribution of influence scores can maintain the correlation between cell types. Thus, we recommend users permute edges to the cell type hierarchy of the query dataset. However, this might reduce the power in detecting associations.

To generate a null distribution for mappings between annotations and cell type labels, we permute the edges between cells and cell type labels using the strategy above. To estimate the Z-scores for cell type-specific TFs, we permute the appearance of TF motifs in genomic regions. For each TF motif, we resample the genomic regions that contain it. The probability of choosing each genomic region is proportional to the number of motifs in that region. Then we recompute the edge weight between each TF and a cell by summing up all the reads in the regions that contain the TF after permutation while keep other edges the same. We found that this strategy performs better than permuting cells to cell type labels in identifying more cell type-specific TFs than common TFs. The reason might be that as connections from regions to cell types are kept when generating null distribution, any cell type bias of the chromatin accessibility of input regions will also be maintained in the null distribution so they won’t be reflected in Z-scores. Moreover, by permuting TF-to-region edges when identifying cell type-specific TFs, CellWalker2 ensures the preservation of the correlation between cell types in the cell type tree.

We recommend sampling 50-100 permuted graphs. For each one, we compute an influence matrix using each randomized graph and estimate the Z-score by 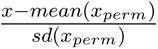 for entry x in the influence matrix. The Z-score reflects the statistical significance of the entries of influence matrix.

### Details for simulating scRNA-Seq data

In the first simulation without batch effect (easy), we simulated 4000 cells with 1000 cells in each of four cell types, 1400 non-marker genes, 200 marker genes between cell type (1,2) and (3,4), 150 marker genes which are differentially expressed between 1 and 2 but not in 3 and 4, and another 150 marker genes between 3 and 4. Then, we split the cells into two equal-sized reference and query datasets. In the simulation scenario 2 (medium) with batch effect and dropouts, we simulated 1000 non-marker genes, 200 marker genes between cell type (1,2) and (3,4), 200 marker genes between 1 and 2 and between 3 and 4 respectively. We added batch effects between these two datasets using the default parameters in Splatter. We set ‘dropout.mid = 0, 2’ and ‘dropout.shape = −1, −0.5’ in Splatter such that the query dataset has more dropouts than the reference one. We also varied the number of cells of cell type 4 in the reference dataset (100, 300 and 500 cells), but kept the number of cells as 500 for others. In the simulation scenario 3 (hard) with batch effect and more dropouts, we simulated 1200 non-marker genes, 100 marker genes between cell type (1,2) and (3,4), 150 marker genes between 1 and 2 and between 3 and 4 respectively with ‘de.facScale = 0.2’. We added batch effects between these two datasets using the default parameters in Splatter. We set ‘dropout.mid = 1, 2’ and ‘dropout.shape = −1, −0.5’ in Splatter.

To apply CellWalker2 for cell type annotation, we either used cells from the query dataset only or combined cells from both datasets to generate cell graph. For the simulation varying the number of cells, we used cells from both datasets. The marker genes for each cell type were computed using Seurat given the true cell type labels of cells in the reference dataset. We filtered for genes with | log_2_ FC| > 0.5 and adjusted p-value < 0.05. We input the true cell type hierarchy when used. For Seurat, we integrated cells from both datasets and used the ‘TransferData’ function to transfer labels from reference to query dataset. As Seurat computes a joint embedding of cells from both reference and query dataset, we denoted its results as “ref+query” in Figure 2.

We designed four other simulation scenarios where cell types in dataset 2 (DS2) are differ than those in dataset 1 (DS1) (Figure S5) in order to test performance for mapping cell types. We simulated batch effects between these two datasets with ‘batch.facLoc = 0.01’ and ‘batch.facScale = 0.1’, used default dropout rates for both. We simulated cell types A,B,C,D in DS1 and A,B,C,E in DS2 with 400 cells per cell type in each dataset. Cell type E was added to the cell type hierarchy in DS1 in several different ways. For ‘Divergent cell type’, cell type E is a new cell type in the lineage of C and D. We simulated 1000 non-marker genes, 200 marker genes between (A,B) and (C,D,E), 200 marker genes between A and B and 300 marker genes among C, D, and E. For ‘Ancestor cell type’, cell type E is the ancestor cell types of C and D a.k.a. cell type CD. We simulated 1000 non-marker genes, 200 marker genes between (A,B) and (C,D,E), 200 marker genes between A and B and 300 marker genes between C and D. For ‘Altered cell type’, cell type E being a slightly altered cell state of cell type D, i.e. E is more similar to D than C. We simulated 900 non-marker genes, 200 marker genes between (A,B) and (C,D,E), 300 marker genes between A and B, 200 marker genes between C and (D,E), and 100 marker genes between D and E with smaller fold changes. For ‘Convergent cell type’, cell type E sharing cell type markers from both cell type D and B from different lineages, i.e. E is in the lineage of C and D but share some features with B. We simulated 1000 non-marker genes, 100 marker genes between (A,B) and (C,D,E), 300 marker genes between A and (B,E) and 100 marker genes between C and (D,E). In the simulation where we varied cell numbers, we adjusted the number of cells of cell type D in DS1 to 100, 200, and 400 in the ‘Divergent cell type’ case, while maintaining 400 cells for the others. To isolate the effect of cell numbers, we only included cell type A,B and E in DS2 for this simulation, so cell type E should have equal distance to cell type C and D in DS1. Each simulation scenario is repeated 50 times.

To apply CellWalker2 for cell type mapping, we combined single-cell data from both datasets to generate cell graphs and obtained cell type markers for both datasets using the procedure described above. We connected all the cells to both sets of labels. For treeArches, we input the combined data from both datasets and used scVI to remove batch effect and project onto a 20 dimensional latent space, as in the default pipeline of treeArches. For MARS, we input integrated and scaled data, after removing batch effects using Seurat as MARS does not have a batch effect removal step integrated. We set pretrain epochs to 50 and used 500 and 200 for hidden dimensions 1 and 2, respectively, in the neuron network model. For both CellWalker2 and treeArches, we input the true cell type labels in each dataset, but for MARS, we can only input cell type labels in DS1 and it did *de novo* clustering for DS2. Therefore, for MARS, we mapped its clustering results with the cell type labels of DS2 by the Hungarian algorithm then obtained the relationships among cell types across datasets. We input cell type trees from both datasets for CellWalker2, which allows cell types to map to ancestral nodes of the cell type tree.

### Processing scRNA-Seq data in human developing cortex

The Nowakowski et al. dataset has better coverage over low expressed genes, contains more developmental stages, and collected samples from different brain areas and coritcal layers [29]. On the other hand, the Polioudakis et al. dataset has more cells, which might capture cells in transient states between cell types [30]. In addition to these factors, the cell type classification criteria of the two studies were based on different clustering algorithms.

We downloaded the scRNA-Seq raw counts from Polioudakis et al. [30]and Nowakowski et al. [29]. We obtained cell type hierarchies from the original publication. We selected marker genes with |avg_diff| > 1 from the supplementary material of Nowakowski et al.. For Polioudakis et al. dataset, we computed differentially expressed genes per cell type using Seurat and selected marker genes with | log_2_ FC| > 0.5 and adjusted p-value < 0.01.

For cell type annotation, we treated Nowakowski et al. dataset as reference and Polioudakis et al. as query. We subsampled around 7000 random cells from Polioudakis et al. to ease computational time. For CellWalker2, we used cells from Polioudakis et al. to build the cell-cell graph and marker genes of each cell type in Nowakowski et al. as labels. For Seurat, we integrated cells from both datasets and applied ‘TransferData’ function with latent dimension set to 30 to transfer labels from Nowakowski et al. to Polioudakis et al..

For mapping cell types between these two datasets, we integrated all the cells from both datasets, removed batch effects using Seurat, and incorporated cell type labels from both datasets into the graph. To obtain Z-scores, we permuted the edges from cells to cell type labels of Polioudakis et al.. We tried different permutation schema (e.g., permute cells to cell type labels of Polioudakis et al., cell type labels of Nowakowski et al., or both), but the result did not change much (Figure S31). Since we showed the results of mapping cell type of Polioudakis et al. to the cell type hierarchy of Nowakowski et al. in the main text, we used the first permutation schema as it maintains the correlation of cell types on the tree we mapped to. For treeArches, we input combined raw counts from the two datasets then used scVI as default to integrate the data. For MARS, we input integrated and scaled data after removing batch effects using Seurat, and set hidden dimensions 1 and 2 to be 1000 and 100 respectively in the neuron network model. We treated Nowakowski et al. as annotated data and Polioudakis et al. as unannotated data and mapped MARS identified clusters with the original labels in Polioudakis et al. to get the final cell type mapping results.

### Processing multimodal data in human developing cortex

We downloaded scATAC-Seq, scRNA-Seq, and 10X Genomics multomic data from GSE162170. We integrated scRNA-Seq and multiomic data from week 21, subsampled 9,000 cells from scRNA-Seq data, and used all the 8981 multiomic cells to construct the cell graph in CellWalker2. When further integrating with scATAC-Seq data from week 21, we subsampled 9000 cells with ATAC-Seq from week 21 and added them to the cell graph. When further integrating with scATAC-Seq data from week 16, we used all 6423 cells with ATAC-Seq from week 16. We used Seurat to integrate scRNA-Seq and the RNA-Seq part of multiomic data. We used the cell type labels from scRNA-Seq combining different ages during mid-gestation from the supplementary information of [24] for CellWalker2, and we built the cell type tree using ‘BuildClusterTree’ function in Seurat based on first 50 PCs of the gene expression data. We generated a cell type tree using the scRNA-Seq data, using only the leaf nodes in CellWalker but the full hierarchy in CellWalker2. We used the cell type labels from the multiome data for Wilcoxon and Fisher’s exact tests. A comparison for cell type labels between multiome and scRNASeq data is described in the Note S6.

We obtained predicted regulatory elements (pREs) from different brain regions (i.e., basal ganglia versus cortical plate, upper versus deeper layer pre-frontal cortex) in Markenscoff-Papadimitriou et al. [23], and connected region-specific pREs to cells by the overall accessibility in each set of regions. To compute Z-scores of the association between region-specific pREs and cell types, we permuted the edge weights between cells and cell type labels.

### Processing multispecies scRNA-Seq and SNARE-Seq data

#### mapping cell types across species by CellWalker2 using scRNA-Seq data

We downloaded scRNA-Seq data from human, mouse and marmoset from the BICCN data portal (https://data.nemoarchive.org/publication_release/Lein_2020_M1_study_analysis). By analyzing scRNA-Seq data in each species separately, Bakken et al. identified 72, 52 and 59 inhibitory neuron cell types in human, marmoset and mouse, respectively. These are grouped into 6 subclasses in human and 7 subclasses in marmoset and mouse, which have an additional Meis2 subclass. We obtained cell type hierarchy from the three species from the original publication [31], and identified cell type or subclass specific marker genes using Seurat after balancing the number of cells to 100 in each group. We used ‘roc’ test and selected markers with power > 0.65 for each species. We computed cell-to-cell type label edge weights within each species based on the log fold-change of marker genes and gene expression profiles of each cell. Then, we constructed the cell-to-cell graph based on the integrated gene expression data using consensus genes across species provided by [31].

#### Identify genes expressed specifically in different cell types

We used 767 expressed gene sets from HGNC and SynGO [31] as labels, and the weights of edges connecting each gene set and cells are the average standardized expression level of all genes in the gene set. The cell-to-cell graph and cell-to-cell type edges are the same as above. The edge weight between a cell and a gene set is the average standardized expression level in the cell, and we filtered for gene set that express in at least 20 cells. To obtain Z-scores, we permuted the edges between cells and cell type labels.

#### Comparison of cell type-specific regulatory regions in human and marmoset

We downloaded marmoset SNARE-Seq data from BICCN portal, which includes 1451 marmoset inhibitory neurons with both gene expression and chromatin accessibility. We used the same cell type labels and marker genes derived from scRNA-Seq as above. We used around 50K DARs from 18 human cell clusters identified from scATAC-Seq as labels [31]. To map human DARs to marmoset cell types, we used liftOver to transfer human DARs to the marmoset genome and removed DARs that are mapped to more than 3 regions. To run CellWalker2, we used marmoset cells to generate a cell-to-cell graph, treated human DARs from different cell types as annotation nodes and linked these nodes to each marmoset cell by computing the proportion of reads in the peaks overlapping with that group of DARs. To obtain Z-scores, we permuted the edges between cells and marmoset cell type labels.

#### Identify cell type-specific transcription factors in human and marmoset

We downloaded human and marmoset SNARE-Seq data with 22217 and 1451 cells, respectively, from the BICCN portal. We used the cell subclass label and identified markers by Seurat using the scRNA-Seq data. We treated TFs as annotation nodes in CellWalker2. We obtained subclass DARs from the supplementary information in [31] and used Signac to find the presence of 697 TF motifs from JASPAR2020 within these DARs. We further selected expressed TFs to run CellWalker2, which resulted in 379 TFs for human and 405 TFs for marmoset. Each TF is connected to cells through the chromatin accessibility of the DARs containing the TF motif. To obtain Z-scores, we permuted the genomic regions that contain the motif of each TF. We further computed the average standardized gene expression in each cell subclass using the RNA-Seq part of the SNARE-Seq data. For internal nodes, we averaged the gene expression across all the cells in its descendants. For each tip or internal node on the cell subclass tree, we selected TFs with Z-score > 2.5 and standardized expression > 0.2 or 0.15 for human and marmoset respectively to compare between human and marmoset. The threshold for calling expressed TFs of a cell type for marmoset is lower because we observed less variation of marmoset data. We varied these cutoffs to maximize the percentage of shared TFs between human and marmoset.

### scRNA-Seq and 10X multiomic datasets for human PBMC

For mapping cell types in PBMCs, we downloaded the scRNA-Seq dataset of healthy adult blood tissue from https://celltypist.cog.sanger.ac.uk/Resources/Organ_atlas/Blood/Blood.h5ad.We subsampled 20% of cells to run CellWalker2, resulting in 33123 cells from 46 cell types in Ren et al. [26]and 9361 cells from 33 cell types in Yoshida et al. [27]. We identified cell type markers and built a cell type tree using Seurat. We kept differentially expressed genes per cell type with | log_2_ FC| > 0.5 for Ren et al. and 0.25 for Yoshida et al. and adjusted p-value < 0.05 so that the number of cell type markers are in a similar range for both datasets. We used Seurat to integrate the two datasets based on the top 30 canonical directions and generated a cell-to-cell graph based on K nearest neighbors with k = 20. As we illustrated mapping cell types from Yoshida et al. to the cell type tree in Ren et al., we permuted cells to cell type labels from Yoshida et al. 100 times to get Z-scores. For treeArches, we input combined raw counts from two datasets then used scVI as default to integrate the data. For MARS, we input integrated and scaled data after removing batch effects using Seurat, set hidden dimensions 1 and 2 to be 1000 and 100 respectively in the neuron network model, and trained for 100 epochs. We treated Ren et al. as annotated data and Yoshida et al. as unannotated data and mapped MARS identified clusters with the original labels in Yoshida et al. to get the final cell type mapping results. For Z-score differences between mapping stimulated and un-stimulated monocytes from Yoshida et al. to monocyte related cell types in Ren et al., we used Monocyte CD14 IFN stim - Monocyte CD14, Monocyte CD16 IFN stim - Monocyte CD16, and Monocyte CD14 IL6 - Monoctyte CD14.

For identifying cell type-specific TFs, we downloaded 10X multiomic data from https://www.10xgenomics.com/datasets/pbmc-from-a-healthy-donor-granulocytes-removed-through-cell-sorting-10-k-1-standard-1-0-0. We followed the Signac pipeline https://stuartlab.org/signac/articles/pbmc_multiomic for calling peaks and cell-type annotation. Signac identified 131,364 peaks in total. We removed cell types with less than 30 cells to identify cell type-specific genes and peaks. We followed https://stuartlab.org/signac/articles/motif_vignette.html for identifying cell type-specific peaks and motifs from JASPAR2020 within these peaks. 58,385 cell type-specific peaks are identified using Signac (adjusted p-value < 0.05). We selected TFs that expressed in at least 100 cells, which results in 303 TF motifs. Then we ran CellWalker2 using the cell type labels, cell type-specific peaks and motifs within these peaks by Signac. We further computed the standardized expression level for each TF across cell types and selected TFs with standardized expression level > 0.5 and adjusted P-value < 0.01 (after converting Z-scores to p-values and applying Bonferroni correction) for each cell type, as well as Spearman’s rank correlation coefficient between expression level and Z-score > 0.4. For an internal node, the expression level is averaged across all the cells of its descendants. If a TF was selected for multiple nodes in the same lineage, it was only shown on the most ancestral node. We also ran motif enrichment analysis by Signac directly as a comparison. We used the same criteria as in CellWalker2 for filtering TFs after obtaining the enrichment p-values.

We ran ArchR according to https://www.archrproject.com/bookdown/index.html except using the cell type label by Signac for direct comparison. ArchR identified 169,218 peaks and 97,920 cell type specific peaks (FDR < 0.05 and log2FC > 1.5). Then we identified enriched motifs for each set of cell type-specific peaks. We used the same criteria as in CellWalker2 for filtering TFs after obtaining the enrichment p-values. We ran SCENIC+ according to https://scenicplus.readthedocs.io/en/latest/pbmc_multiome_tutorial.html#Tutorial:-10x-multiome-pbmc except using the cell type labels by Signac for direct comparison. We selected cell type-specific TFs with rho > 0.4.

## Supplemental information

Document S1. Figures S1–S31, Table S1-S2 and Notes S1-S6

## Data and code availability

CellWalker2 is available on Github https://github.com/PFPrzytycki/CellWalkR/tree/dev and the codes for simulations and data analysis in this article is available at https://github.com/xyz111131/CellWalker2_supple_codes/tree/main. scRNA-Seq dataset in Polioudakis et al.: http://solo.bmap.ucla.edu/shiny/webapp/. scRNA-Seq dataset in Nowakowski et al.:https://cells.ucsc.edu/cortex-dev/. Human developing cortex multimodal data: GSE162170. Mammalian motor cortex data from BICCN: https://data.nemoarchive.org/publication_release/Lein_2020_M1_study_analysis. Human PBMC scRNA-Seq data: https://celltypist.cog.sanger.ac.uk/Resources/Organ_atlas/Blood/Blood.h5ad. Human PBMC 10X multiomic data: https://www.10xgenomics.com/datasets/pbmc-from-a-healthy-donor-granulocytes-removed-through-cell-sorting-10-k-1-standard-1-0-0.

