## Supplemental Document for "CellWalker2: multi-omic discovery of hierarchical cell type relationships and their associations with genomic annotations"

**Supplemental Tables**

| Cell type | Group | Cell type full name |
|---|---|---|
| ExDp1 | ExDp | Excitatory neuron deep layer 1 |
| ExDp2 | ExDp | Excitatory neuron deep layer 2 |
| ExM | ExM | Maturing excitatory neuron |
| ExM-U | ExM | Maturing excitatory neuron (upper layer enriched) |
| ExN | ExN | Migrating excitatory neuron |
| InCGE | InCGE | Interneuron CGE |
| InMGE | InMGE | Interneuron MGE |
| IP | IP | Intermediate Progenitor |
| PgG2M | Pg-Div | Cycling progenitor G2/M phase |
| PgS | Pg-Div | Cycling progenitor S phase |
| oRG | RG | Outer Radial Glia |
| vRG | RG | Ventricular Radial Glia |

Table S1: Polioudakis's cell type groupings with abbreviations used in the text and figures.

| Cell type | Group | Cell type full name |
|---|---|---|
| EN-V1-1 | EN-early | Early Born Deep Layer/subplate Excitatory Neuron V1 |
| EN-PFC1 | EN-early | Early Born Deep Layer/subplate Excitatory Neuron PFC |
| EN-PFC2 | EN-late | Early and Late Born Excitatory Neuron PFC |
| EN-V1-3 | EN-late | Excitatory Neuron V1 - late born |
| EN-PFC3 | EN-late | Early and Late Born Excitatory Neuron PFC |
| EN-V1-2 | EN-late | Early and Late Born Excitatory Neuron V1 |
| IN-STR | IN-STR | Striatal neurons |
| IN-CTX-CGE1/2 | InCGE | CGE/LGE-derived inhibitory neurons |
| IN-CTX-MGE1/2 | InMGE | MGE-derived Ctx inhibitory neuron, Cortical Plate-enriched |
| IPC-div1 | IPC-div | Dividing Intermediate Progenitor Cells RG-like |
| IPC-div2 | IPC-div | Intermediate Progenitor Cells RG-like |
| IPC-nEN1/2/3 | IPC-nEN | Intermediate Progenitor Cells EN-like |
| MGE-div | MGE-div | dividing MGE Progenitors |
| MGE-IPC1/2/3 | MGE-IPC | MGE Progenitors |
| MGE-RG1/2 | MGE-RG | MGE Radial Glia 1/2 |
| nEN-early1/2 | nEN | Newborn Excitatory Neuron - early born |
| nEN-late | nEN | Newborn Excitatory Neuron - late born |
| nIN1-5 | nIN | MGE newborn neurons |
| vRG | RG | Ventricular Radial Glia |
| oRG | RG | Outer Radial Glia |
| tRG | RG | Truncated Radial Glia |
| RG-div2 | RG-div | Dividing Radial Glia (G2/M-phase) |
| RG-div1 | RG-div | Dividing Radial Glia (S-phase) |
| RG-early | RG-early | Early Radial Glia |

Table S2: Nowakowski's cell type groupings with abbreviations used in the text and figures.

### Supplemental Figures

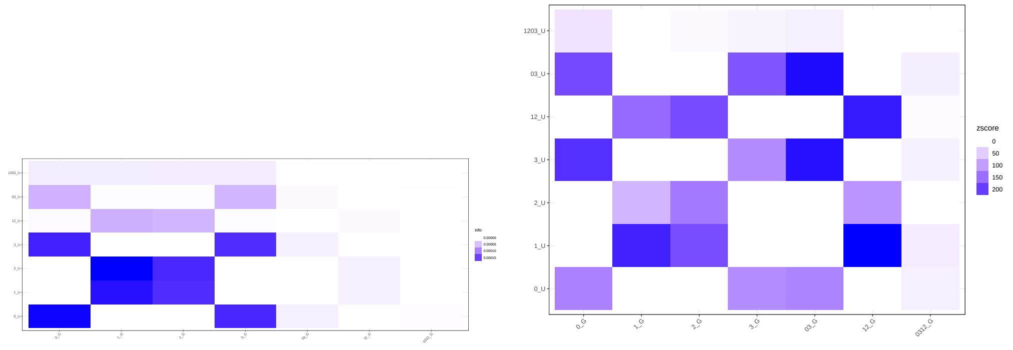


(a) Influence matrix

(b) Zscore matrix

Figure S1: Influence (a) and Z-score (b) matrices between two sets of cell type labels in one dataset from Simulation-easy scenario. This simulation has identical cell types in two scRNASeq datasets. We used suffix 'U' to denote the cell types in the first data set and 'G' for the second. Leaf nodes in general have larger influence scores even for mapping the ancestor cell type. Thus it is hard to tell if a cell type should be mapped to leaves or internal nodes in the other hierarchy by influence scores. However, Z-score can map the cell types to the correct level as the ancestor node has a larger Z-score for mapping the corresponding ancestor node in the other dataset. Some of the leaf cell types also have relatively large Z-scores for mapping to their ancestor cell type.

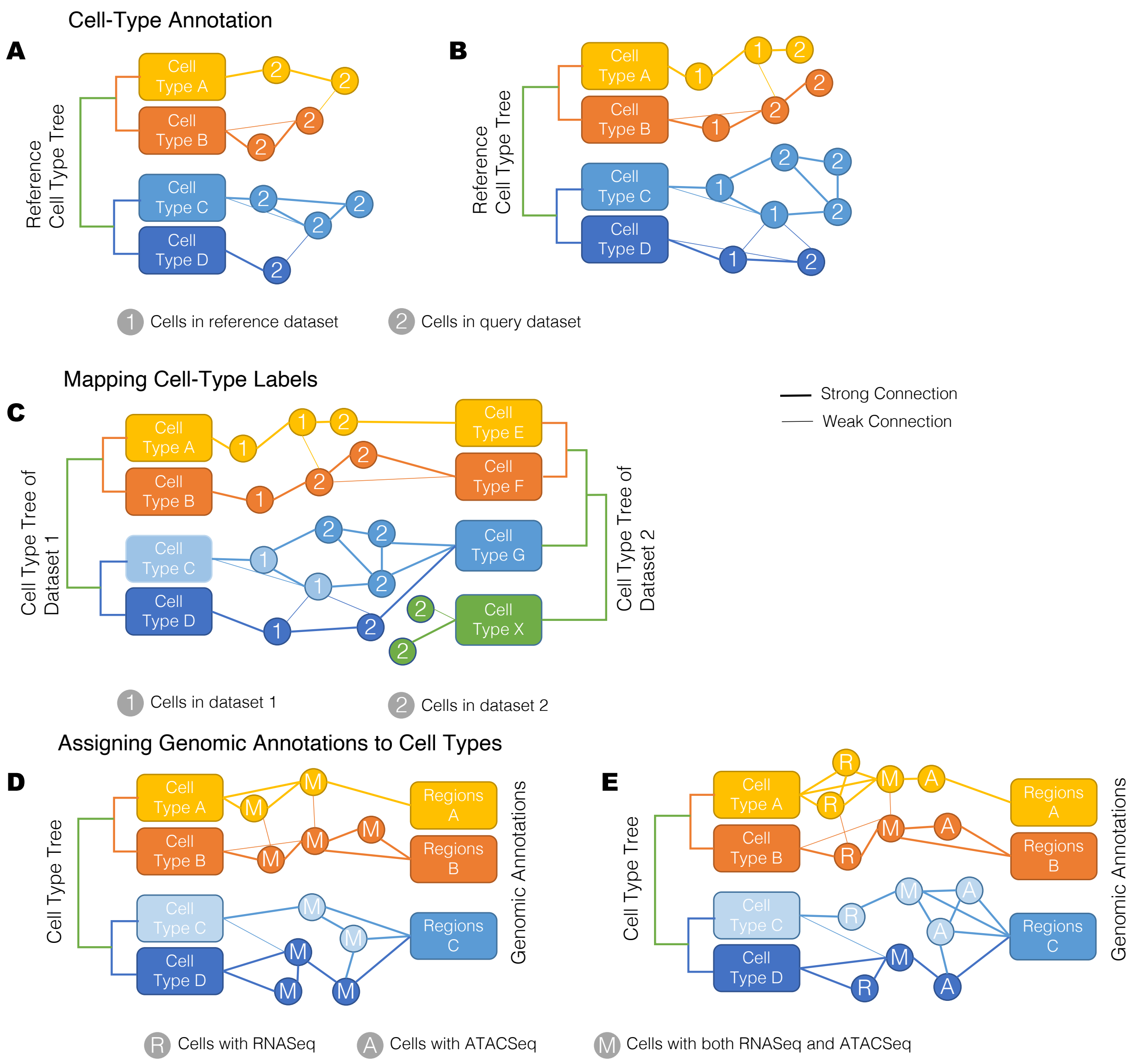


Figure S2: Inputs, outputs, and graphs for different applications of CellWalker2. (A-B) Examples of models for cell type annotation. Input: reference cell type labels with optional hierarchical structure and cells with RNA-Seq or ATAC-Seq data. The edge weight is specified based on the assay. Cell type label is unknown; cells are colored for illustration only. Output: influence matrix and Z-scores for mapping cells to labels. (A) Using cells from only the query dataset. (B) Using cells from both reference and query dataset. (C) Model for mapping cell type labels between two or more ontologies. Input: cell type labels from each dataset with optional hierarchical structure and cells with RNA-Seq or ATAC-Seq data. In this case, cell type E $\approx$ A, F $\approx$ B, G $\approx$ CD (the parent of C and D) and X is not seen in dataset 1 so it is not mapped to any cell type. The correspondence between labels in the two datasets is unknown; nodes are colored for illustration only. Output: influence matrix and Z-scores for mapping cell type labels between datasets. (D-E) Example models for mapping annotations to cell types. Input: cell type labels with optional hierarchical structure, annotations with genome coordinates, and cells with multiomic, scRNA-seq, or scATAC-seq data. The cell types in which the annotations are active are unknown; nodes are colored for illustration only. In these examples, annotation sets A and B are assigned to cell types A and B, respectively. Annotation set C is activated in cells from both cell type C and D, so it would be mapped to their parent cell type (CD). (D) Using cells with multiomic data. Annotations are connected to cells with multiomic data using the ATAC-Seq channel, while cell type labels are connected via the RNA-Seq channel. (E) Integrating cells with multiomic data, only RNA, or only ATAC. Multiomic cells serve as bridge to connect the cells with only one type of data.

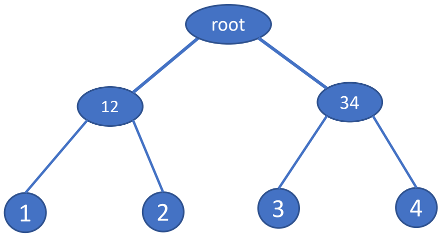


(a) cell type tree

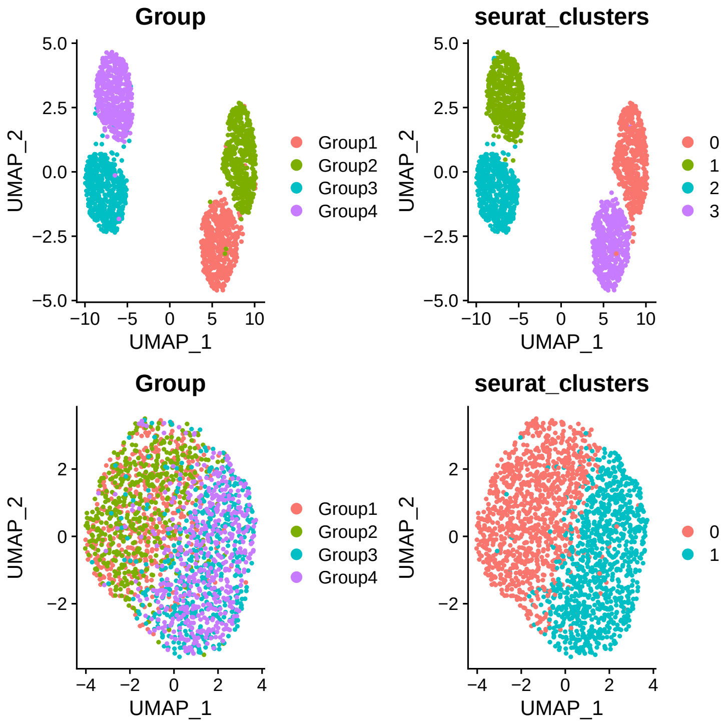


(b) UMAP for medium scenario

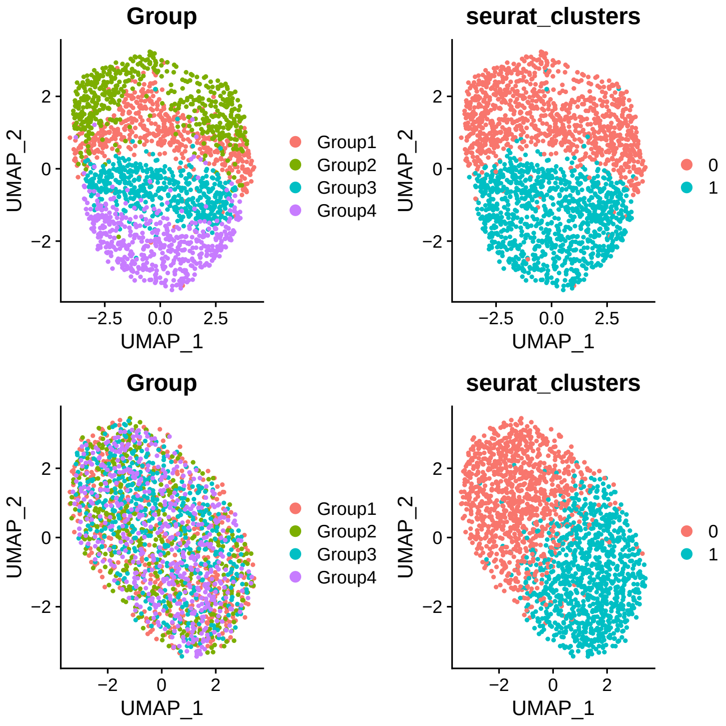


(c) UMAP for hard scenario

Figure S3: Cell type tree and UMAPs for cell annotation simulations. (a) Cell type tree for all simulation scenarios. (b) and (c) UMAP of the simulated data from medium and hard scenarios, respectively. Top and bottom: simulated reference and query datasets. Left panel is colored by true cell type and right is colored by Seurat clusters. In the medium scenario, the query dataset has more dropouts than does the reference dataset, and hence only two cell subclasses are visible on UMAPs generated by Seurat in the query dataset. In the hard scenario, we added more dropouts in reference datasets and cell types are less distinguishable even for the reference data.

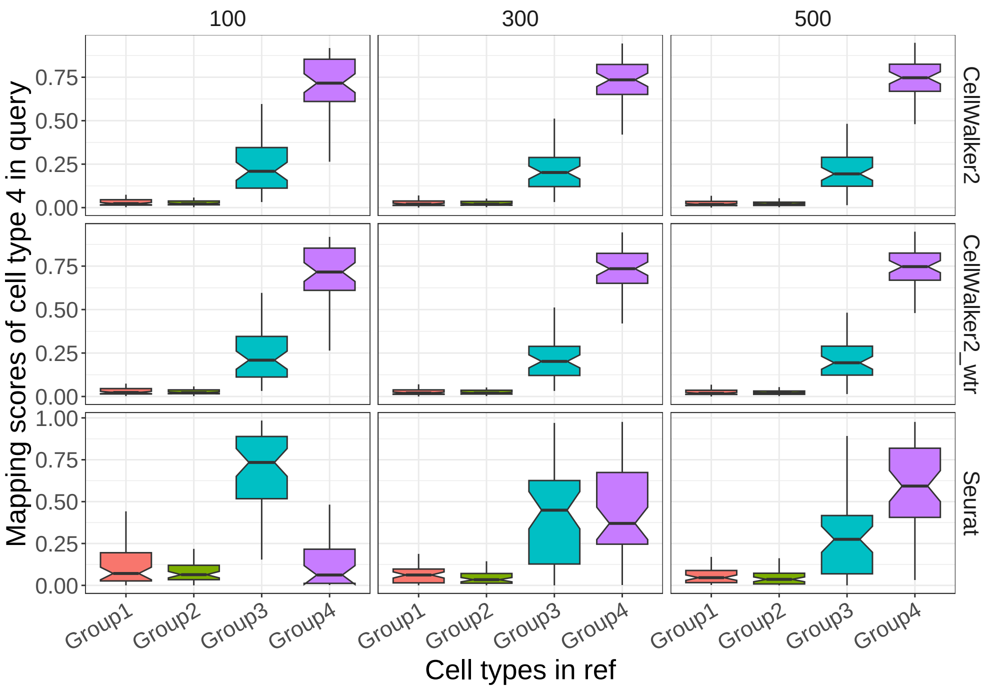


Figure S4: Comparison of performance of CellWalker2 and Seurat for cell-to-label mapping when varying the number of cells of cell type 4 in the reference dataset. The boxplots show the percentage of cells of cell type 4 in the query data set that is mapped to different reference cell types. Columns: number of cells from cell type 4 (range: 100 to 500). Rows: methods, in which 'CellWalker2_wtr' is running CellWalker2 with the tree structure of reference cell types and 'CellWalker2' is without the tree structure.

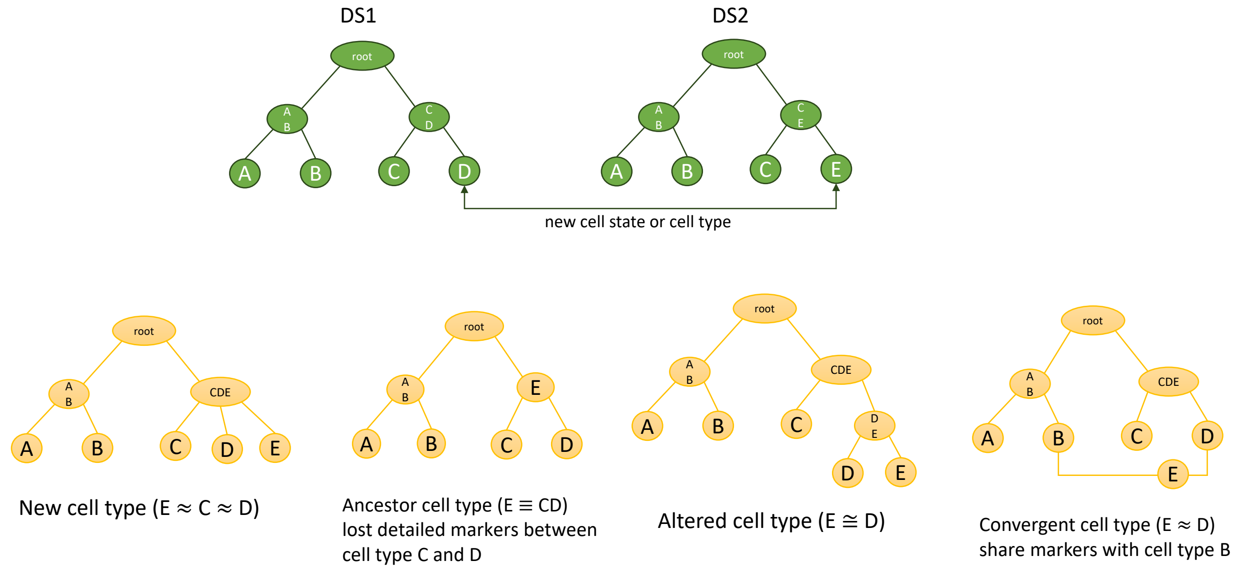


Figure S5: Simulation scenarios for mapping cell types between two hierarchical ontologies. Top: both datasets have four cell types that form a balanced tree. But cell type E in dataset2 (DS2) is a new cell state or cell type compared to DS1. Bottom: four simulation scenarios where E has a different hierarchical relationship with the other cell types. We simulated some marker genes for every split of the cell type tree (see STAR Methods).

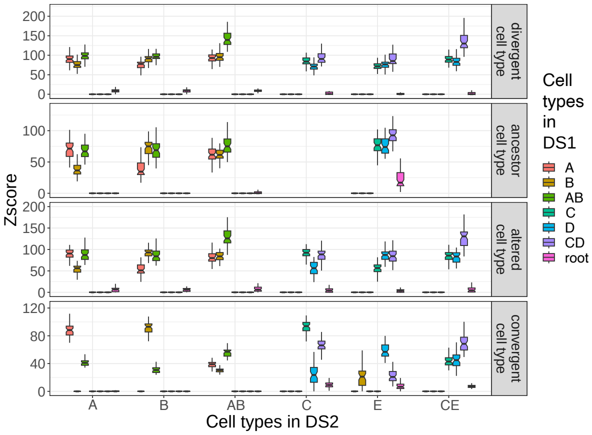


(a) CellWalker2

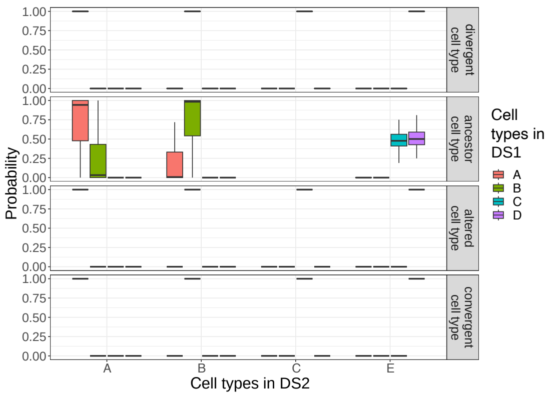


(b) MARS

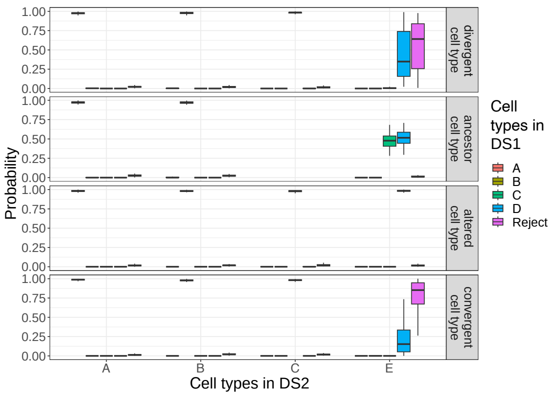


(c) treeArches

Figure S6: Comparison of performance of CellWalker2, MARS, and treeArches for mapping the cell type ontology in dataset 2 (DS2) to the one in dataset 1 (DS1). Four simulation scenarios with different placements of cell type E in DS2 are shown. For both the 'Ancestor' and 'Divergent' cell type cases, CellWalker2 assigned a high Z-score for mapping cell type E to cell type CD, but in the former case, the Z-score to the root node was also large as CD connects to the root on the cell type tree; for the 'Convergent cell type' case, CellWalker2 assigned a lower Z-score to the cell type CD but a larger Z-score to cell type B. MARS assigned a probability near 1 to cell type D in all three cases except 'Ancestor cell type'. treeArches could not distinguish the 'Divergent' and 'Convergent' cell type cases, as it fails to detect the similarity to cell type B that shares features with E.

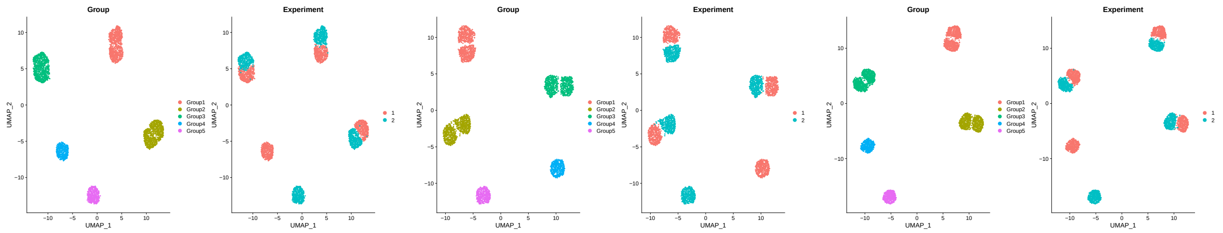


(a) UMAP: low similarity (b) UMAP: medium similarity (c) UMAP: high similarity

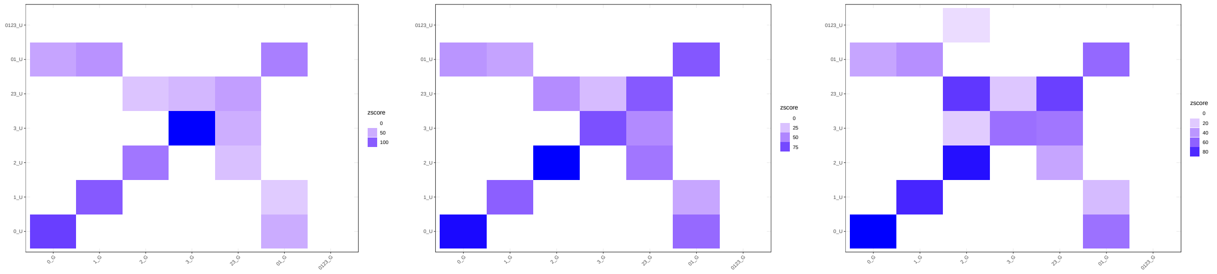


(d) Z-score matrix: low similarity (e) Z-score matrix: medium similarity (f) Z-score matrix: high similarity

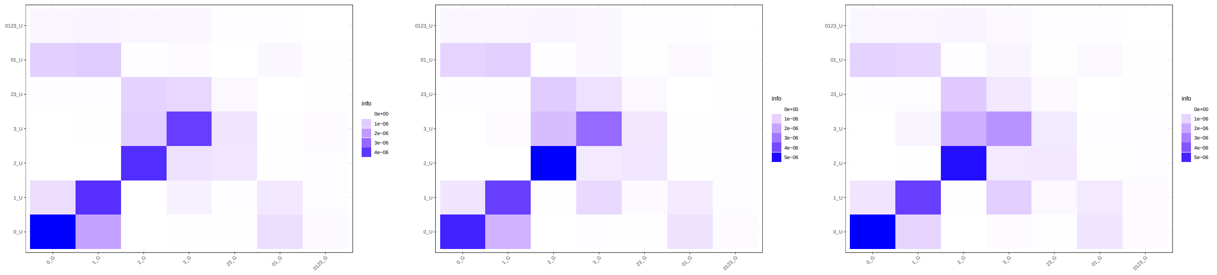


(g) Influence matrix: low similarity (h) Influence matrix: medium similarity (i) Influence matrix: high similarity

Figure S7: UMAP of the simulated data, influence matrix, and Z-score matrix from simulations with a ‘convergent cell type’ scenario in which one of the cell types in the second dataset has features of two cell types in the first dataset. For each row, figures from left to right show the result for scenarios with increasing similarity between cell type 4 (‘3_G’) and cell type {0,1} lineage. The UMAPs contain cells from both datasets, and the left figure is colored by true cell types, while the right one is colored by dataset.

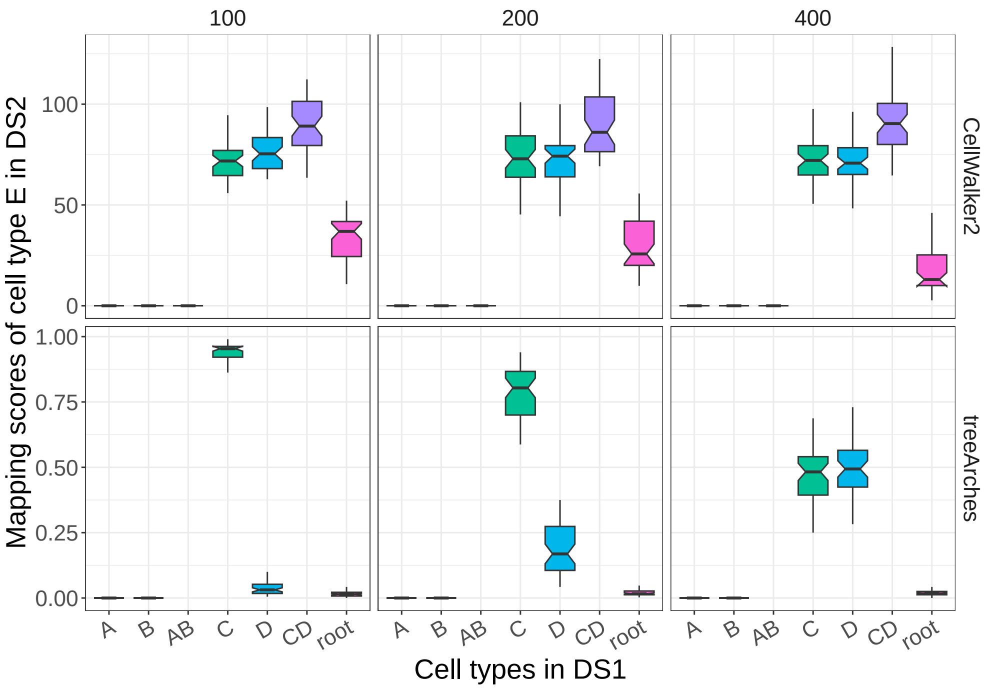


Figure S8: Comparison of CellWalker2 and treeArches for mapping cell types when varying the number of D cells in dataset 1. The boxplots show mapping Z-scores for cell type E in dataset 2 to different cell types in dataset 1. Different columns are different numbers of D cells, ranging from 100 to 400.

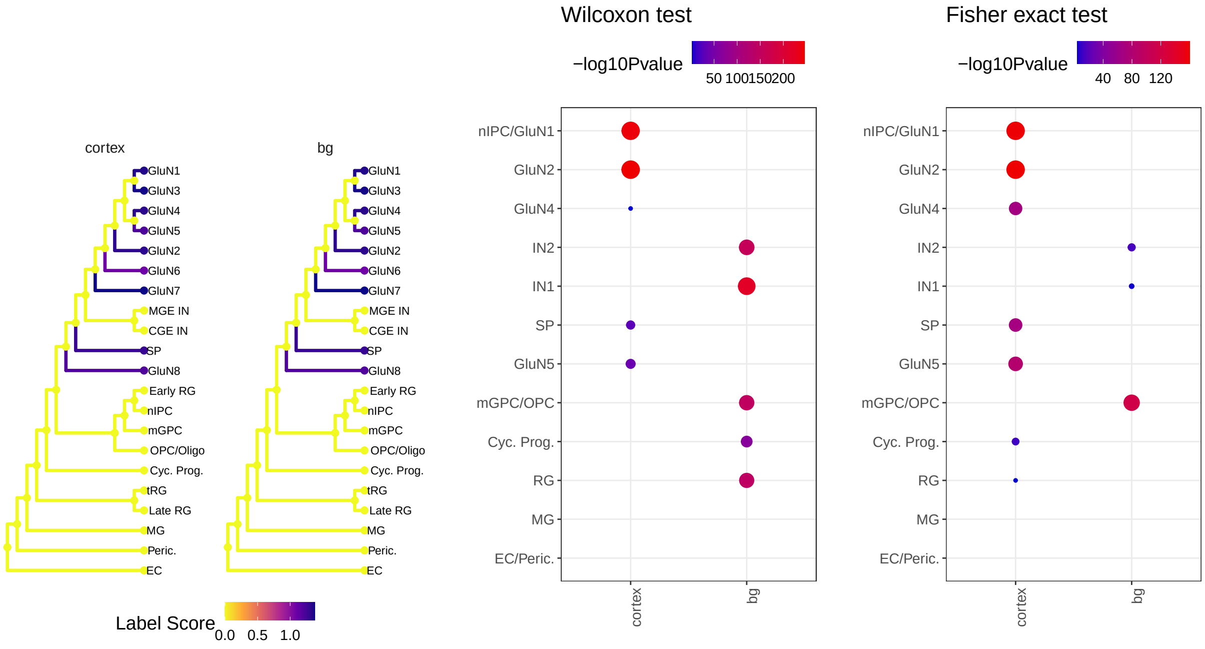


(a) labelScore (b) Wilcox Test (c) Fisher's exact Test

Figure S9: Comparison of different methods mapping basal ganalia versus cortex specific pREs [1] to different cell types in Trevino et al.. (a) labelScore from CellWalker mapping region specific pREs to cell type hierarchy in [2] using both multiomic and RNASeq data. (b) Mapping region specific pREs to cell types identified from transcriptomic profile in the multiomic data using the Wilcoxon test. We compared the distribution of edge weights to basal ganglia versus cortex specific pREs for each cell type. (c) Mapping region specific pREs to cell types identified from transcriptomic profile in the multiomic data using Fisher's exact test. We identified differentially accessible regions (DARs) for each cell type first, and then tested if region specific pREs are over-represented in these DARs. Errors may be due to difficulties calling DARs using scATAC-seq data.

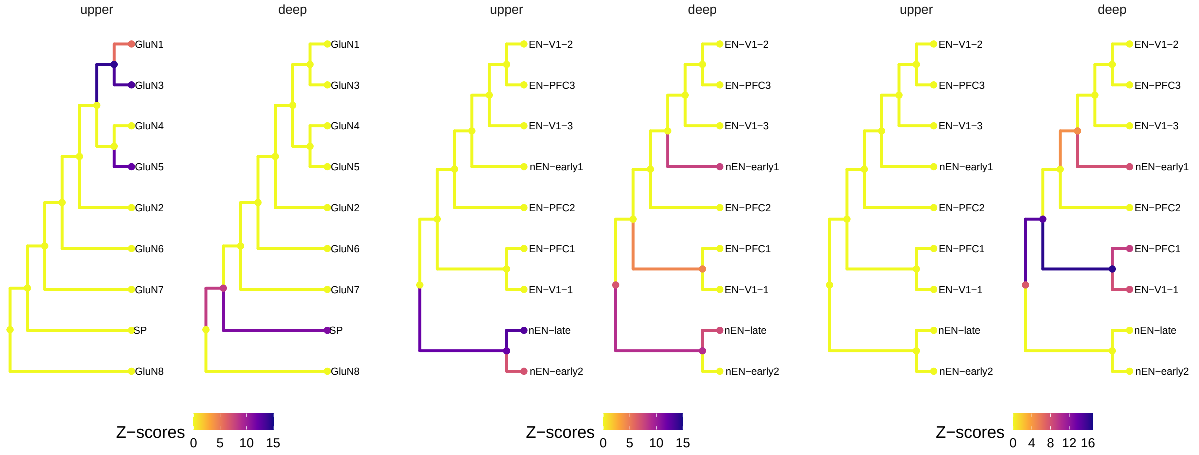


(a) pcw21 (b) pcw21 (c) pcw16

Figure S10: Mapping layer-specific pREs to different cell type lineages by integrating ATACSeq from different developmental ages and using different cell type ontologies. As neurons from deep laminae layers develop at an earlier age than those from upper layers, we incorporated either scATAC-Seq cells from a later stage (week 21) or an earlier stage (week 16) into the cell graph. (a) Z-scores mapping upper vs deep layer specific pREs to cell type hierarchy in [2]. (b) and (c) Z-scores mapping upper vs deep layer specific pREs onto cell type hierarchy of excitatory neurons in [3], which annotates more fine-grained cell types. scATACSeq from post-conception week (pcw) 21 and pcw 16 was used, respectively. As expected, pREs from the upper layer have higher chromatin accessibility in late-born newborn excitatory neurons (nEN-late) and this increased accessibility appears in week 21. On the other hand, pREs from deep layer have higher chromatin accessibility in early-born deep layer/subplate excitatory neurons (EN-PFC-1 and EN-V1-1) as well as early-born newborn excitatory neurons (nEN-early1), and the cell type specificity is weakened at week 21 with smaller Z-scores.

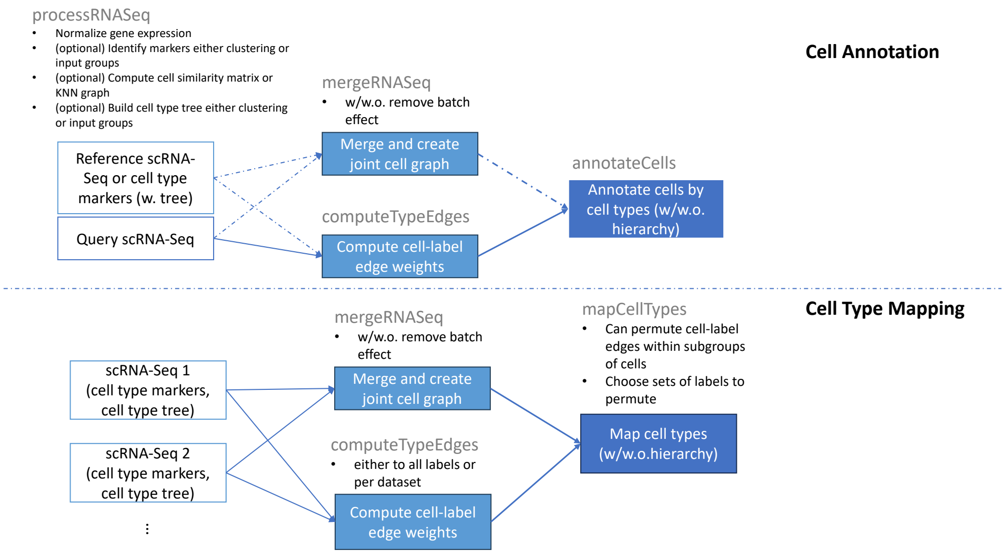


(a)

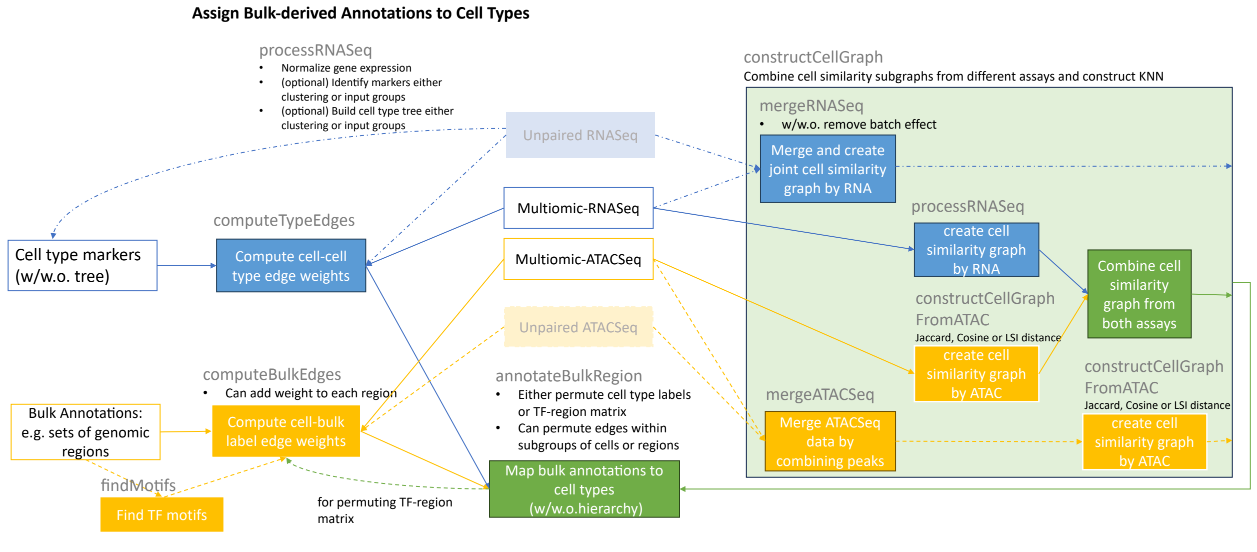


(b)

Figure S11: CellWalker2 pipelines in CellWalkR package. (a) Cell annotation and cell type mapping pipeline. (b) Mapping bulk-derived annotations to cell types pipeline. Solid line, required steps; dashed line, optional steps. Box with no filling, required input data; Box with shaded filling, optional input data. CellWalker2 functions for RNASeq data, blue box; CellWalker2 functions for ATACSeq data, orange box; CellWalker2 functions for both assays, green box. The function name is labeled above each box.

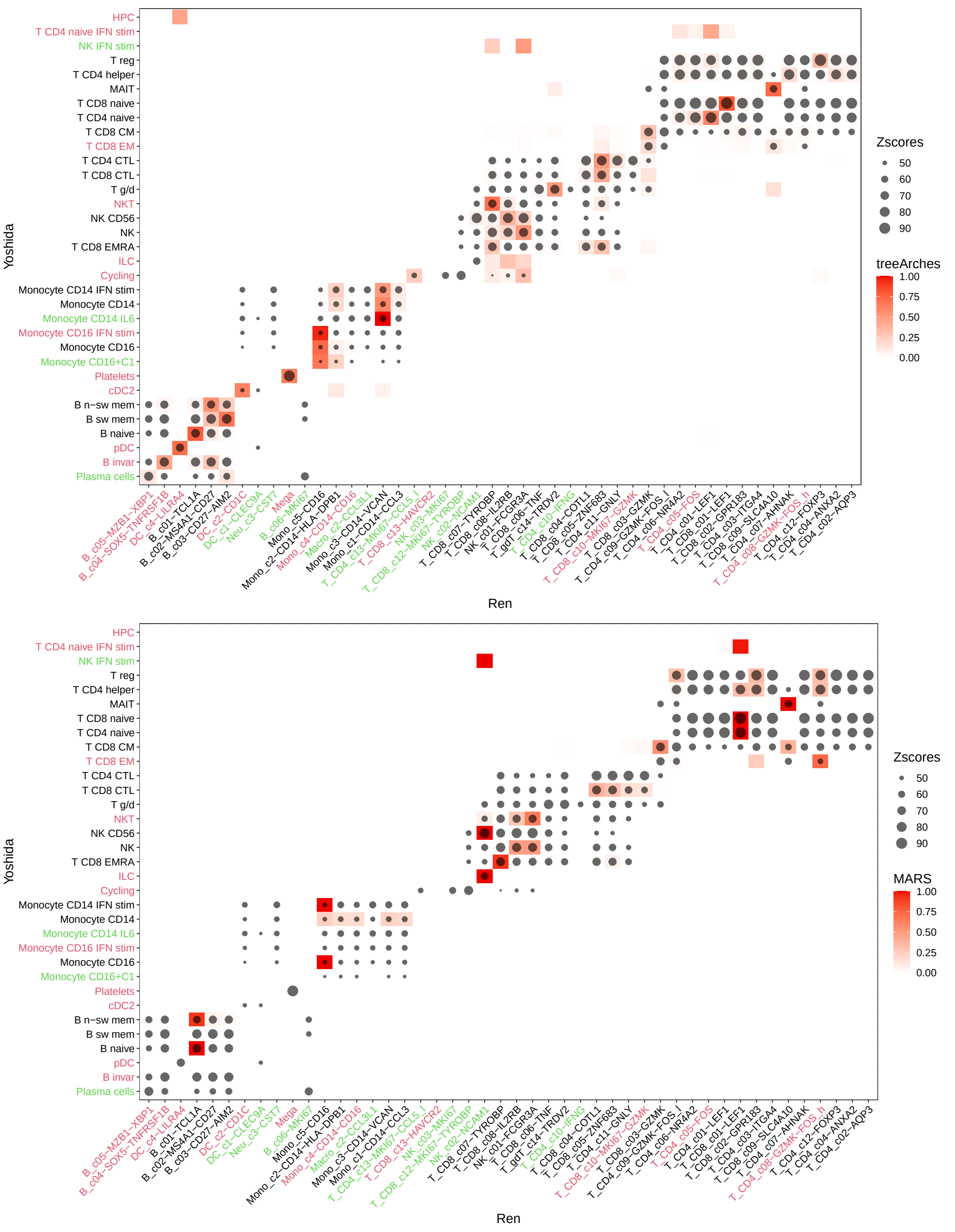


Figure S12: Mapping cell types between Yoshida et al.[4] and Ren et al.[5] datasets with CellWalker2, treeArches, and MARS. Each column is a cell type from Ren et al. and each row is a cell type from Yoshida et al.. Ren's cell types are ordered by the cell type tree. Yoshida's cell types are ordered by the matching cell types of Ren's. Only tips are shown. The size of the dot represents Z-score. The color represents mapping probabilities by treeArches (left) and MARS (right). MARS made many errors, such as mapping all naive T cells (CD4/CD8/IFN stimulated CD4) to naive CD8 T cells (T CD8 c01-LEF1). treeArches was biased towards more prevalent cell types, e.g., treeArches maps CD4 cytotoxic T lymphocytes (T CD4 CTL) to T CD8 c05-ZNF683 (CD8 cytotoxic T lymphocytes), which is one of the four most prevalent cell types and a common map for treeArches, rather than to the less prevalent T CD4 c11-GNLY, which matches expectations given that both are CD4 cells and GNLY is a marker of cytotoxic T lymphocytes. Z-score $> 47$ are shown (set the threshold to include all treeArches hits with probability $> 0.5$). Green: cell type $< 0.1\%$, Red: cell type $< 1\%$. CM, central memory; CTL, cytotoxic T lymphocytes; EM, effector memory; EMRA, effector memory re-expressing CD45RA; invar, invariant; n-sw, non-switched; sw, switched; IFN-stim: interferon-stimulated subpopulations. For Ren et al., cell types are named based on marker genes. "high" or "low" labels indicate the relative expression levels and genes without high or low labels are specifically expressed in the cell type.

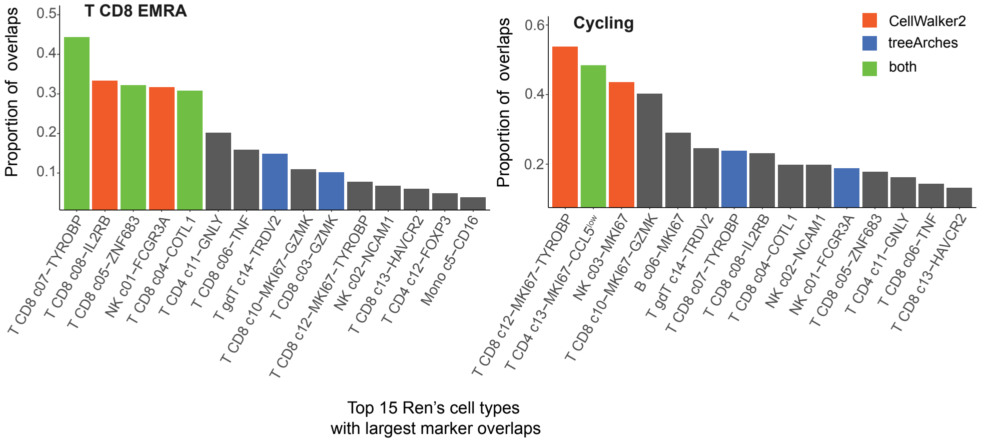


Figure S13: Proportion of positive markers overlapping between T CD8 EMRA (left) and Cycling (right) in Yoshida et al. versus different cell types in Ren et al.. Only the top 15 cell types with the largest overlaps are shown. Orange: Top 5 (T CD8 EMRA) or 3 (Cycling) mapped cell types by CellWalker2; Dark blue: Top mappings by treeArches; Green: Top mappings by both methods.

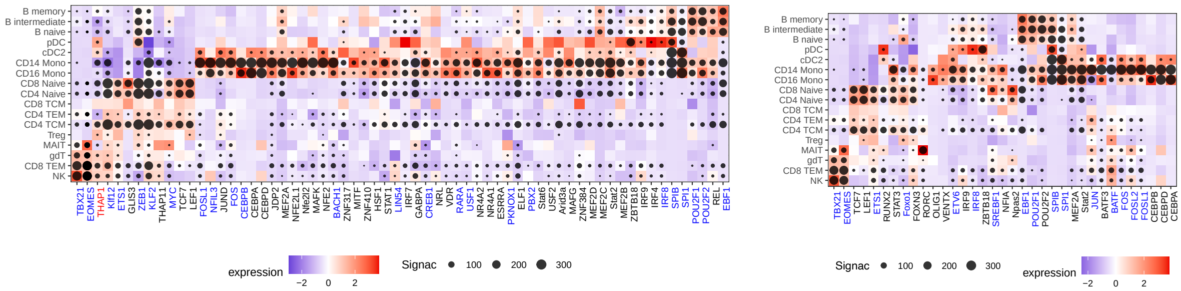


(a) Signac

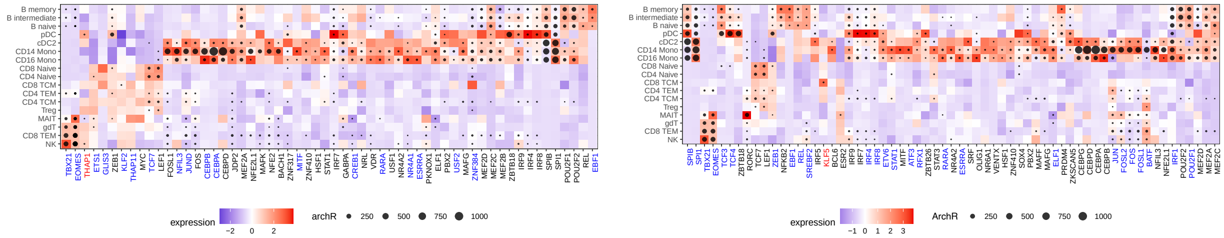


(b) ArchR

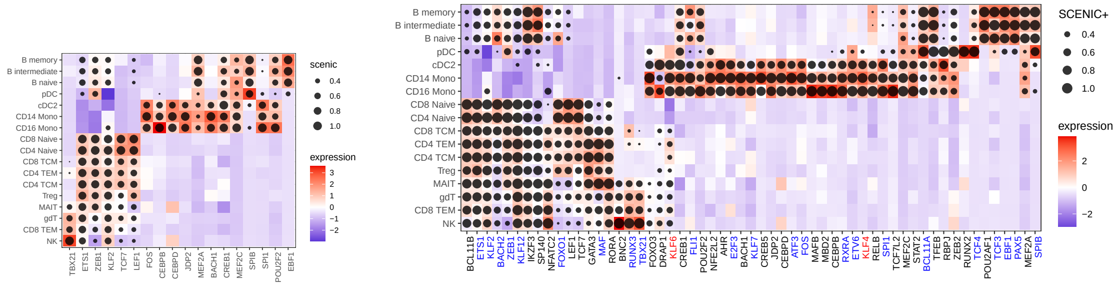


(c) SCENIC+

Figure S14: Cell type-specific transcription factors in PBMC by different methods. Each row is a cell type and each column is a transcription factor (TF). The color represents normalized expression level for each TF across cell types. The size of the dot represents the score by different methods: -log10 P-values for Signac (a) and ArchR (b), AUC for SCENIC+ (c). In each panel, the left figure shows TFs selected by CellWalker2, and the right figure shows TFs selected by one of the other methods. TFs with names in red font are human universal stripe factors, while those in blue font are other stripe factors. As SCENIC+ and CellWalker2 use different TF motif databases, not all TFs selected by CellWalker2 are included in the SCENIC+ analysis, and hence the left heatmap in (c) shows fewer TFs. CellWalker2's result is in Figure 3D. In general, TFs show more cell type specificity with CellWalker2 compared to Signac and ArchR, which show broad enrichment patterns for several TFs that are known to be regulators of specific lineages. For example, *TBX21* and *EOMES* are significant in B cells in addition to T and NK cells for both methods, and the B cell-specific TF *EBF1* is significant in monocytes and T cells for Signac.

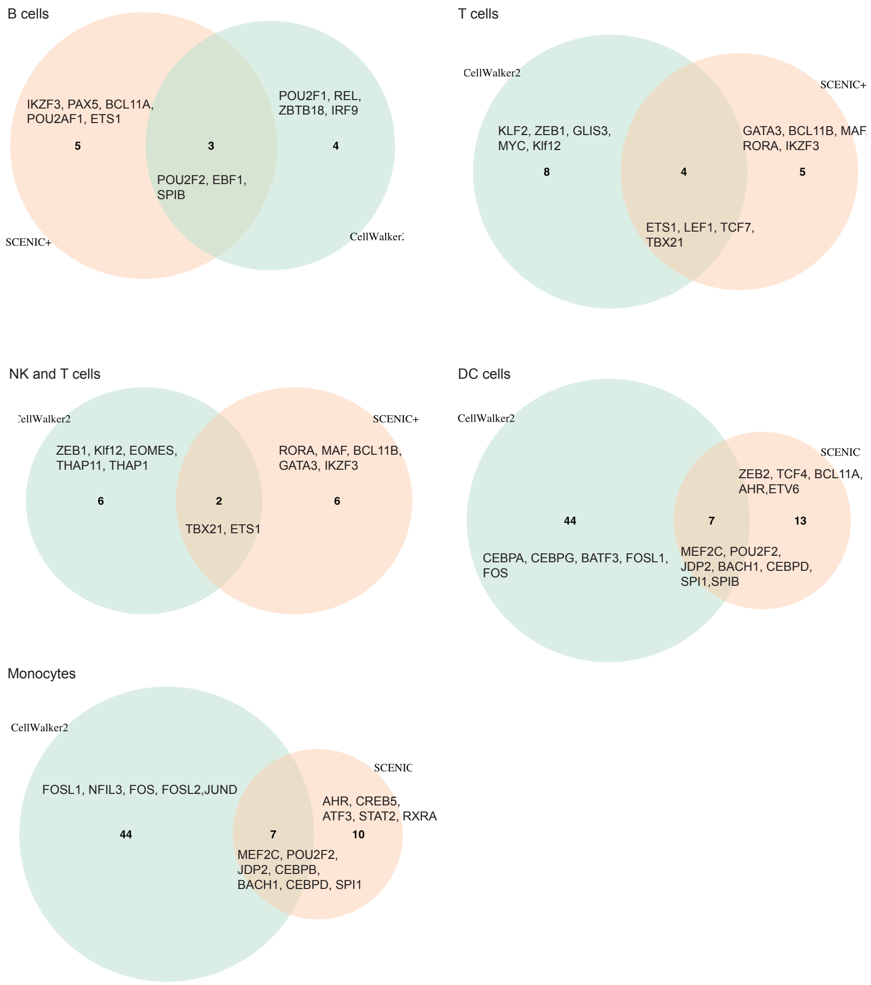


Figure S15: Comparison of transcription factors identified by SCENIC+ and CellWalker2 in major PBMC cell types. The number of TFs uniquely identified and found in common by each method are shown. For cell types with too many uniquely identified, only the top 5 TFs with largest Z-scores or AUC score are named.

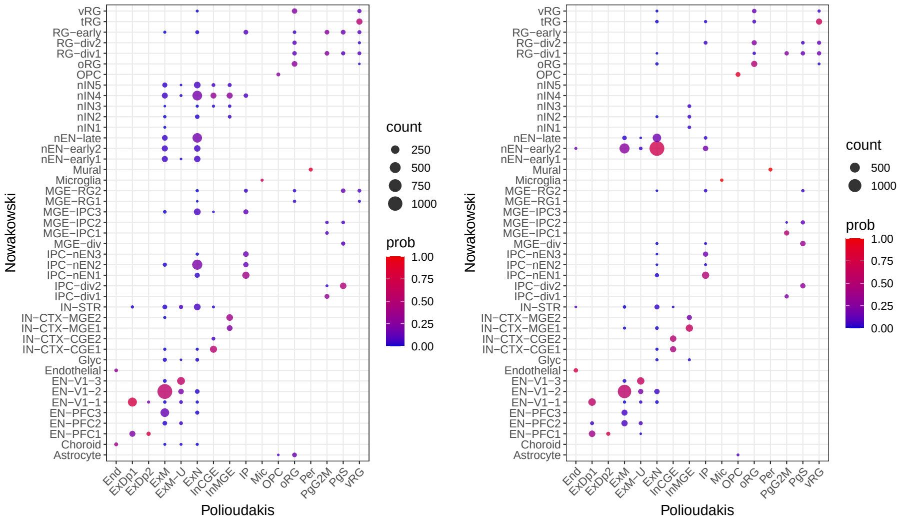


(a) CellWalker2 vs Seurat

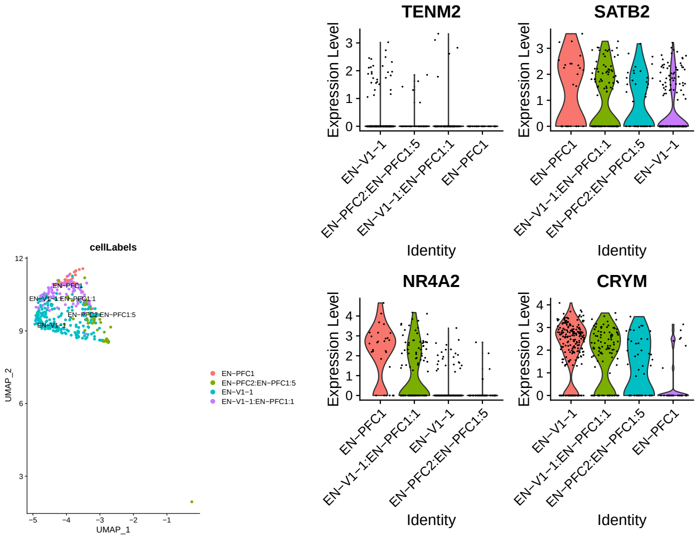


(b) ExDp1

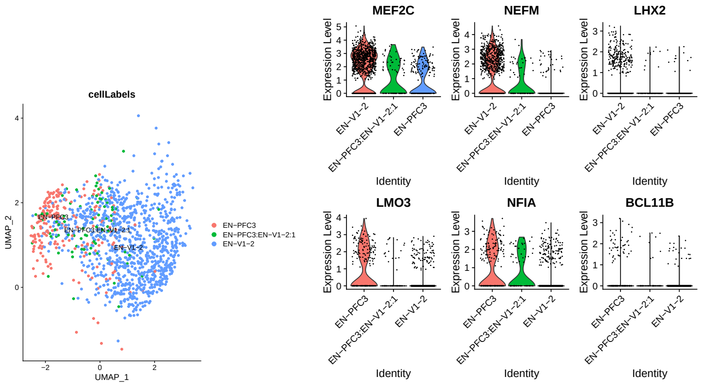


(c) ExM

Figure S16: (a) Mapping cells to cell types using CellWalker2 and Seurat using human developing cortex scRNA-Seq data. We annotated cells in Polioudakis et al.'s dataset (query) with the cell types from Nowakowski et al. (reference). The color of each dot represents the probability of mapping each cell type in query dataset to reference cell type labels, and the size of each dot represents the number of cells mapped (mappings with at least 5 cells are shown). (b-c) UMAP and expression level of markers of ExDp1 (b) and ExM (c). UMAP is computed using all cells from Polioudakis et al. but only a random subset of cells are plotted.

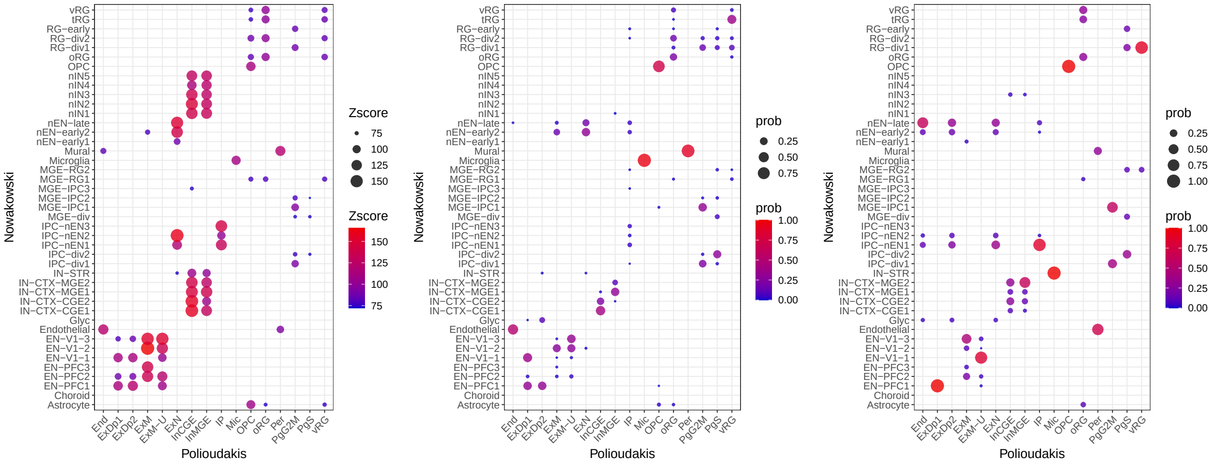


Figure S17: Comparison of mapping cell types between Polioudakis et al. and Nowakowski et al. dataset. Z-scores from CellWalker2 (left), probabilities from treeArches (middle), and probabilities from MARS (right). Each column is a cell type from Polioudakis et al., and each row is a cell type from Nowakowski et al.. As treeArches and MARS do not consider hierarchical structure of the cell types, we only include terminal cell types as well for CellWalker2 for this figure. MARS made errors for several rare, non-neuronal cell types including pericytes (Per; mapped to endothelial cells) and microglia (Mic; mapped to Striatal neurons (IN-STR)). Only Z-scores > 70 and probabilities > 0.003 are shown.

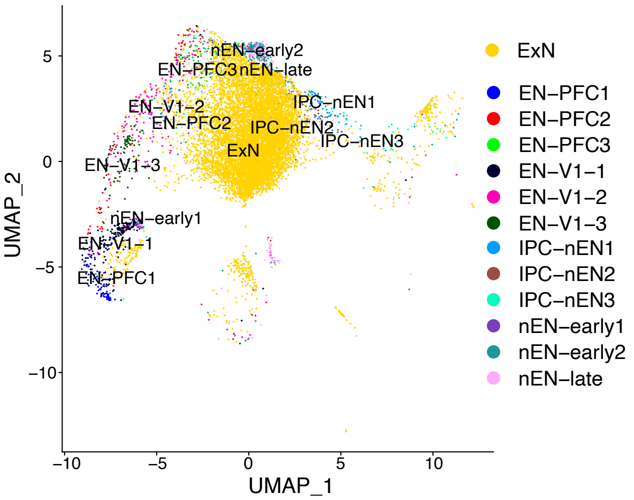


Figure S18: ExN shows proximity to the cell types mapped by CellWalker2. In a UMAP plot, ExN is close to IPC-nEN2, nEN-late, and nEN-early2. The UMAP is drawn using integrated data including ExN from Polioudakis et al. and excitatory neuron related cell types from Nowakowski et al..

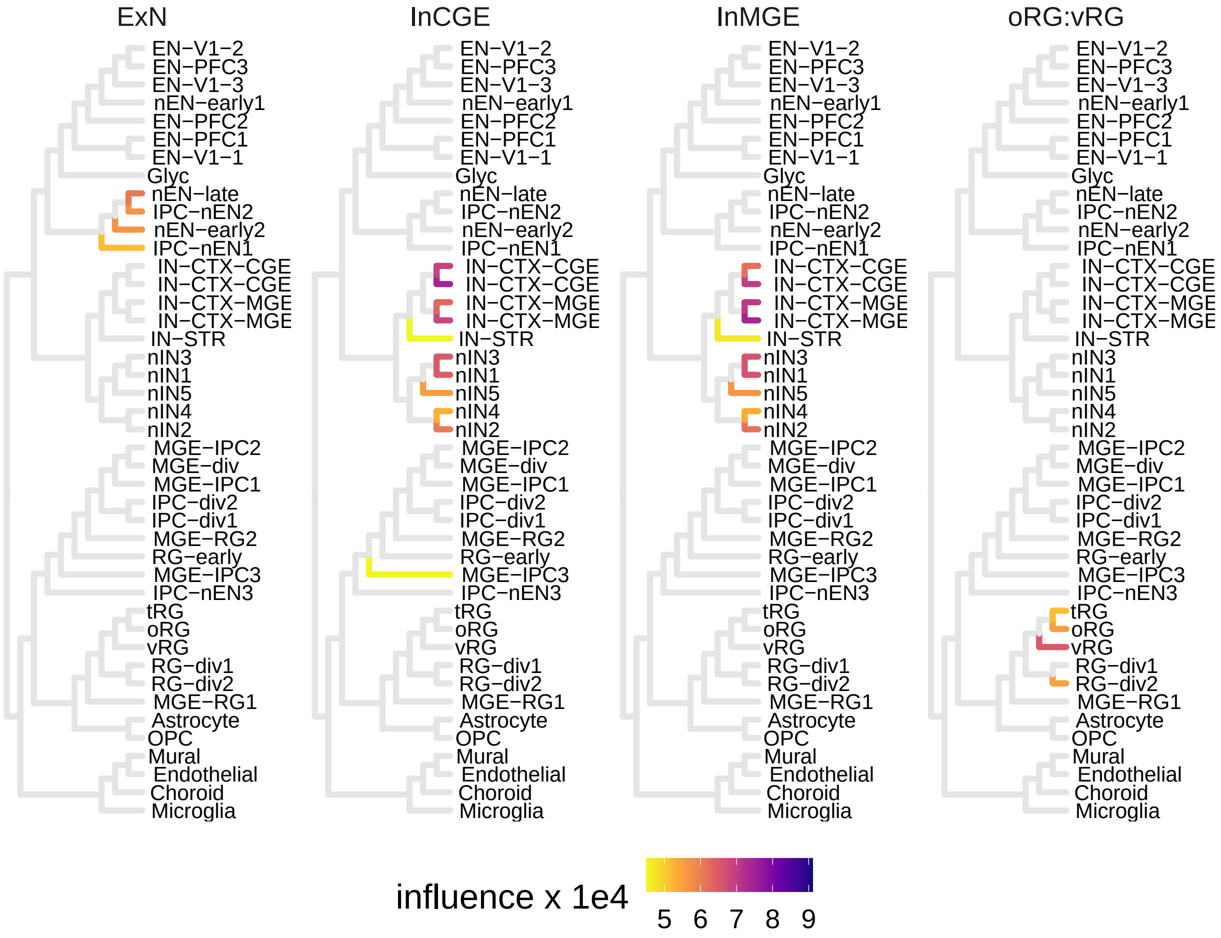


Figure S19: Mapping ExN, InCGE, InMGE and the ancestral cell type oRG:vRG in Polioudakis et al. onto the cell type hierarchy in Nowakowski et al. by influence scores ($> 1e-4$ are shown).

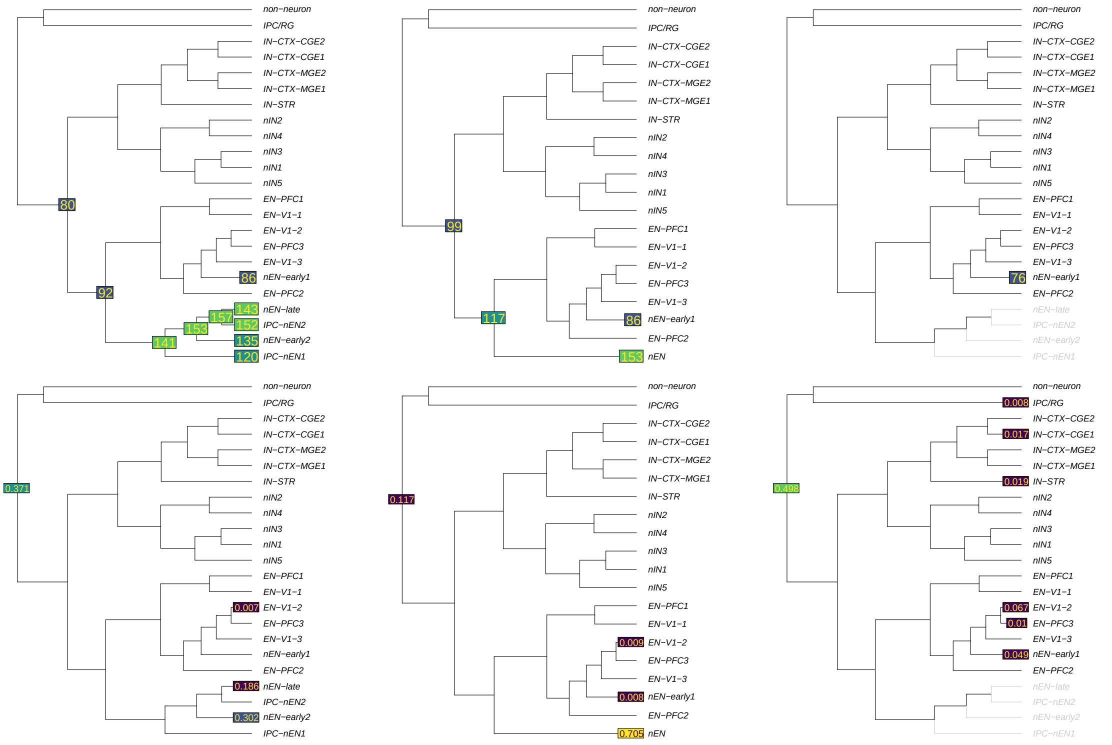


(a) (b) (c)

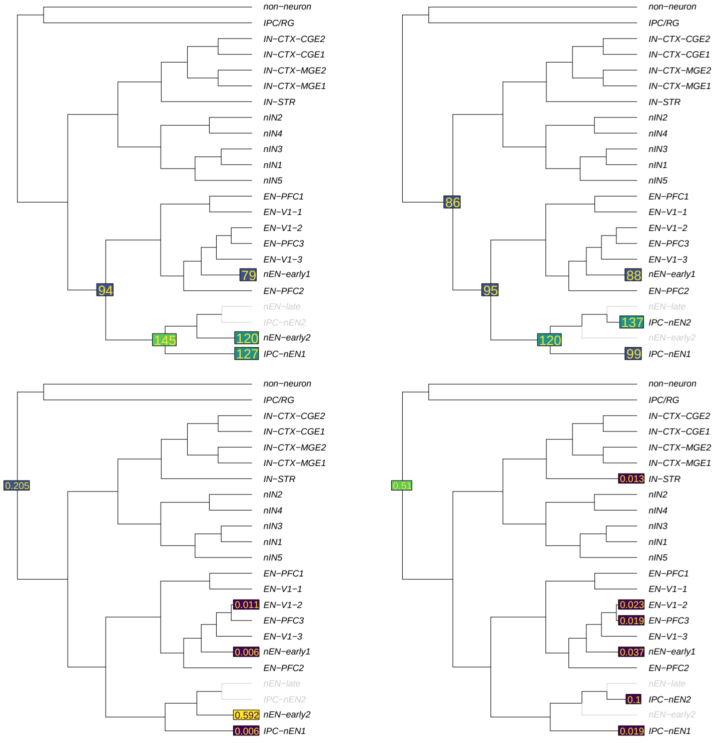


(d) (e)

Figure S20: Mapping migrating excitatory (ExN) in Polioudakis et al. onto the hierarchical cell type ontology of Nowakowski et al.. treeArches and CellWalker2's cell type mapping results are shown across scenarios in which some of the cells and subtypes on the cell type tree are removed or subtypes are combined. (a) Original (b) Combine all cell types in the newborn excitatory neurons subtree. (c) Remove all cell types in the newborn excitatory neurons subtree. (d) Remove nEN-late and IPC-nEN2. (e) Remove nEN-late and nEN-early2. Top: Z-scores from CellWalker2; Bottom: mapping probabilities from treeArches. The rejection probabilities for treeArches are 0.10, 0.14, 0.30, 0.15 and 0.25 for these five scenarios respectively. Z-score cutoff is 75 and treeArches cutoff is 0.005. Only the top 10 nodes and above the cutoff values are shown.

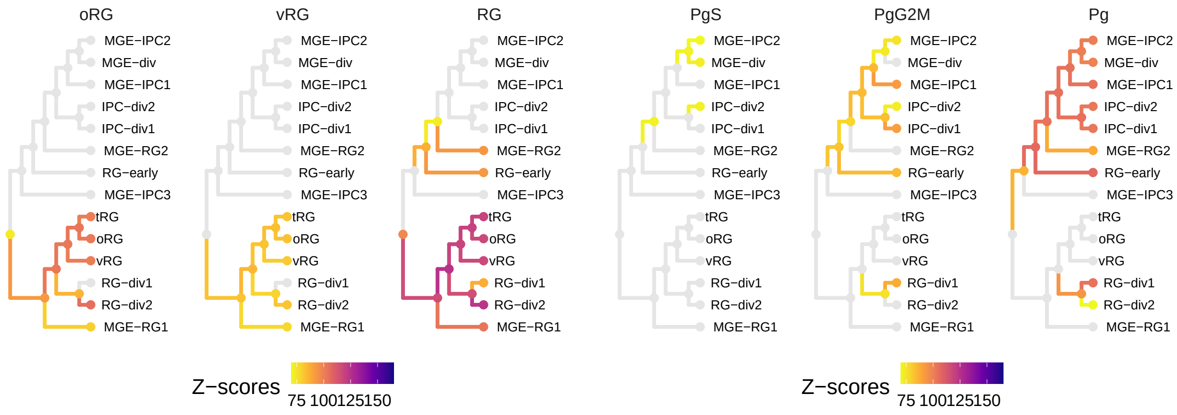


Figure S21: Mapping radial glias and cycling progenitors in [6] onto the cell type hierarchy in [3] by Z-scores ($> 70$ are shown). RG represents the parent node of oRG and vRG, and Pg represents the parent node of PgS and PgG2M. oRG and vRG map to the radial glia subtree, but show no one-to-one correspondence with specific cell types. The RG node shows larger Z-scores mapping to the subtree of radial glia, with the largest Z-score at the internal node vRG:RG-div1. Cycling progenitors in G2M phase (PgG2M) maps to dividing radial glia in G2/M-phase (RG-div1), dividing intermediate progenitor cells RG-like (IPC-div1) and MGE Progenitors(MGE-IPC1). Cycling progenitors in S phase (PgS) maps to dividing intermediate progenitor Cells RG-like (IPC-div2) and dividing MGE Progenitors (MGE-div) weakly. The Pg node maps to dividing progenitors, MGE progenitors and early/dividing RG subtree with larger Z-scores, and the internal nodes of this subtree have the top Z-scores (e.g. RG-early:MGE-IPC2 and IPC-div1:MGE-IPC2).

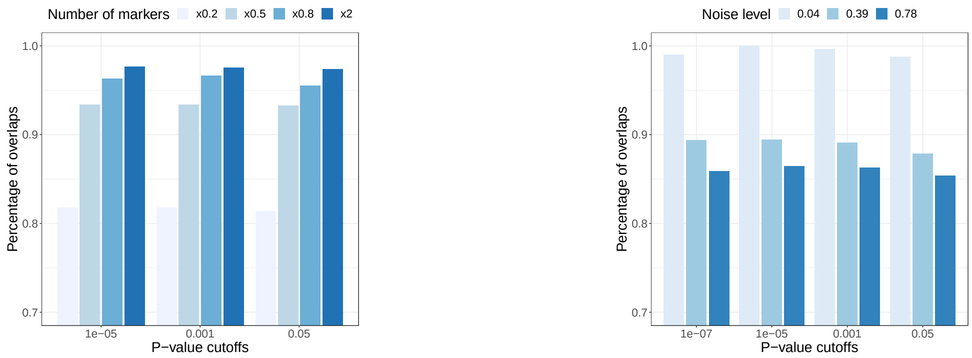


(a) Varying number of markers

(b) Adding noise to gene expression

Figure S22: Robustness of CellWalker2's results for mapping cell types between Polioudakis et al. and Nowakowski et al.. (a) Varying the number of markers of each cell type from top 20%, 50%, 80% and 200% of the original markers in the order of log fold change. (b) Adding different levels of Gaussian noise to each PC coordinate of cells derived from gene expression matrix and reconstructing cell-cell graph. Y-axis: the percentage of significant entries overlaps with original result. X-axis: different cutoffs for P-values (converted from Z-scores).

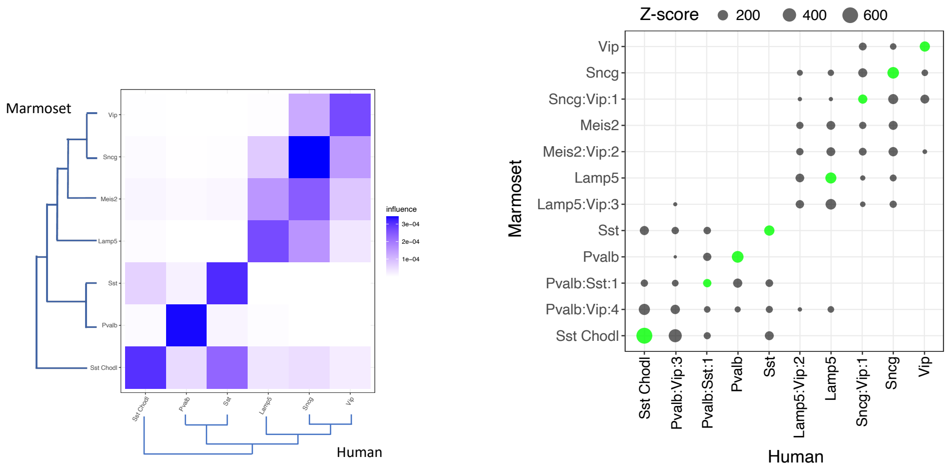


Figure S23: Mapping inhibitory neurons in motor cortex between human and marmoset at the subclass level. CellWalker2's influence scores (left) and Z-scores (right) reflect the corresponding inhibitory neurons subclasses between human and marmoset. Subclasses cluster into two groups based on their Z-scores (Lamp5, Sncg, Vip and Meis2 versus Sst, Pvalb and Sst Chodl), which correspond with CGE- and MGE-derived inhibitory neurons. The only exception is marmoset Meis2 cells showing weak similarity to human Sncg cells but no strong mapping to any human subclass, consistent with the lack of Meis2 cells in human motor cortex. Each row is a marmoset cell subclass and each column is a human cell subclass, including ancestral nodes for Z-scores. The ancestral nodes are labeled with the two tips within its descendants and the depth of the node. The one-to-one relationship of the nodes on the human and marmoset tree is shown as green circle. Only Z-scores $> 3$ are shown.

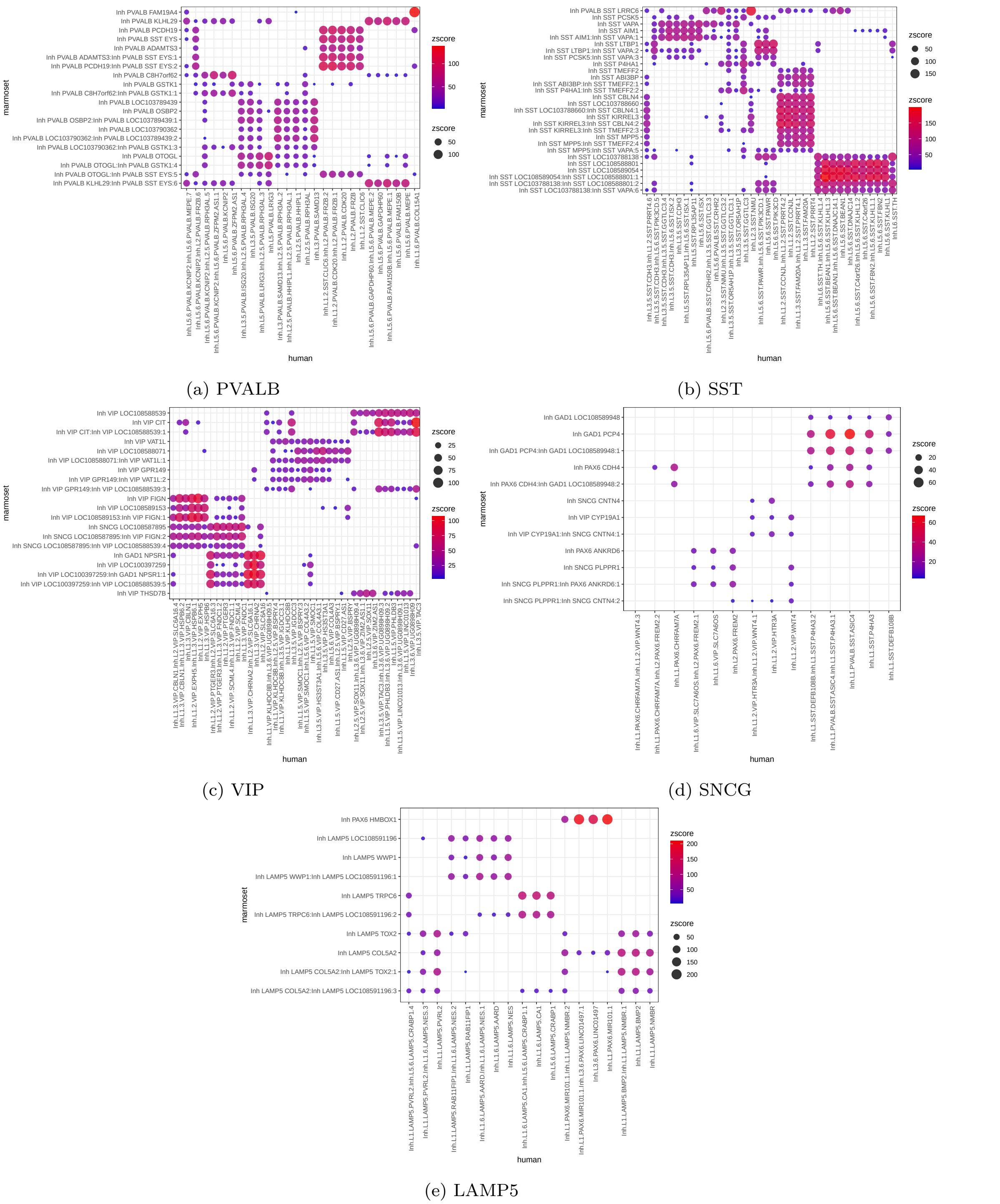

(a) PVALB

(b) SST

(c) VIP

(d) SNCG

(e) LAMP5

Figure S24: Mapping cell types within each GABAergic neuron subclass between human and marmoset using CellWalker2. Each row is a marmoset cell type, and each column is a human cell type. Z-score is represented by the size and color of the dot.

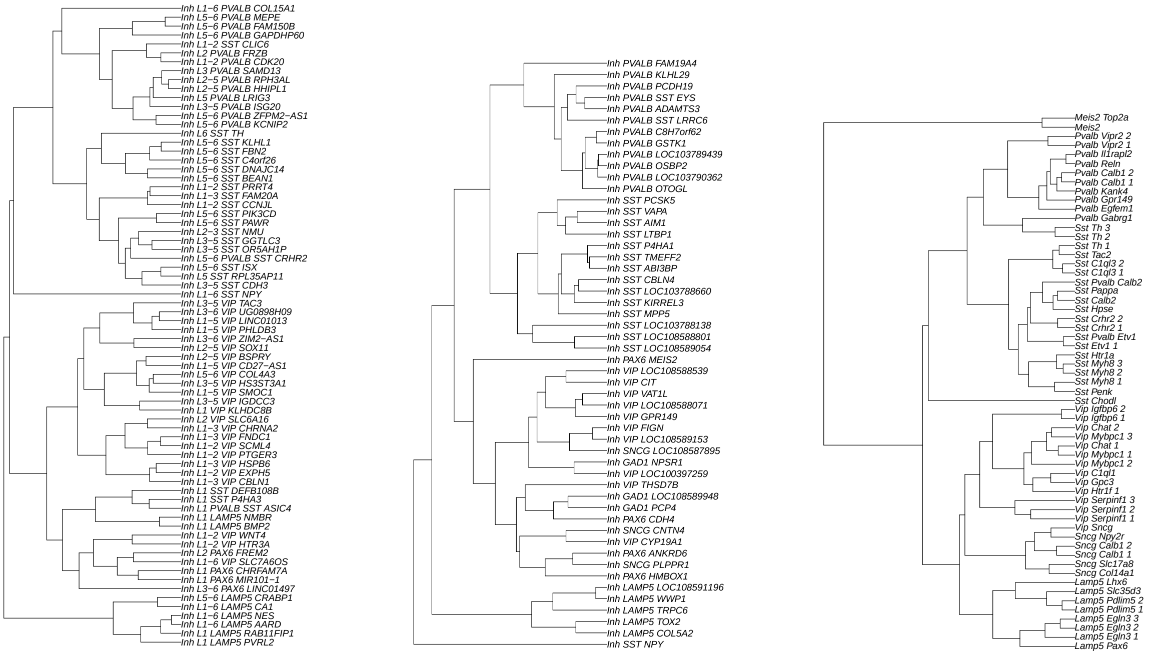


(a) Human (b) Marmoset (c) Mouse

Figure S25: Cell type trees in human, marmoset and mouse. Adapted from [7].

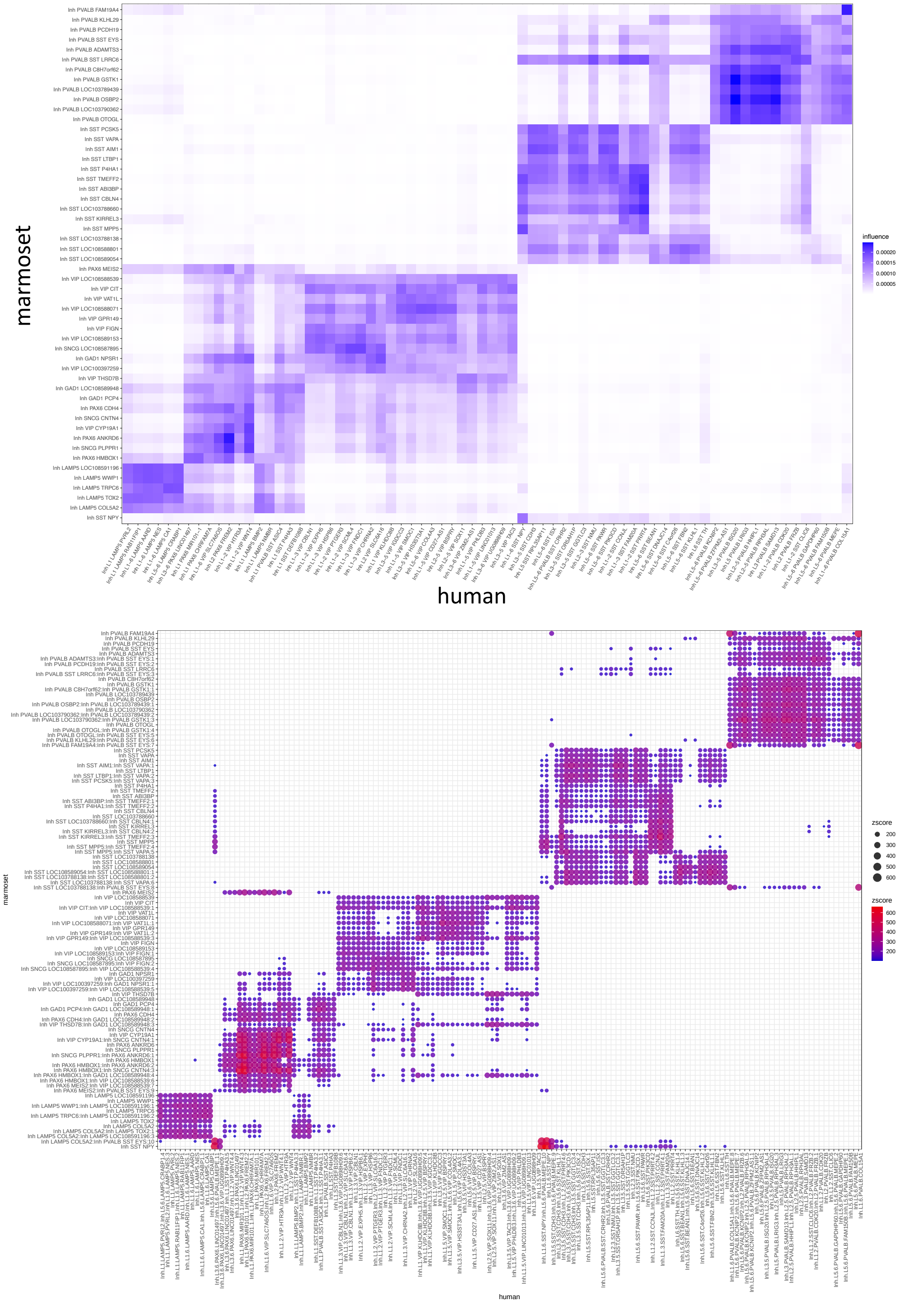


Figure S26: Mapping different cell types of inhibitory neurons in motor cortex between human and marmoset. Left: influence scores from mapping marmoset to human cell types. Right: Z-scores from mapping the human and marmoset cell type trees (including ancestral nodes). The marmoset Sst chodl cell types (Inh SST NPY) is separated from other Sst cell types, while this cell type is grouped together with other Sst cell types in the human ontology. Although human and marmoset Sst chodl cell types have high Z-scores for each other, human Sst chodl cells also map to another subtree of marmoset Sst cell types, in which Inh SST MPP5 is also labeled as Sst chodl in the consensus taxonomy [7]. A marmoset Sst cell type (Inh PVALB SST LRRC6) that express both *PVALB* and *SST* and shows similarity not only to the human Sst subtree but even higher similarity to the Pvalb subtree, consistent with this cell type having features of both Pval and Sst cells. Conversely, on the human cell ontology, a subtree within Sst subclass (Inh L5-6 SST DNAJC14 and Inh L5-6 SST BEAN1) not only maps to the marmoset Sst subtree but also to some Pvalb cell types. Only Z-scores $> 100$ are shown.

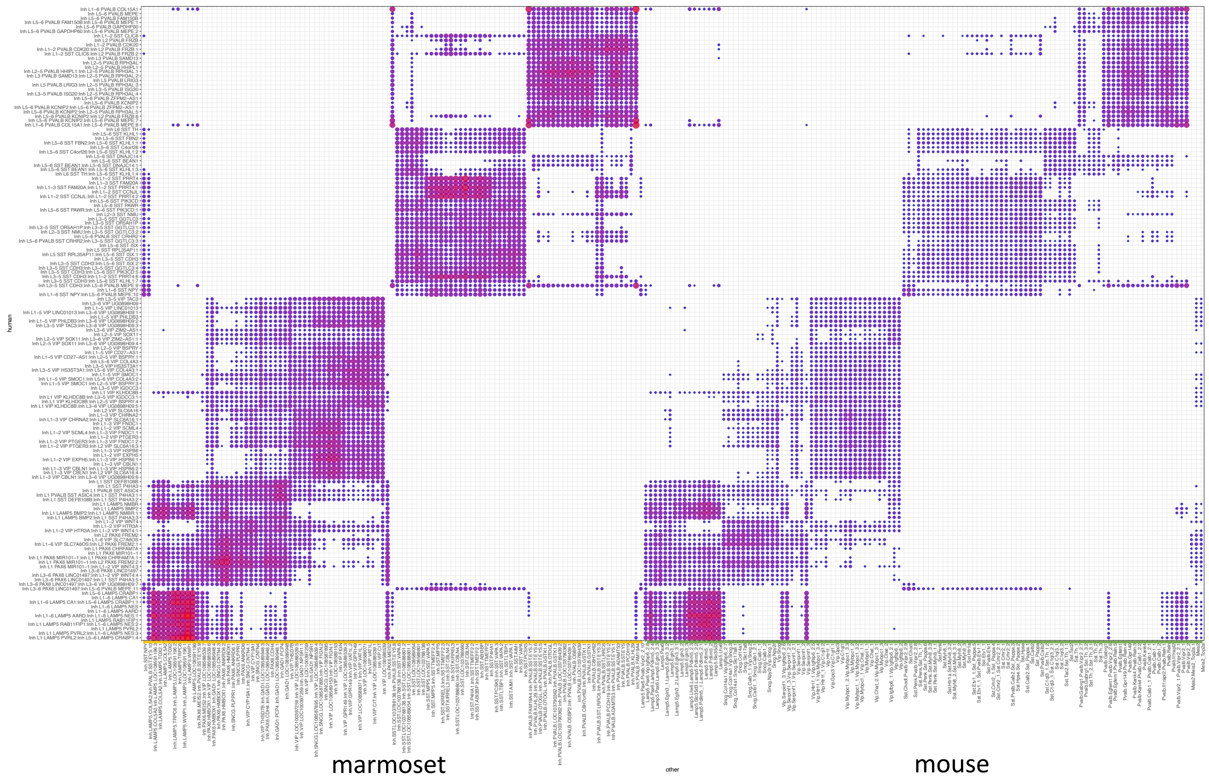


(a) Z-score

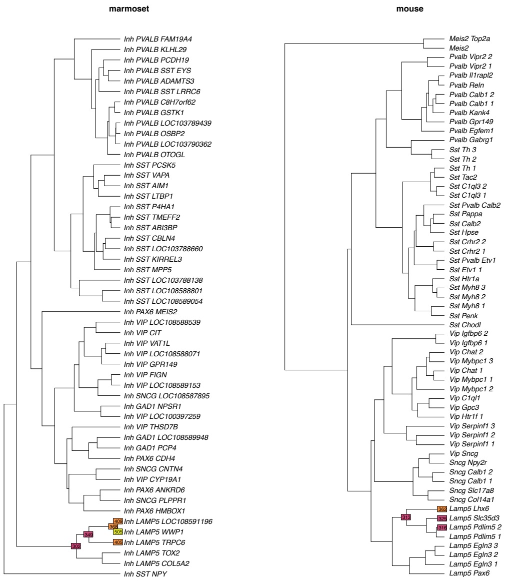


(b) LAMP5 AARD

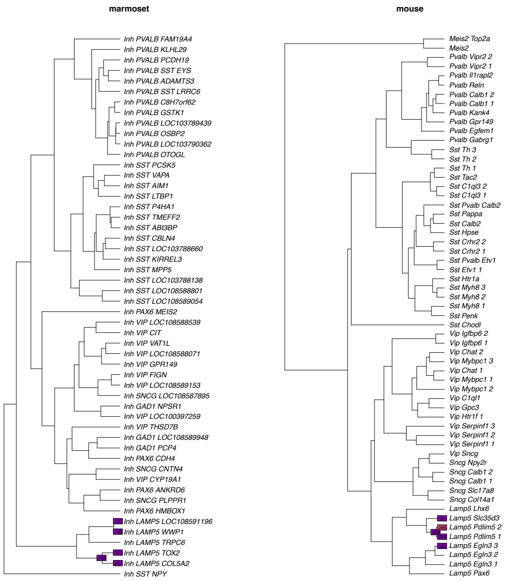


(c) LAMP5 NMBR

Figure S27: Mapping human cell types to marmoset or mouse cell types by CellWalker2's Z-score. (a) The mapping between human and marmoset cell type trees (including ancestor nodes). Each row is a human cell type, and each column is a cell type from marmoset or mouse. Only Z-scores > 3 are shown. (b-c) Top cell types on marmoset or mouse cell tree that are mapped to human LAMP5 AARD (b) or LAMP5 NMBR (c). Top 10 nodes with largest scores are shown. The color of the node represents the magnitude of the Z-score and the number on each mapped nodes is the value of the Z-score.

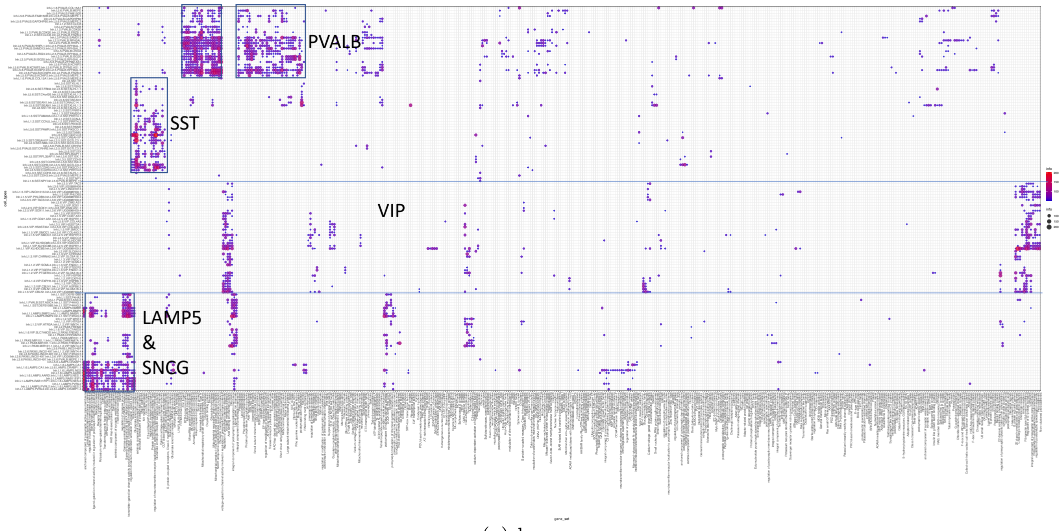


(a) human

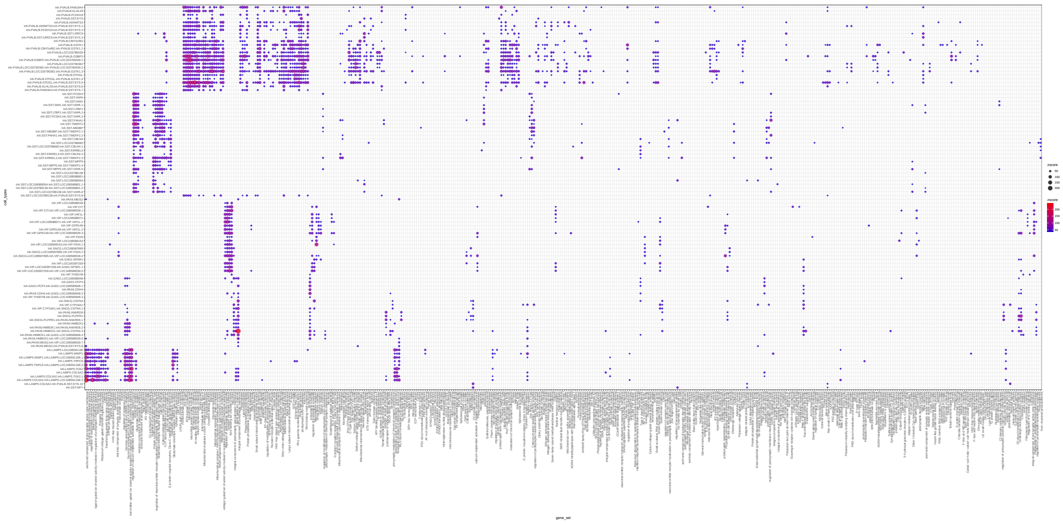

(b) marmoset

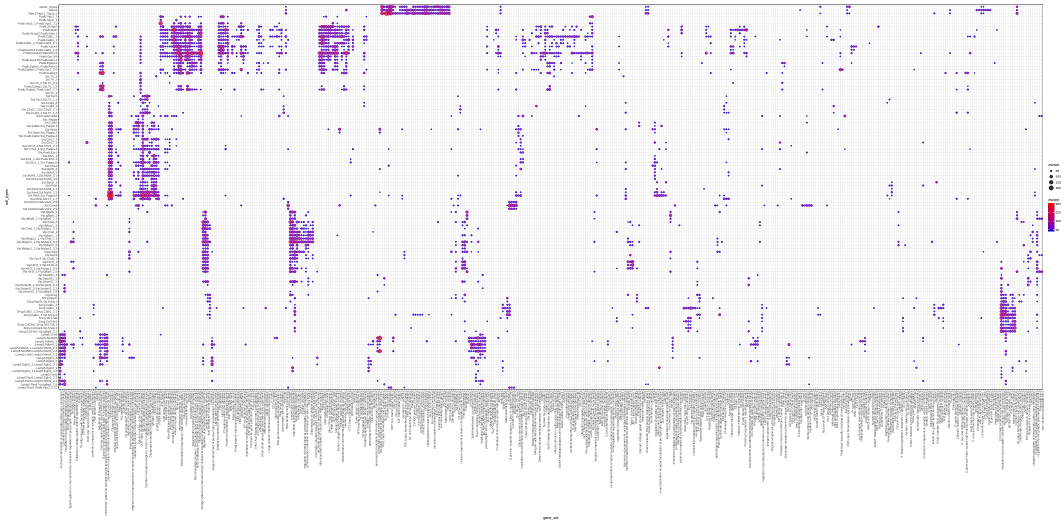

(c) mouse

Figure S28: Mapping active gene sets to cell types for different species. The heatmap shows the Z-scores from CellWalker2, where each row is a cell type (including ancestral nodes), and each column is a gene set. The color and size of the dot represents Z-score. TALE class homeobox transcription factors, which include Meis2, PBAF/BAF complex, SRY-box, and E2F, map to Meis2 and Top2a cell types that belong to the Meis2 subclass that is absent from the human ontology. Only top 5% of the entries are shown.

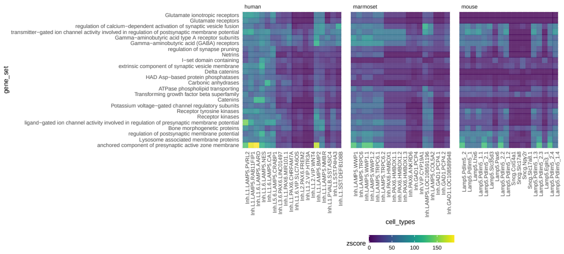

(d) gene sets active in human LAMP5 and SNCG cells

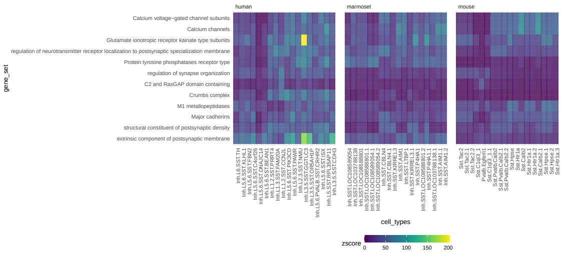

(e) gene sets active in SST cells

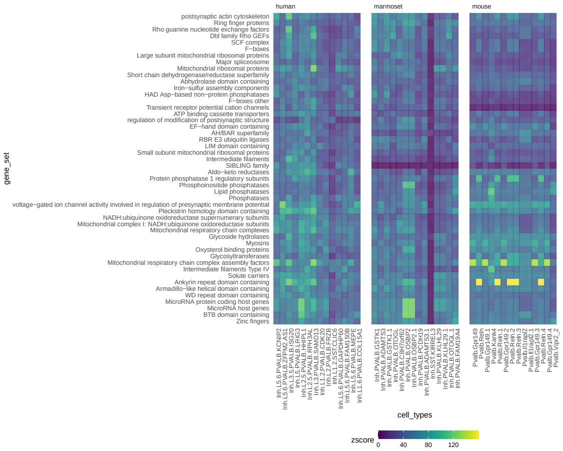

(f) gene sets active in human PVALB cells

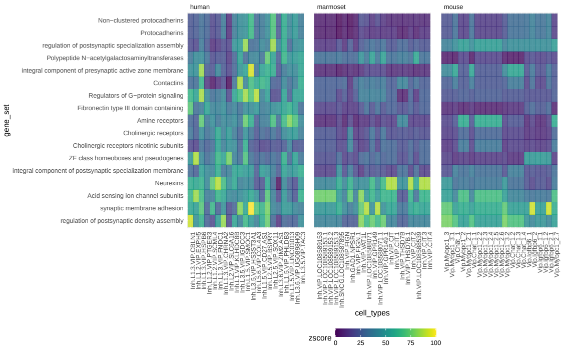

(g) gene sets active in human VIP cells

Figure S28: Gene sets that are activated in different subclasses of human cells show different cell type specificity across species. The three panels of each heatmap show the Z-scores between gene sets and cell types in human, marmoset and mouse. In each panel, each row is a gene set, each column is a cell type and the color represents Z-score. For each human cell type, the matched cell type by CellWalker2 in marmoset or mouse is shown.

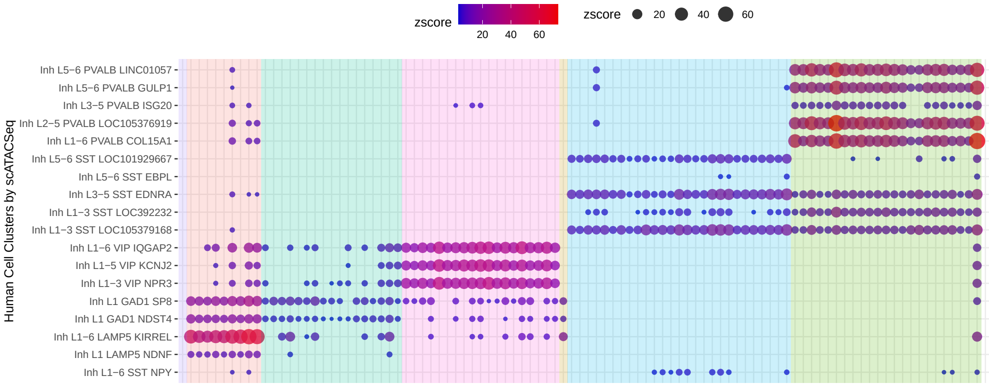


Figure S29: Mapping between human cell type-specifc DARs with marmoset cell types of inhibitory neurons using Z-score. Each row is a human cell type cluster identified by scATAC-Seq data and each column is a node on the marmoset cell type tree. Only Z-scores > 3 are shown. The color and size of the dot represent the Z-score. The background color indicates the cell subclass of marmoset.

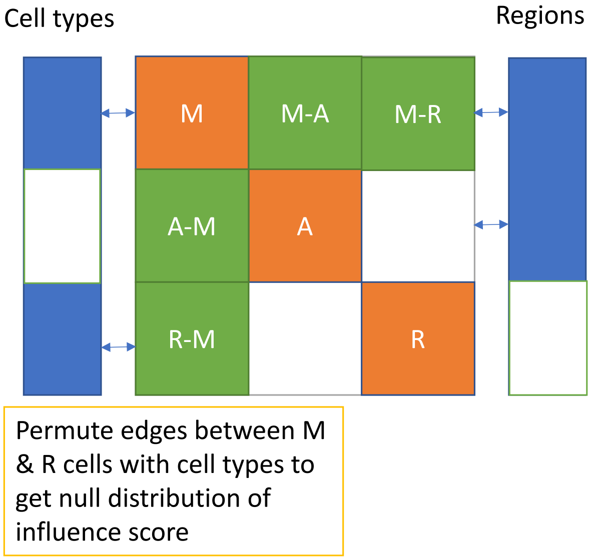


Figure S30: Illustration of integrating multiome data with scRNA-Seq and scATAC-Seq using CellWalker2. The left column represents cell types, the middle matrix is the cell-to-cell similarity matrix, and the right column represents annotations (i.e., different sets of genomic coordinates). A is the submatrix between cells with ATAC-Seq data only, whose distance is the Jaccard or Cosine distance computed from open chromatin peaks. R is the submatrix between cells with RNA-Seq data only, in which scaled Euclidean distance of gene expression in PCA space is used. M is the submatrix between cells with multiome data, whose distance is a weighted combination of RNA-Seq and ATAC-Seq distance. M-A and A-M are the cell similarity between cells with multiome and with scATAC-Seq data, in which Jaccard or Cosine similarity is computed from chromatin peaks. M-R and R-M are the cell similarity between cells with multiome and cells with scRNA-Seq data by computing Euclidean distance of gene expression in PCA space. M & R cells are connected to cell types by expression of markers. M & A cells are connected to regulatory regions by reads in each region. Blank areas have no edges. CellWalker2 computes a KNN graph based on the similarity matrix and uses shared KNN as edge weight. To estimate Z-scores for mapping from annotations to cell types, CellWalker2 permutes the edges between M & R cells and cell type labels to get the null distribution of the influence score.

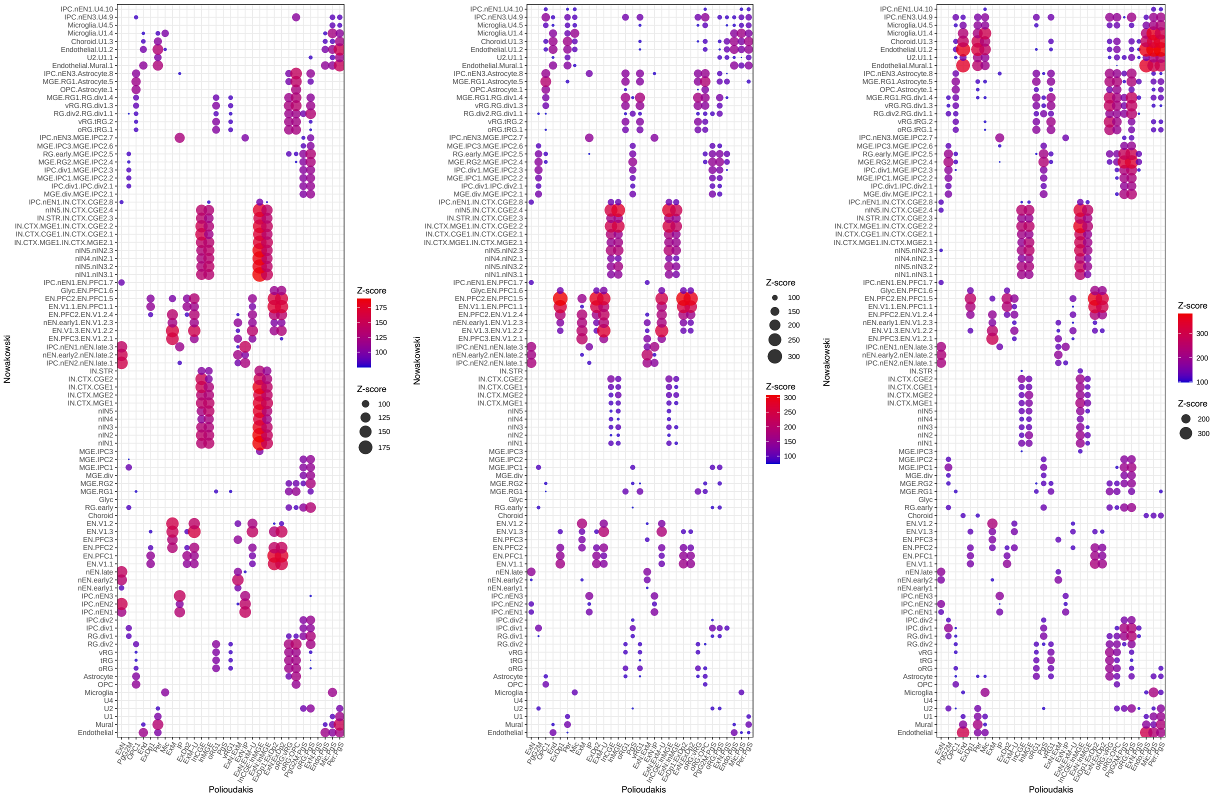


Figure S31: Different permutation schema for computing Z-scores for mapping cell types between the Polioudakis et al. and Nowakowski et al. datasets. We permute edge weights between cells and cell type ontology of Nowakowski et al. (left), between cells and cell ontology of Polioudakis et al. (middle), or between cells and both cell type ontology (right).

### Supplemental Note S1

**Varying similarity in the convergent cell type simulation scenario** This simulation illustrates a new cell state in a second dataset that combines features from different cell lineages in the first dataset. This could be the result of a disease or multipotent state. In the first dataset, we still have cell types 0,1,2,3 but the second dataset contains cell types 0,1,2,4, where cell type 4 has both features from cell type 1 and 3. In this case, the cell types in the second dataset do not follow a tree structure strictly. We simulated 4000 cells with 800 cells in each cell type, 900 non-marker genes, 300 marker genes which are differentially expressed between 0 and {1,4} but not 2 and 3, and 300 marker genes between 2 and {3,4}. In addition, we varied the distance of cell type 4 to cell type {0,1} by changing the number of marker genes between cell type {0,1} and {2,3,4} and between {0,1,4} and {2,3}. We simulated three scenarios where cell type 4 is getting closer to cell type {2,3} than cell type {0,1}: 1) high similarity, 120 marker genes between cell type {0,1} and {2,3,4}, 80 marker genes between {0,1,4} and {2,3}; 2) medium similarity, 150 marker genes between cell type {0,1} and {2,3,4}, 50 marker genes between {0,1,4} and {2,3}; 3) low similarity, 200 marker genes between cell type {0,1} and {2,3,4} only. Then, we split the cells into two equal sized datasets and added batch effects and dropouts.

From UMAPs, it is hard to tell the distance between cell type 4 and others (Figure S7a to S7c), but both influence score and Z-score between cell type 4 ('3_G') and cell type 3 ('3_U') decreases as expected (Figure S7d to S7i), and the influence score between cell type 4 ('3_G') and cell type 3 ('1_U') increases at the same time (Figure S7g to S7i). This simulation shows that CellWalker2 can provide a probabilistic mapping between cell type labels, even when some of the cell types do not follow a tree structure. Furthermore, influence scores and Z-scores from CellWalker2 can reveal the subtle change of cell states, which could be used to compare cell state across different conditions.

### Supplemental Note S2

**More details in cell type mapping of human developing cortex datasets** CellWalker2 maps (highest Z-score) migrating excitatory neurons (ExN) to the ancestor of late newborn excitatory neurons (nEN-late) and EN-like intermediate progenitors (IPC-nEN2) (Figure 5A), suggesting that excitatory neurons are more broadly defined in Polioudakis et al. than in Nowakowski et al.. Supporting CellWalker2's mapping, ExN is close to both cell types in UMAP space (Figure S18), and cell-to-cell distances from ExN to other cell types are correlated with Z-scores (Figure 5B). treeArches, on the other hand, maps ExN to the root node. It assigns relatively high probability to nEN-late and early newborn excitatory neurons (nEN-early2), but no probability to the third cell type in this clade (IPC-nEN2), most likely because it is comprised of fewer cells (4% vs >20%) and treeArches is sensitive to compositional bias. If we manipulate the cell type resolution of the Nowakowski et al. ontology, for instance by amalgamating excitatory neuron cell types, treeArches assigns larger scores to the combined cell type and reduces its score at the root node (Figure S20b). Alternatively, if we remove cell types with the highest scores for ExN, treeArches' scores increase for other cell types, including maturing excitatory neurons (EN-V1-2) that are from a different developmental stage but are very prevalent (Figures S20c and S20e). On the other hand, CellWalker2's top cell types stay the same with only minor decreases in Z-scores. These findings indicate that treeArches may be better able to map cell types when the two ontologies have comparable resolution, while CellWalker2's use of internal nodes enables mappings between fine-resolution and broad cell types.

Inhibitory neurons provide another example of closely related cell types. The Nowakowski et al. ontology contains many types of inhibitory neurons from different locations and developmental stages. CellWalker2 maps inhibitory neurons of the medial ganglionic eminence (InMGE) from Polioudakis et al. to one type of MGE inhibitory neurons (IN-CTX-MGE1), whereas those from the caudal ganglionic eminence (InCGE) map to the ancestor of two types of CGE inhibitory neurons (Figure 5A), reflecting uncertainty about either 1:1 mapping. With treeArches, the root node has a high probability for both InMGE (56%) and InCGE (33%). This observation, along with the ExN results described above, suggests that the inclusion of closely related cell types in a dataset may lower treeArches' mapping scores to each of them but encourages CellWalker2's mapping to all these cell types and their ancestor.

CellWalker2 facilitates the interpretation of cell types by mapping labels in another ontology that carry additional information about cell state. There are two groups of RG-like Intermediate progenitor cells in Nowakowski et al. with IPC-div1 having the highest Z-score for cycling progenitor G2/M phase (PgG2M) and IPC-div2 for S phase cycling progenitors (PgS), while the parent node of IPC-div1 and IPC-div2 has roughly equal Z-scores for PgG2M and PgS (Figure 5C).

### Supplemental Note S3

**Vary cell type composition in cell type mapping** To show how cell type composition would affect the results from CellWalker2 and treeArches, we removed or combined cell types from a group of newborn excitatory neurons and intermediate progenitors that was where ExN mostly mapped. In general, the ranking of Z-scores is robust to either removing or combining related cell types, but treeArches' results can vary depending on which cell types are included in the dataset (Figure S20). If we combined all the cell types in the subtree to which ExN is mapped into one cell type (labeled as 'nEN'), treeArches assigned larger scores to the combined cell type and smaller scores to the root compared to including separate cell types (Figure S20b). This indicates that treeArches might be more capable of mapping cell types on at a similar resolution rather than mapping a more broadly defined cell type to several subtypes. If we removed the entire subtree from the cell type hierarchy, treeArches showed increased scores to other related cell types, such as early born newborn excitatory neuron (nEN-early1) and early and late born excitatory Neuron V1 (EN-V1-2) (Figure S20c). This is consistent with the simulation of a 'convergent cell type', in which treeArches does not recover the similarity to the cell type from a different subtree and assigns large probability to the root. On the other hand, the Z-score mapping to 'nEN-early1' is similar or smaller when the subtree is excluded. If we removed Intermediate Progenitor Cells EN-like (IPC-nEN2) and late born Newborn Excitatory Neuron (nEN-late) cells as well as these cell type labels on the tree, which are the top cell types that ExN was mapped to by CellWalker2 (Figure S20a), CellWalker2 showed similar or smaller Z-scores to the remaining cell types; while treeArches assigned larger scores to early born newborn excitatory neuron (nEN-early2) (Figure S20d). Lastly, if we removed 'nEN-early2' and 'nEN-late' cells as well as these cell type labels on the tree, which were the top cell types that ExN was mapped to by treeArches (Figure S20a), treeArches assigned larger scores to 'IPC-nEN2' and 'nEN-early1' as well as the root node (or being rejected), while CellWalker2 shows similar Z-scores mapping to the remaining cell types (Figure S20e). The robustness of CellWalker2 to cell type composition might be due to using marker genes as well as the cells from each dataset. Furthermore, the marker genes of the remaining cell type labels stay the same even when cell type composition is changed.

**Vary the number of markers or add noise in cell type mapping** We varied the number of markers per cell type or added noise to the gene expression for constructing cell-to-cell graphs to see how CellWalker2's results on mapping cell types change. We ordered the cell type marker by log fold change and varied the number of markers for each cell type from top 20%, 50%, 80% and 200% of the original markers. We reran CellWalker2 with different marker sets and computed the proportion of significant mappings overlapping with CellWalker2's original results. Specifically, suppose A and B are lists of entries above certain significance level under different experiment conditions, we computed $|A \cap B|/|A \cup B|$ as the "percentage of overlaps". Figure S22a shows the results with different p-value cutoffs, where p-values are converted from Z-scores using the standard Normal distribution. With fewer marker genes, the percentage of overlaps decreases. However, there are still above 80% overlap even with only 20% of markers. On the contrary, with twice the number of markers, the percentage of overlaps is above 95% at various significance levels. Next, we added Gaussian noise with different standard deviations to each of the top principal component coordinates derived from the gene expression matrix. We recomputed the cell-cell similarity matrix based on the noisy coordinates and reconstructed the cell-to-cell graph. Figure S22a shows the percentage of significant entries overlapping with the results without noise at different p-value cutoffs and different noise levels. The noise level is the ratio of the standard deviation of noise and average of each coordinate. As the noise level increases, the percentage overlapping decreases. At noise level 0.78, which means that adding noise about 80% to the overall variability of the data, the percentage of overlaps is still above 85%. Interestingly, the percentage overlapping increases as we increase the noise level above the overall variation of the data (data not shown). We speculated that as cell-cell similarity is overwhelmed by noise, the label-to-cell information flow dominates the noisy cell-to-cell graph during CellWalker2's random walks. Label-to-cell edges can still guide towards the correct cell type mappings.

#### Supplemental Note S4

**Cell subclass-specific transcription factors in human and marmoset** Top TFs shared between human and marmoset within similar cell subclasses include *TCF4* and *ZEB1* in Vip cells, *NFIB* and *NFIX* in all CGE-derived cell types, *NFIA* in Lamp5 cells, *SP8* and *SP4* in Sncg and Lamp5 cells and *RORA*, *MEF2C*, *MEF2A*, *BACH1*, *ESRRA*, *ESRRG* in Pvalb cells. Cell type-specific TFs unique to marmoset include *THAP2A*, *SOX10*, and *EGR3* in Vip and Sncg cells, *KLF11*, *POU3F2*, and *RREB1* in Lamp5 cells, and *RARB*, *MAFB*, *JDP2* and *BACH2* in Pvalb and Sst cells (Figure 6E).

Then, we compared CellWalker2 results with ArchR running on the same data. We used CellWalker2's direct output, without filtering for expression, because ArchR does not do so. CellWalker2 identifies more cell subclass-specific TFs compared to ArchR and also some TFs shared among similar cell subclasses (Figure S32). Z-scores separate TFs into two major groups, one corresponding to CGE lineages (Vip, Sncg, and Lamp5 subclasses plus their ancestral nodes) and the other to MGE lineages (Pvalb and Sst subclasses plus their ancestors). Moreover, while CellWalker2 identified several TFs unique to Pvalb and Vip cells, which originate from distinct lineages (MGE- versus CGE- derived), while ArchR mostly identified TFs that are shared between Pvalb and Vip cells. In addition, CellWalker2 identified a group of TFs having higher Z-scores in both Sst chodl and Sst cells, probably due to the similarity between Sst chodl and Sst cells and the small sample size of Sst chodl cells.

### Supplemental Note S5

**More discussion about permutation schema** Under the permutation null distribution in the current CellWalker2, the edges from a cell are randomly assigned rather than preferentially assigned to related cell types. This is a strong null distribution that accurately estimates the null value of the mean influence score, while potentially underestimating the standard deviation of the influence score, because permutations do not preserve correlations between cell types. We tried sub-sampling cells to estimate the standard deviation of influence scores for computing Z-scores, but the results were similar or worse than the current approach, as the cells are interconnected in the graph and hence not properly connected in sub-samples.

### Supplemental Note S6

**Mapping cell type annotation between multiome and scRNASeq data** As a sanity check, we compared cell type annotations in the multiome data to the cell type tree built from scRNA-Seq across developmental stages (excluding VLMC and RBC which have less than 5 cells in pcw 21) in Trevino et al.. Because of different resolutions and developmental stages of the two datasets, the clusters within glutamateric neurons do not have one-to-one correspondence with different groups of glutamateric neurons in the scRNA-Seq data, and some of the cells in the multiome data are mapped to internal nodes on the cell tree, even though the major cell types are matched (Figure S33c). The IN1 cluster from the multiome data is mapped to CGE IN in the scRNA-Seq ontology, while IN2 is mapped to both MGE IN and the the parent node of IN. The multiome RG cluster is mostly mapped to late RG, but also early RG. The EC/peric cluster is mapped to the parent node

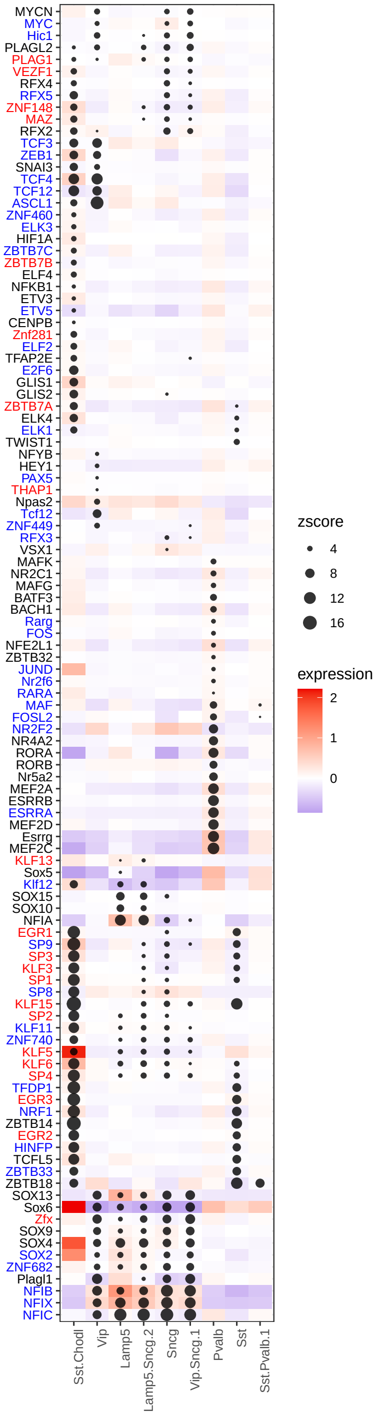


(a) CellWalk2, Human

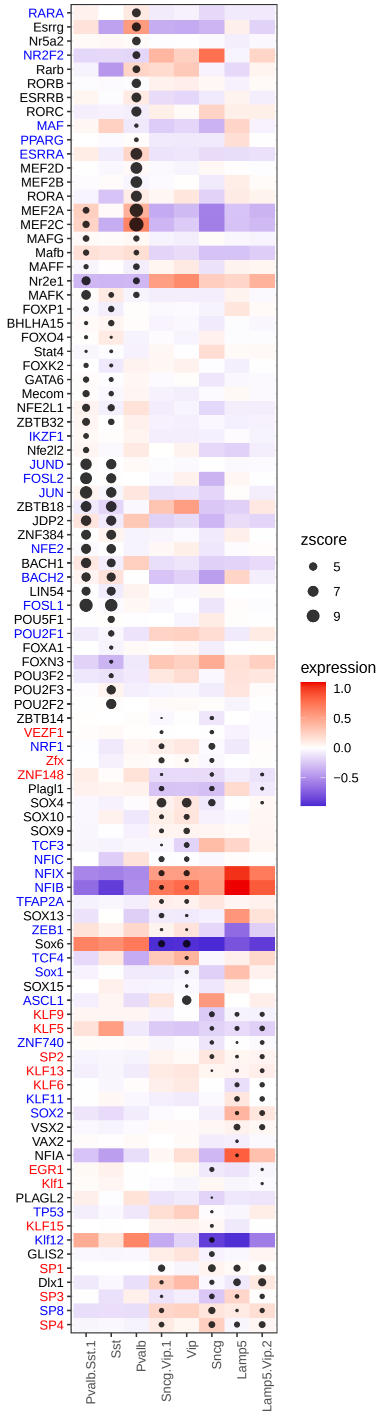


(b) CellWalk2, Marmoset

of Peric and EC. The mGPC/OPC cluster is mapped to the ancestor node of both mGPC and OPC/Oligo. nIPC/GluN1 is mapped to multiple glutamateric neurons types and nIPC.

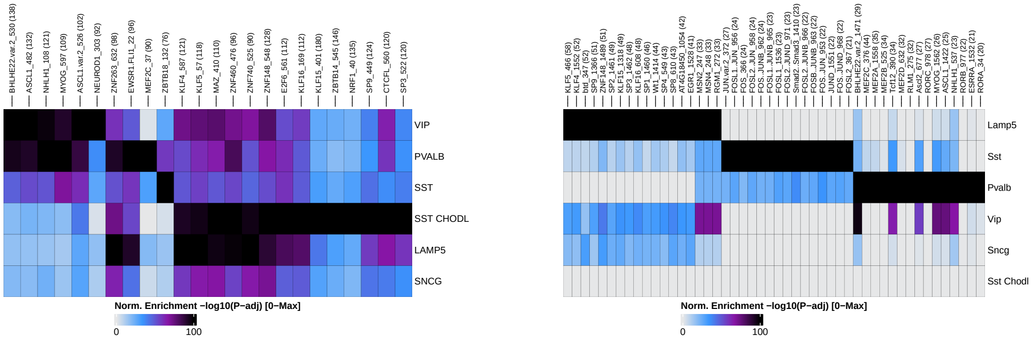


(c) archR, Human

(d) archR, Marmoset

Figure S32: Mapping between TFs and marmoset or human cell subclasses for inhibitory neurons using CellWalker2 and archR. For CellWalker2, each row is a transcription factor (TF), and each column is a node on the cell tree. Only Z-scores $> 3$ are shown. The size of the dot represents the Z-score, and the color represents the TF's standardized expression level. Sst chodl is excluded for Marmoset because it has too few cells. For archR, each row is a cell type, and each column is a TF. The color represents -log10 Pvalues.

The mapping between cell labels without considering tree structure is shown in Figure S33b.

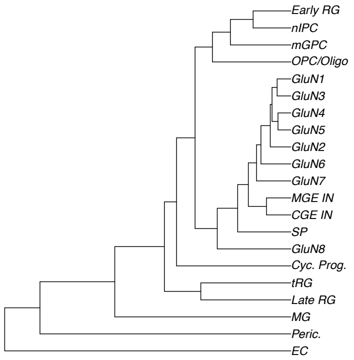


(a) Cell type tree from scRNASeq

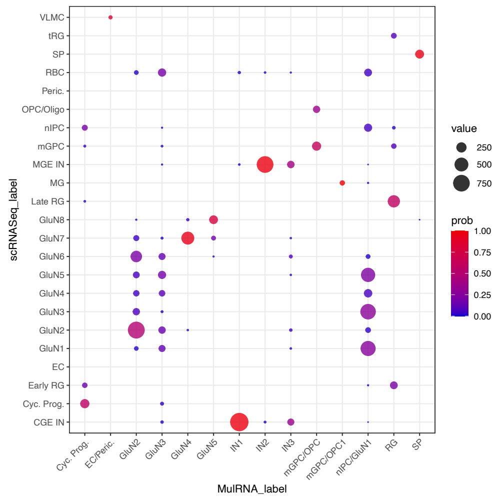


(b) tip cell types

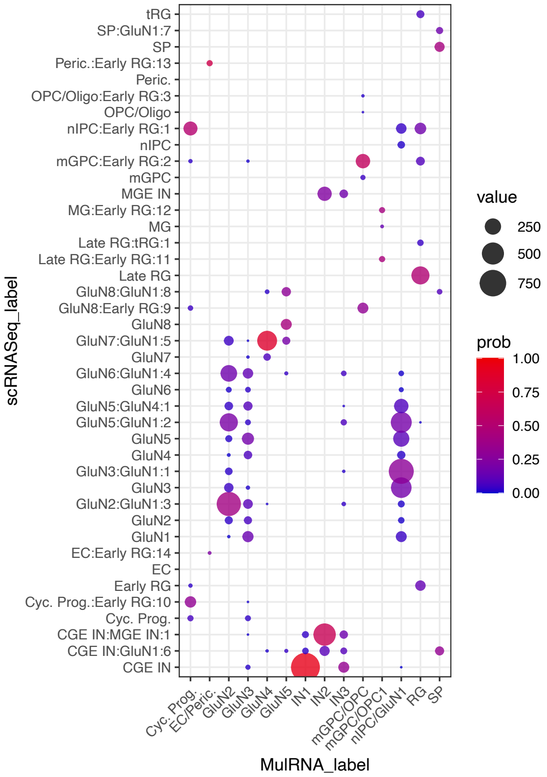


(c) all nodes on the cell type tree

Figure S33: Comparing cell type annotation in multiome data with scRNASeq using CellWalker2. The dotplot shows the assignment of cells of each cluster in multiome data by cell type hierarchy from scRNASeq data. The size and color of the circle represents the number and the proportion of cells, respectively, in each cluster in the multiome data assigned to each cell type label (b) or each node on the cell type hierarchy (c) in scRNA-Seq data. The cell type tree is shown in (a).
